# Single-cell multi-omics map of human foetal blood in Down’s Syndrome

**DOI:** 10.1101/2023.09.25.559431

**Authors:** Andrew R. Marderstein, Marco De Zuani, Haoliang Xue, Jon Bezney, Shuo Wong, Tim H. H. Coorens, Stephen B. Montgomery, Ana Cvejic

## Abstract

Down’s Syndrome (DS) predisposes individuals to haematological abnormalities, such as increased number of erythrocytes and leukaemia in a process that is initiated before birth and is not entirely understood. To understand dysregulated hematopoiesis in DS, we integrated single-cell transcriptomics of over 1.1 million cells with chromatin accessibility and spatial transcriptomics datasets using human foetal liver and bone marrow samples from three disomic and 15 trisomic foetuses. We found that differences in gene expression in DS were both cell type- and environment-dependent. Furthermore, we found multiple lines of evidence that DS haematopoietic stem cells (HSCs) are “primed” to differentiate. We subsequently established a DS-specific map of enhancer-gene relationships in disomic and trisomic HSCs using 10X Multiome data. By integrating this map with genetic variants associated with blood cell variation, we discovered that trisomy restructured enhancer-gene maps to dysregulate enhancer activity and gene expression critical to erythroid lineage differentiation. Further, as DS mutations display a signature of oxidative stress, we validated both increased mitochondrial mass and oxidative stress in DS, and observed that these mutations preferentially fell into regulatory regions of expressed genes in HSCs. Altogether, our single-cell, multi-omic resource provides a high-resolution molecular map of foetal haematopoiesis in Down’s Syndrome and indicates significant enhancer-gene restructuring giving rise to co-occurring haematological conditions.

## Introduction

Children with trisomy of chromosome 21 (Ts21), also known as Down’s syndrome (DS), have a 150-fold higher risk of developing myeloid leukaemia in the first five years of life compared to the general population^1,2^. The current leukaemogenesis model in DS proposes cell-intrinsic events that start in Ts21 foetal liver via specific somatic mutations in the transcription factor GATA1, leading to preleukaemia development^3^. Preleukaemia spontaneously resolves in most newborns, but it can progress to leukaemia when persistent GATA1 mutant clones acquire additional mutations post-birth, which are often in chromatin regulators and signalling factors^4,5^. The proposed model, while insightful, does not explain further numerous haematological abnormalities present in Ts21 foetal liver and newborns^4^. Neonates with DS are often born with defects in red blood cell production, such as polycythaemia (increased number of erythrocytes) or macrocytosis (large erythrocytes)^6^, in a process that is not fully understood but independent of any leukaemia-inducing mutations. These haematological abnormalities are highly variable in both severity and penetrance across individuals, suggesting that factors such as epigenetics, the composition of the microenvironment, and gene-environment interactions may also contribute to the disease progression^7–9^. For example, during foetal development, haematopoiesis occurs simultaneously in both the foetal liver and bone marrow (BM). Previous research in DS has shown an increase in megakaryocyte-erythroid progenitors in the foetal liver but not in the BM, along with a reduction in granulocyte-monocyte progenitors, emphasising the microenvironment’s relevance.

Notably, Ts21 foetal liver haematopoietic stem and progenitor cells (HSPCs) not only display an elevated mutational rate but also exhibit a unique mutational signature related to oxidative stress^10^. Oxidative stress is a potent driver of mutagenesis and is considered a potential cause for the GATA1 mutations observed in Ts21 preleukemia and acute megakaryocytic leukaemia^10,11^. However, given the well-know and substantial impacts of chromatin organisation on mutation patterns^12,13^, and Ts21’s effect on reconfiguring the entire 3D genome^14^, it remains to be evaluated whether Ts21-induced genome-wide reshaping of chromatin organisation in a cell-type-specific manner underlies distinct haematological phenotypes observed in Ts21, such as an increase in erythroid cells but an impairment in the lymphoid lineage. Further, it remains to be determined how altered chromatin organisation in Ts21 in combination with heightened oxidative stress exposure could potentially create an environment conducive to acquiring mutations that contribute to the development of preleukaemia and leukaemia.

Here we combined single-cell RNA-Seq (scRNA-Seq) from 1.1 million cells, 10X multiome, spatial transcriptomics and single-cell *in vitro* differentiation assays to examine how foetal haematopoietic cells with an altered dosage of chromosome 21 genes interact with particular niches (foetal liver versus BM) and how these interactions differ from those in disomic foetuses. We further assessed the regulatory landscape of haematopoietic cells in Down’s Syndrome and its impact on gene expression, cellular differentiation, and the regional distribution of somatic mutations.

## Results

### Multiomics analysis of human disomic and Down’s syndrome foetal liver and bone marrow

The foetal liver and the BM are complex cellular ecosystems made up of many haematopoietic and non-haematopoietic (niche) cell types (such as endothelial, stellate, osteo-lineage, and other niche populations). In order to capture a full diversity of blood and niche cell types present in the foetal liver and BM (isolated from the femur), we implemented a variety of sorting and column-enrichment strategies. We isolated cells from 15 Ts21 and three disomic foetuses (median age 14 post-conception weeks (pcw)), from each of the following three populations: CD235- (niche and haematopoietic cells depleted of erythrocytes), CD34+/Lin- (haematopoietic progenitors), and CD45+ (pan-haematopoietic marker), and sequenced their transcriptomes using 10x Genomics (Figure 1a, Supp. Figure 1, 2). In addition, we collected part of the liver from 11 Ts21 and five disomic foetuses for spatial transcriptomics (STx) analysis using 10X Genomics Visium. Finally, for three Ts21 and three disomic liver samples, we performed 10X Multiome analysis of CD45+ cells, combining a single-cell assay for transposase-accessible chromatin sequencing (scATAC-seq) and snRNA-seq from the same cells (Figure 1a).

**Figure 1.**
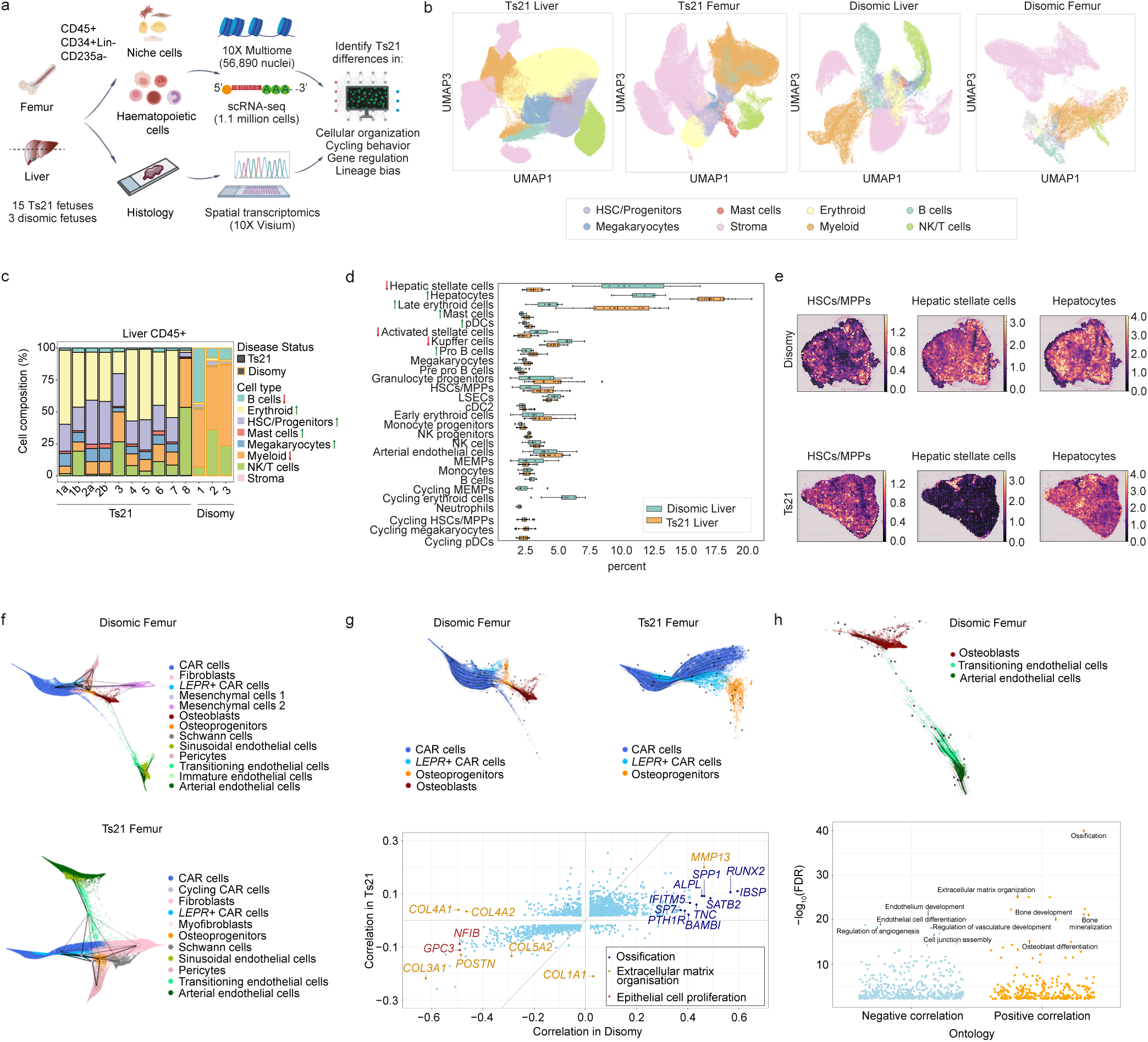
Single-cell transcriptome analysis of human Down’s syndrome (Ts21) foetal liver and bone marrow. **a**, Schematic overview of the experimental workflow. Cells (CD45+, CD34+lin- and CD235a-) from disomic and Ts21 foetuses (median age 14 and 13 PCW, respectively), were isolated from BM (femur) and liver and processed for scRNA-Seq. In addition, CD45+ cells from foetal liver (k=3) were processed for Multiome and tissue sections from disomic and Ts21 foetuses were used for spatial transcriptomics. **b**, 3D Uniform manifold approximation and projection (UMAP) visualisation of cells from Ts21 liver (n=780,299, k=15), Ts21 femur (n=162,775, k=12), disomic liver (n=110,671, k=3) and disomic femur (n=53,807, k=3) coloured by broad cell type categories. n refers to the total number of cells, k indicates the number of biologically independent samples (foetuses). **c**, Stacked bar plot of relative abundances of different broad cell types in stage-matched Ts21 (black outline) and disomic foetuses (orange outline). Arrows at the side of the labels indicate a significant increase (↑) or decrease (↓) in the number of cells in Ts21 liver, compared to disomic liver. Difference in cell proportion was tested by Wilcoxon Rank Sum test followed by Bonferroni correction. **d**, Dodged boxplot of relative spatial abundances of different cell types in Ts21 and disomic liver, as estimated by cell2location. Arrows at the side of the labels indicate a significant increase (↑) or decrease (↓) in the number of cells in Ts21 compared to disomic liver. Difference in cell proportion was tested by Wilcoxon Rank Sum test followed by Benjamini-Hochberg correction. **e,** cell2location mapping of haematopoietic stem cells/multipotent progenitors (HSCs/MPPs), hepatic stellate cells, and hepatocytes on two representative Visium sections from Ts21 foetal liver (bottom) and disomic liver (top). The colour of each spot indicates the cell type abundance estimated by cell2location (mean value). **f,** PAGA graphs overlaid on the diffusion maps, computed from the 11 (Ts21) and 13 (disomic) cell types representing the BM niche population and their connectivity (top-disomic, bottom-Ts21). **g,** Top: trajectory with the directionality of the differentiation process from CAR cells to osteoprogenitors and osteoblasts in disomic and Ts21 samples. Stream plot overlaid on Force-directed (FLE) graph of diffusion map of CAR cells, *LepR*+ CAR cells, osteoprogenitors and osteoblasts (the latter only being detected in disomic BM). Bottom: scatter plot of the Pearson correlation between gene expression and CellRank’s terminal state absorption probabilities in disomic and Ts21. Only significantly correlated genes (FDR < 0.05) were plotted. Genes were coloured based on the GO terms they belong to (blue: “ossification”; orange: “extracellular matrix organisation”; red: “epithelial cell proliferation”). **h.** Top - stream plot overlaid on FLE graph of diffusion map of arterial endothelial cells, transitioning endothelial cells and osteoblasts. Bottom - scatter plot of GO terms estimated from top 500 genes most positively/negatively correlated with the absorption probabilities.

### Cellular and spatial organisation of Ts21 and disomic foetal samples

Following quality control (QC) (see Methods), our scRNA-seq dataset comprised 943,074 cells from Ts21 and 164,478 cells from disomic foetuses (Supp. Table 1 and 2). We identified eight major blood and niche cell types in both the Ts21 and disomic foetal liver and femur (Figure 1b, Supp. Figure 2) which were further resolved into 37 cell types in femur and 25 in liver (see Methods, Supp. Results, Supp. Figure 1c-f, 2, 3, & 4, Supp. Table 3-7). Overall, the frequency of most cell types was similar between Ts21 and disomic liver samples, however, there were notable differences. In Ts21, we detected an increase in HSC/Multipotent Progenitors (MPPs) (as previously described^15^), erythroid cells, mast cells and megakaryocytes (FDR < 0.1) (Figure 1c, Supp. Figure 5, Supp. Table 8). Furthermore, there was an additional population of highly cycling HSCs in Ts21 liver (Supp. Figure 3). In line with previously reported impairment in B cell development^15^, we did not detect a distinct cluster of mature B cells in the Ts21 foetal liver (Supp. Figure 3 and 4). In contrast, the proportion of myeloid cells was decreased in Ts21 liver compared to disomic liver (FDR < 0.1 for each population) (Figure 1c, Supp. Figure 5, Supp. Table 8). Importantly, we experimentally validated that the Ts21 causes wide perturbation in the stem and progenitor compartment in the foetal liver using flow cytometry (Supp. Figure 6), which underscores the robustness of our computational classification of cell types in scRNA-seq. Haematopoietic differentiation appeared to be less perturbed in Ts21 samples within the BM compared to the liver, as the population of cycling HSCs and dysregulated progression of B cell maturation was mainly detected in the latter (Supp. Figure 3 *versus* Supp. Figure 4). We subsequently confirmed the robustness of our cell-type annotations and differences in frequency of cell types between Ts21 and disomic liver samples after integration using Harmony^16^ and scVI^17^, as well as reference-query mapping from a published disomic foetal liver atlas^18^ and our Ts21 liver cells to disomic liver cells (Supplementary Results; Supp. Figure 7-9).

To validate differences in cell abundances and examine how scRNA-seq-identified cell types are spatially arranged in DS, we performed spatial transcriptomics. We performed 10X Visium on two consecutive 10 µm sections from 11 Ts21 foetal livers (all matched to the original samples used for the scRNA-seq) and five disomic foetal livers (Supp. Table 9). We estimated the abundance of cell types within each Visium spot by leveraging cell-type expression profiles from the scRNA-seq^19^. Once cell types were resolved on tissue, we examined the frequency of different cell types across all sections in Ts21 *versus* disomic samples (Figure 1d). As in the scRNA-seq dataset, the number of HSCs was higher (although not statistically significant) in Ts21 compared to disomic liver. Furthermore, hepatic stellate cells (FDR=2.3E-07), activated stellate cells (FDR=0.003), and Kupffer cells (FDR=0.01) were less abundant in Ts21 foetal liver compared to the disomic tissue, whereas mast cells (FDR=0.002), late erythroid cells (FDR=1.8E-06), pDC (FDR=0.002), pro B cells (FDR=0.03), and hepatocytes (FDR=2.3E-06) showed higher frequency in Ts21 (Figure 1d, e). The high concordance in the results obtained in scRNA-Seq and Visium suggests that our spatial “map” of different cell types accurately represents their distribution in the tissue and revealed significant alterations in myeloid and lymphoid lineage abundances.

To investigate whether observed differences in cell frequency affected the spatial organisation of cells in Ts21 *versus* disomic liver, we quantified the colocalization of cell types residing in the same Visium spot (see Methods). Our analysis showed that haematopoietic populations with close lineage relationships tended to co-localise in the same spot in disomic foetal liver (Supp. Figure 10). For example, relative abundance of HSC/MPPs was highly correlated with megakaryocyte-erythroid-mast progenitors (MEMPs), granulocyte progenitors, pre-pro B cells and early erythroid cells; monocyte progenitors were correlated with monocytes and NK progenitors with NK cells (Supp. Figure 10a). In addition, Kupffer cells, hepatic stellate cells and liver sinusoidal endothelial cells (LSECs), known to reside in close vicinity in foetal liver^20^, were also highly correlated in our Visium data (Supp. Figure 10). LSECs act as gatekeepers that prevent hepatic stellate cell activation and in disomic liver activated stellate cells were often co-localising with arterial endothelial cells (Supp. Figure 10a-b). In Ts21 however, several close interactions were perturbed; hepatic stellate cells were more correlated with arterial endothelial cells and pre-pro B cells lost their connection with HSCs (Supp. Figure 10a-b). This indicates that both the cell abundance and co-localisation of cell types in Ts21 foetal liver are markedly distinct from that in disomic samples.

We next calculated, based on the co-expression of genes in different cell types clusters^21^, which ligand-receptor pairs (L-Rs) are potentially involved in HSC/MPP cell interactions in Ts21 and the disomic foetal liver (see Methods). HSC/MPPs shared the most L-Rs with arterial endothelial cells and the least with hepatocytes (Supp. Figure 10c). We identified a number of well-known L-Rs involved in HSC/MPPs development and function which were present in both Ts21 and disomic samples, such as VEGFB-FLT1^22^, ANGPT1-TEK^23^, and NOTCH1- JAG1^24^(Supp Figure 10d). Gene-set enrichment analysis of L-Rs expressed in HSC/MPPs (used for their interaction with arterial endothelial cells) revealed engagement of ERK1/2 cascade (Supp Figure 10e) which is a common target of growth factors regulating haematopoiesis^25^. Activation of ERK1/2 pathways promotes HSC proliferation, a hallmark of foetal haematopoiesis, at the expense of quiescence. The interaction of HSCs with hepatocytes (both in Ts21 and disomic) included L-Rs relevant for iron ion uptake such as TFRC-TF (Supp Figure 10d).

### Osteo-lineage development is altered in Ts21 bone marrow

Osteo-lineage is an important regulator of the HSC’s activity in the BM. Therefore, we examined if Ts21 affects differentiation and gene expression of osteoblasts in BM (Supp. Figure 4, Supp. Table 5-6, Supp. Results for annotations). Interestingly, the osteo-lineage appeared perturbed in Ts21 with many of the key osteo-lineage genes (such as *BGLAP, IBSP*, or *SP7*) being sparsely expressed (Supp. Figure 4). To examine this further, we generated a differentiation trajectory within the BM niche and calculated dynamically expressed genes along the osteo-lineage trajectory^26–28^ (from CXCL12-abundant reticular (CAR) cells to leptin receptor positive CAR (*LepR*+ CAR) cells to osteoprogenitors to osteoblasts, the latter being present only in disomic BM) (see Methods; Figure 1f,g; Supp. Figure 11a). The 25 genes whose expression had the strongest positive correlation with osteoblast differentiation in disomic samples included *IBSP*, *RUNX2*, *SP7*, and *SPP1* (Figure 1g), which are all related to “ossification”. In contrast, genes related with “extracellular matrix organisation” and “epithelial cell proliferation” were negatively correlated. In Ts21, however, there was a weaker correlation between terminal state probability (a quantitative measure of osteo-lineage differentiation) and gene expression (both positive and negative) (Figure 1g) suggesting a less coordinated regulation of osteo-lineage related genes (Supp. Figure 11b) that partially impeded osteo- lineage development.

We further observed that a population of endothelial cells (here termed transitioning endothelial cells) formed a continuum that connected clusters from endothelial cells with the osteo-lineage (Figure 1f, h; Supp. Figure 11c), suggesting that endothelial cells can go through endothelial-to-osteoblast transition in disomic foetuses. It has been recently reported that endothelial cells can give rise to BM stromal niche cells through endothelial to mesenchymal transition (EndoMT)^29^. Along the differentiation trajectory transitioning endothelial cells upregulated mesenchymal genes such as *PDGFRA*, *PDGFRB* and *PRRX1* (Supp. Figure 11d) as well as “ossification” related genes such as *IBSP*, *ALPL* and osteoblast cadherin *CDH11*, and downregulated “endothelium development” and “angiogenesis” genes such as *KDR* and *CDH5* (Figure 1h, Supp. Figure 4, Supp. Figure 11c). In Ts21, however, transitioning endothelial cells showed less prominent upregulation of mesenchymal genes pointing again towards a defect in osteo-lineage differentiation in Ts21 BM (Figure 1f, Supp. Figure 4, Supp. Figure 11d). Our data supports the notion that perturbed osteo-lineage differentiation in DS foetuses starts early during foetal development, and may underlie previous clinical observations such as shortened bone lengths and increased risk of osteoporosis^30–33^.

### Gene-environment interactions contribute to phenotypic differences between haematopoietic cells in the liver and BM

An increase in somatic mutation rate in Ts21 foetal HSPCs is not sufficient to explain the increased rate of leukaemia development in DS children. Therefore, it is likely that genetic background (trisomic versus disomic) and haematopoietic environment (liver versus femur), along with the cell–cell competition and their clonal selection, are significant factors^10^. To assess microenvironmental effects in DS, we examined gene expression across all cell types in foetal liver vs BM (see Methods). We observed around 50% increase in expression of chr21 genes across all cell types in Ts21 samples compared to control (Supplementary Results, Supp. Figure 12 a, b). Changes in non-chr21 gene expression appeared to be cell type-dependent but also influenced by the environment from which cells were isolated (liver or BM) (Supp. Figure 12 a, b). We further observed an exponential relationship in dysregulation of genes on chr21 *versus* non-chr21 genes (in contrast to an approximate linear relationship of chr1 versus non-chr1 genes), suggesting that a “critical mass” of around 30%-50% dysregulated genes on chr21 was necessary before widespread (>10%) dysregulation can take place (Figure 2a). This widespread dysregulation was not driven by individual genes, such as particular chromatin modifiers or TFs, indicating a potential contribution of feedback loops or synergistic regulation (Supplementary Results).

**Figure 2.**
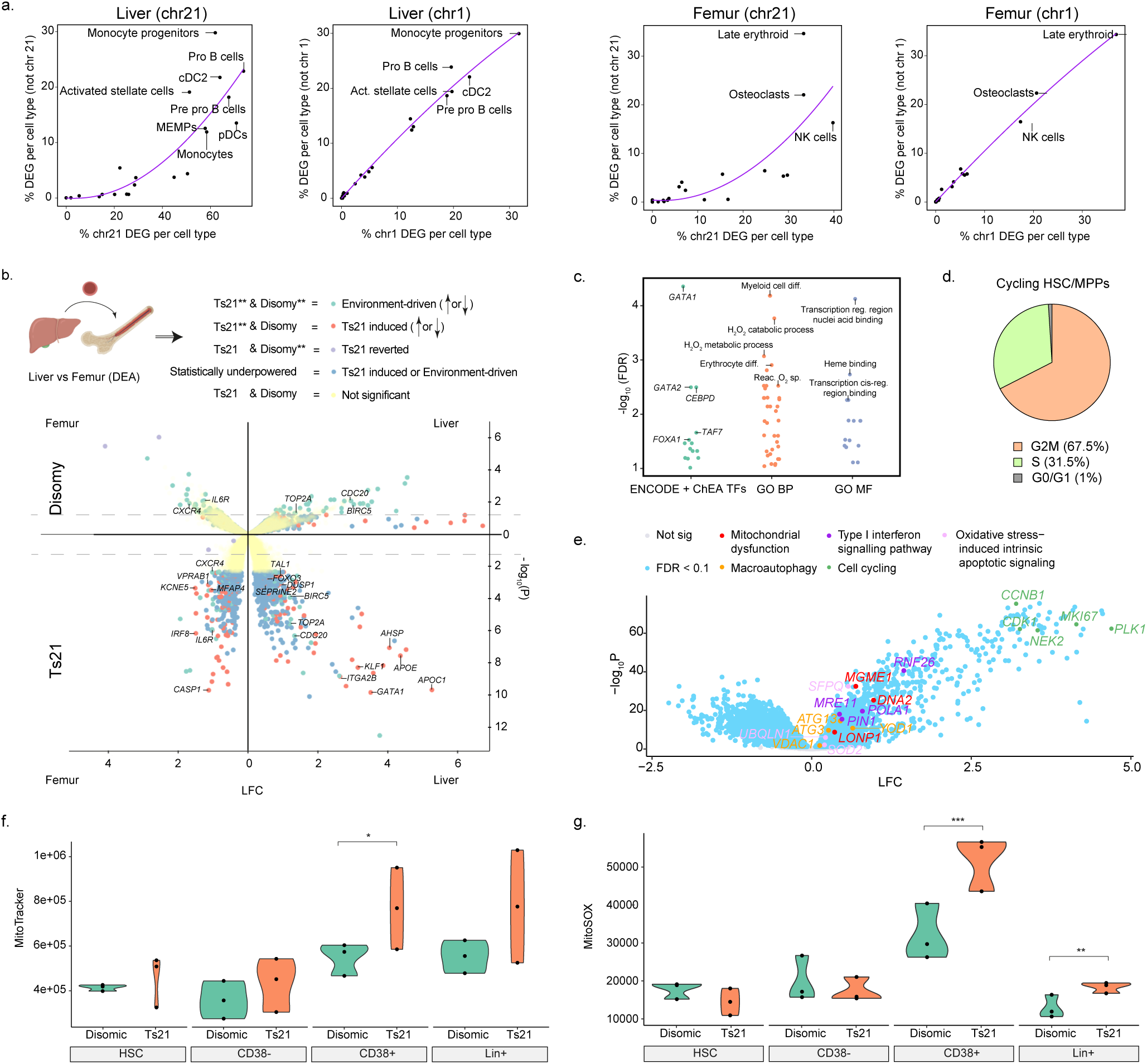
Gene-Environment interactions in Ts21 and disomic haematopoietic cells. **a,** The percentage of differentially expressed genes (DEG) that are significant (FDR < 0.05) on chromosome “N” versus all other chromosomes. Each point represents a different cell type tested. **b,** Within both Ts21 cells and disomic cells, differential expression analysis was performed between liver and femur samples. The genes identified within HSC/MPPs were characterised as *Environment-driven*, *Ts21-induced*, or *Ts21-reverted* by downsampling Ts21 liver and femur cell counts to the same size as disomic liver and femur data (in order to account for sample and cell count differences). *Environment-driven* DEGs had evidence of expression differences in both Ts21 and disomic cells. *Ts21-induced* DEGs were only discovered in Ts21 cells, including when accounting for sample and cell count differences. *Ts21-reverted* DEGs included the genes discovered only in disomic cells. Reduced statistical power led to DEGs found in only the full Ts21 data but not in the downsampled Ts21 or disomic data, such that they could not be conclusively characterised as Ts21-induced or Environment-driven. A volcano plot shows DEGs between HSC/MPPs isolated from foetal liver or BM from Ts21 (lower part of the plot) or disomic foetuses (upper part of the plot), with the x-axis displaying log_2_ fold-change (magnitude of change) and the y-axes showing the -log_10_ FDR (statistical significance). **c**, Gene-set enrichment analysis of upregulated *Ts21-induced* genes was performed using EnrichR. Scatterplots show the top Gene Ontology terms (GO:MF - Molecular Function, GO:BP - Biological Process) or ENCODE and ChEA TFs. **d,** Pie chart displaying the percentage of cells in G1, G2M, and S in cycling HSC/MPPs sorted from Ts21 foetal liver. **e,** Volcano plot between cycling HSCs/MPPs and normal HSCs/MPPs in Ts21. Dots represent genes, with sky blue coloured genes having FDR < 0.05 and logFC > 1. Some genes are specially coloured according to their GO term. In panels **a-e**, analyses were performed using all n=1,107,552 cells and k=18 foetuses, represented by Ts21 liver (n=780,299, k=15), Ts21 femur (n=162,775, k=12), disomic liver (n=110,671, k=3) and disomic femur (n=53,807, k=3). **f-g, Quantitative comparison of** MitoTracker staining (**f**) and MitoSOX staining (**g**) in disomic and Ts21 haematopoietic populations; HSC-haematopoietic stem cells (Lin-/CD34+/Cd38- /CD90+/CD45RA-); CD38- (Lin-/CD45+/CD34+/Cd38- population); CD38+ (Lin-/CD45+/CD34+/CD38+ population); Lin+ (CD3/8/11b/56/14/19, lineage positive population).

As Ts21 leukaemia is believed to originate from foetal liver HSCs^4^, we analysed gene-environment interactions specifically in HSCs. We contrasted the gene expression of HSC/MPPs collected from disomic and Ts21 foetuses between liver and bone marrow environments. By subsampling liver HSCs to balance the number of cells and donors across genotypes and environments, we discerned consistent changes in gene expression between liver and BM that are present independent of Ts21 genotype (environment-driven) *versus* those differences that are only present in Ts21 cells (Ts21-induced) (Figure 2b; see Methods; Supplementary Results, Supp. Figure 12e). Compared to HSC/MPPs in BM, we found that both disomic and Ts21 HSC/MPPs in liver up-regulated genes involved in the cell cycle such as *TOP2A*, *CDC20*, and *BIRC5*, and down-regulated genes involved in cytokine response such as *CXCR4* and *IL6R* (Figure 2b). Subsequent cell cycle analysis confirmed that HSC/MPPs, both in Ts21 (G2/M+S=29.8%) and disomic liver (21.8%) (Supp. Figure 12f, left panel), were cycling more compared to their counterparts in BM (26.9% in Ts21 and 18.3% in disomic, Supp. Figure 12f, right panel)^34^.

Additionally, we identified “Ts21-induced” changes in gene expression that were specifically present in Ts21 foetuses and upregulated in the liver HSC/MPPs compared to BM. These included transcription factor and chromatin binding genes such as *GATA1*, *TAL1*, and *FOXO3*, as well as genes involved in erythrocyte differentiation (*KLF1*, *AHSP)* and haemostasis (*ITGA2B, SERPINE2)* (Figure 2b, c). Notably, genes involved in response to ROS, (*FOXO3*, *APOE*, *DUSP1*) were also specifically upregulated in HSC/MPPs in Ts21 foetal liver compared to BM (Figure 2b, c). Oxidative stress is a proposed cause of GATA1 mutations observed in Ts21 preleukaemia and acute megakaryocytic leukaemia, as suggested by prior mutational signature analysis^11^. Furthermore, oxidative stress in HSC/MPP can stimulate cycling and cause their premature exhaustion^35,36^. This is in line with the presence of “cycling” HSC/MPPs (G0/G1=1%) in Ts21 liver (Figure 2d). Cycling HSC/MPPs comprised 25% of the total HSCs pool in Ts21. Besides increased expression of cell cycle genes (*MKI67, CCNB1, PLK1, CDK1, NEK2*), these cycling HSCs in Ts21 liver showed upregulation of genes involved in oxidative stress-induced intrinsic apoptopic signaling (*SFPQ*^37^, *UBQLN1*^38^*, SOD2*^39^), mitochondrial dysfunction (*MGME1*^40^, *DNA2*^41^, *LONP1*^42^), macroautophagy (*ATG3/13*^43^, *VDAC1*^44^*, YOD1*^45^) and type I interferon signalling pathway (*RNF26*^46^, *POLA1*^47^*, MRE11*^48^*, PIN1*^49^), compared to Ts21 liver HSC/MPPs (Figure 2e). It has been previously demonstrated that mitochondrial dysfunction contributes to the number of DS-associated pathologies and severity of the phenotypes^50,51^. In addition, reduced mitochondrial activity, through increased autophagy of mitochondria, can lead to increased asymmetrical division of HSCs, enlarged pool of progenitors and lineage bias^52^. Overall, by including cells from both liver and BM in our analysis, we observed a complex interplay between genetic background and environment with distinct consequences on gene expression and cellular phenotypes of HSCs. Critically, the increase in ROS, a potent mutagen^10^, was evident in Ts21 HSCs in liver but not in femur, and further exacerbated in the subset of highly cycling HSCs.

### Haematopoietic cells in Ts21 have higher mitochondrial mass and oxidative stress

To validate higher oxidative stress and altered mitochondrial mass in Ts21 foetal haematopoietic cells compared to disomic cells (Figure 2f, g, Supp. Figure 13a), we measured mitochondrial reactive oxygen species (mtROS) levels with MitoSOX probe and mitochondrial mass using MitoTracker green FM dye. MitoSOX is a specific fluorogenic dye for live-cell mitochondria, producing bright green fluorescence upon oxidation by mitochondrial superoxide. MitoTracker green FM accumulates in mitochondria independent of membrane potential and oxidative stress, serving as a reliable tool for mitochondrial mass measurement^53^. Considering dye efflux bias in HSCs and progenitor cells^54^, (Supp. Figure 13b, e), we employed Verapamil treatment to block xenobiotic efflux pumps and mitigate preferential dye efflux (Supp. Figure 13b, e), ensuring a more accurate representation of mitochondrial mass and mtROS levels.

We compared the mitochondrial mass and mtROS levels in Ts21 (n=3) and disomic foetal liver samples (n=3) stained in the presence of Verapamil (Figure 2f, g). We observed that Ts21 CD34+CD38+ cells had higher mitochondrial mass and mtROS levels compared to disomic progenitors, whereas CD34+CD38- and HSCs were comparable between the two conditions (Figure 2f, g). Previous studies have shown that disomic HSCs exhibit very low electron transport chain activity and minimal mROS production as they mostly rely on glycolysis^55^. In contrast, committed progenitors have much higher oxygen consumption, mitochondrial ATP production, and maximal respiratory capacity. Taken together, these results support the hypothesis that as the HSCs and immature CD34+CD38- cells start to differentiate to committed CD34+CD38+ progenitors, they ramp-up oxidative phosphorylation. This leads to higher mtROS production in Ts21 cells compared to disomic cells, possibly due to their higher mitochondrial mass and dysfunction in the electron transport chain activity, which has been implicated in oxidative DNA damage, mutagenesis and leukaemia development^56^.

### Multiome analysis reveals erythroid priming in Ts21 HSCs in foetal liver

Our previous work has demonstrated that changes in motif accessibility in disomic HSCs precede activation of lineage specific transcriptional programs and their cell fate choice^34^. To further investigate if changes in the regulatory landscape between Ts21 and disomic haematopoietic cells contribute to differences in cell type abundances, we performed 10X Multiome sequencing on haematopoietic CD45+ cells from Ts21 (n=3) and disomic foetal liver (n=3). After QC (see Methods), we obtained high-quality open chromatin profiles for 35,633 nuclei from Ts21 (a median of 13,476 unique open chromatin fragments per nucleus) and 21,257 nuclei (median = 16,186 fragments per nucleus) from disomic samples (Supp. Figure 14a, Supp. Table 10). Next, we identified cell populations within Ts21 and disomic samples separately by clustering the snRNA-seq Multiome data (see Methods). Overall, we found cell-type clusters that represent all broader cell types identified in the larger scRNA-seq atlas (Supp. Figure 14b), with transcriptionally distinct cell populations appearing epigenetically distinct within the UMAP of the paired scATAC-seq multiome data (Supp. Figure 14b).

Within the Multiome data, we observed higher erythroid cell proportions, in line with our analysis of scRNA-seq data (Supplementary Results). Following integration of disomic and Ts21 datasets (Figure 3a), we inferred trajectories of myeloid lineage cell populations (Supp. Figure 15a, b) and estimated the likelihood of each individual HSC to transition toward each lineage (see Methods). We found that HSCs displayed the ability to commit toward all lineages; however, Ts21 HSCs showed much greater bias towards the erythroid lineage compared to disomic HSCs (Figure 3b). Furthermore, we found that the GATA1 (a TF essential for erythroid lineage development) gene body and promoter were accessible in a higher percentage of Ts21 HSCs compared to disomic HSCs (19.3% versus 6.7%). This provided evidence that the erythroid lineage skewing starts already at the level of HSCs.

**Figure 3.**
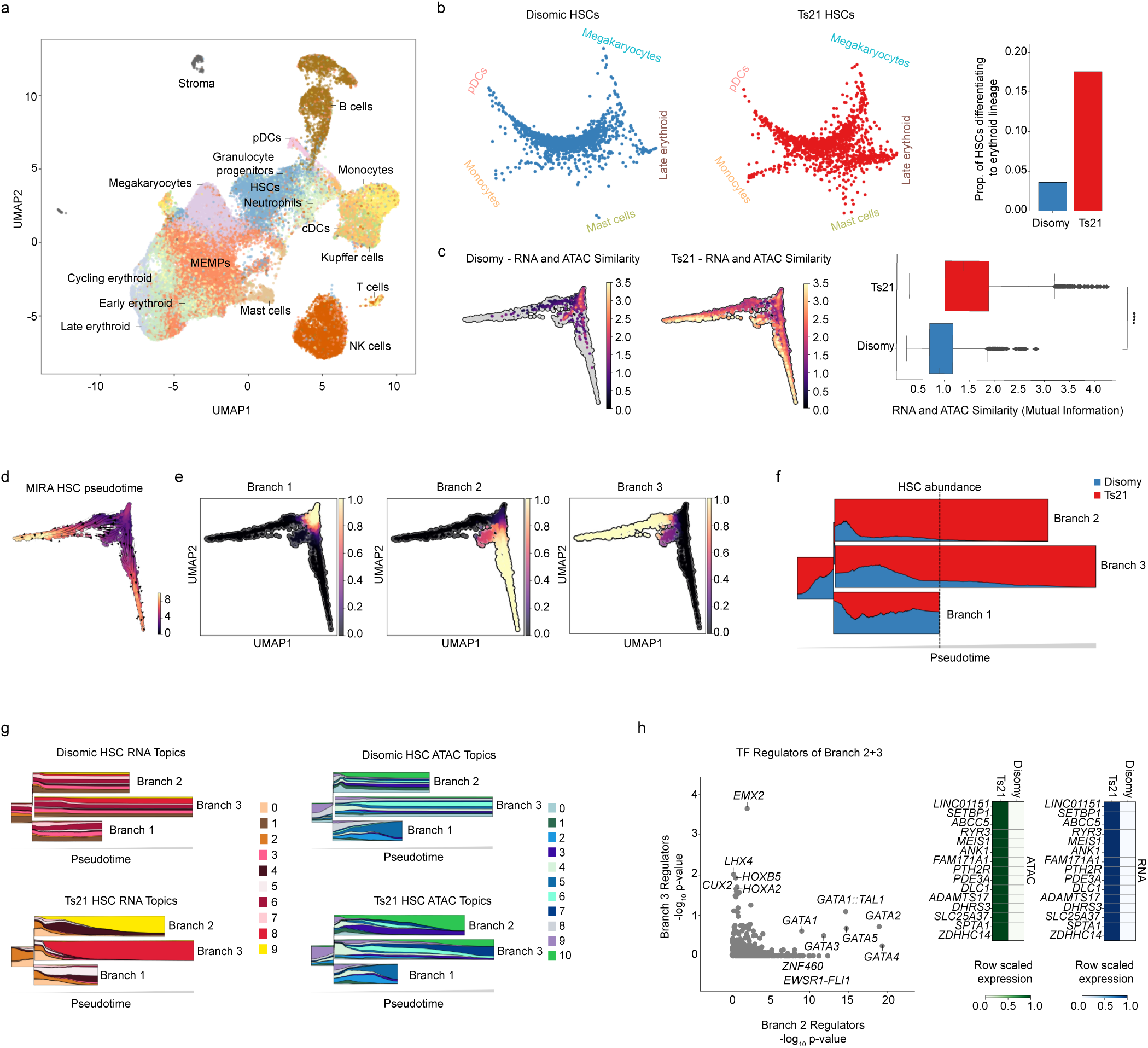
Multiome analysis. **a,** Integration, UMAP, and cell population labels of 10X multiome ATAC profiles from the Ts21 and disomic datasets. MEMPs, megakaryocyte-erythroid-mast progenitors. HSCs, haematopoietic stem cells. MΦ, macrophages. cDCs, classical dendritic cells. pDCs, plasmacytoid dendritic cells. **b,** Left, circular projection of terminal state absorption probabilities. Probability of every HSC moving toward each of the 5 terminal states identified by CellRank (pDCs, Megakaryocytes, Late erythroid, Mast cells, Monocytes). Blue, disomic HSCs. Red, Ts21 HSCs. A cell in the centre represents an equivalent likelihood of differentiating toward all terminal states, while cells close to a terminal state have a greater likelihood of differentiating toward that state. Right, proportion of HSCs differentiating toward the erythroid lineage indicated by an absorption probability > 0.5 (disomic=0.04, Ts21=0.18). **c**, RNA and ATAC similarity in disomic and Ts21 HSCs, as estimated by the mutual information of RNA and ATAC topics in a single cell. HSC latent UMAP space characterised by a joint representation of both RNA topic models and ATAC topic models as characterised by MIRA. **d**, UMAP with cells coloured according to pseudotime (dark=early, light=late). Black arrows denote vectors of the directed cell transition matrix. **e**, Representation of three branches of HSC differentiation in UMAP space as identified by MIRA. Cells are coloured according to the probability of each cell belonging to a specific branch (branch 1: left, branch 2: middle, branch 3: right). **f**, Compositional STREAM graph indicating the abundance of either disomic or Ts21 HSCs in each branch along pseudotime. As HSCs progress along MIRA calculated pseudotime, the abundance of each type of HSC (disomic: blue, Ts21: red) was calculated using a sliding window according to the branch identity of the cell. Dotted line indicates half the pseudotime (t=5). **g**, Compositional STREAM graph indicating the proportion of topic abundance within cells ordered along pseudotime according to each of the three branches. Total topic number was automatically optimised by MIRA. Left, 10 RNA expression topics. Right, 11 ATAC accessibility topics. For Ts21 HSCs (bottom row), as pseudotime progresses, both branch 2 and branch 3 become dominated by a few distinct topics. Within disomic HSCs (top row), there remains an even distribution for both RNA and ATAC topics. **h**, Left, scatter plot comparing the transcription factors (TF) regulating the expression of genes turned on specifically in branch 2 and branch 3. TF association scores calculated using MIRA’s probabilistic *in-silico* depletion function. P-values calculated using a Wilcoxon rank sum test over the association scores. Right, top genes regulated by labelled branch 2 TFs. The expression (blue) and accessibility (green) indicate Ts21 HSC specificity. Data is normalised by row to illustrate preference between disomic and Ts21. Analyses were performed using all n=56,890 cells and k=6 foetuses, represented by Ts21 liver (n=35,633 cells, k=3) and disomic liver (n=21,257, k=3), from which n=6,215 cells are HSCs (n=3,784 Ts21 and n=2,431 disomic HSCs).

To assess how the epigenetic changes lead to the observed erythroid biases, we examined the relationship between chromatin accessibility and gene expression in Ts21 compared to disomic HSCs. While chromatin accessibility and gene expression are highly concordant in terminally differentiated cells, chromatin remodelling and gene expression are often out of sync in HSCs, such that a gene’s promoter and enhancers may be open but little to no transcription occurs^34^. To contrast RNA and ATAC profiles from the same cell, we described the transcriptional and chromatin accessibility state of single cells as a group of co-regulated genes (RNA topics) or *cis*-regulatory elements (ATAC topics) respectively^57^ (see Methods; Supp. Table 11). By quantifying individual cells’ accessibility and expression within topics, we found that Ts21 HSCs showed a greater concordance between RNA and ATAC topic compositions compared to disomic HSCs, suggesting they are “primed” to differentiate^34,57^ (Figure 3c, Supp. Figure 15c).

Since a subset of Ts21 HSCs showed differentiation bias toward the erythroid lineage, we sought to further dissect their heterogeneity compared to disomic HSC. We inferred a cell state tree^57^ (see Methods), which revealed HSCs organisation with three main branches that contained both Ts21 and disomic cells (Figure 3d, e), but with different abundances (Figure 3f). Branch 1 had a similar proportion of Ts21 and disomic HSCs. In contrast, branches 2 and 3 consisted predominantly of Ts21 HSCs, especially later in the pseudotime trajectory (Figure 3f). We then used the topics to characterise HSC cell states. Along inferred pseudotime, RNA topics appeared to differ more between Ts21 and disomic HSCs than ATAC topics (Figure 3g), suggesting larger transcriptional than epigenetic differences in Ts21. Furthermore, the RNA topic composition of HSCs in branch 1 was distributed across multiple topics, which suggests the maintenance of a multipotent state (Supp. Figure 15d). In contrast, HSCs in branch 2 highly expressed RNA topic 9, which contains genes related to “heme metabolism” (Supp. Figure 15d, e-left), and were cycling as evidenced by high expression of *MKI67* (Supp. Figure 15f). The main RNA topic associated with branch 3 was topic 8 and was associated with “down-regulated genes in blood” (Supp. Figure 15d, e-right). Our findings indicate that Ts21 HSCs encompass a diverse population of cells that share transcriptional and epigenetic similarities with disomic HSCs. However, a distinct subset within Ts21 HSCs displays an increased erythroid bias, a characteristic not observed in disomic samples (branch 1).

We next identified TFs that regulate the expression of genes specific to each of the three HSC branches by simulating “computational knockouts” of each TF (see Methods). The key TFs regulating HSCs in branch 1 included *STAT2*, *ETV1/4*, and *IKZF1* (Supp. Table 12); in branch 2 (defined by topic 9): the *GATA* family, *ZNF460*, and *EWSR-FLI1* (FDR < 0.05; Figure 3h, Supp. Table 12); and in branch 3 (defined by topic 8), there were no topics significantly driven by particular TFs (FDR < 0.05), but there was suggestive evidence of regulation by TFs such as *LHX4*, *CUX2*, *EMX2*, and the *HOX* family (e.g. *HOXA2* and *HOXB5*) (nominal P < 0.05), (Figure 3h, Supp. Table 9). Several genes regulated by the *GATA* family^58^ TFs relevant to branch 2 are known regulators of erythropoiesis and were specifically expressed in Ts21 cells (Figure 3h). Further, we found evidence that additional factors beyond local accessibility (such as additional signalling) are necessary to enact the expression of erythroid lineage genes specific for Ts21 cells in branch 2 (Supp. Results, Supp. Figure 16). Thus overall, we observed systemic evidence, on both chromatin accessibility and transcriptional levels, that a subpopulation of Ts21 HSCs is “primed” to differentiate towards erythroid lineage.

### Ts21 significantly perturbs interactions between enhancers and their target genes

Given that epigenetic and transcriptional priming was prevalent in Ts21 HSCs, we aimed to identify regulatory elements and causal genes that potentially drive the observed lineage bias in Ts21. We first set out to identify maps of enhancer-gene relationships in disomic and Ts21 HSCs. We then used these maps to assess whether trisomy influences enhancer-gene interactions to alter gene expression. We established maps of enhancer-gene relationships in disomic and Ts21 HSCs by correlating peak accessibility with gene expression (Figure 4a; see Methods). We identified 191 and 1,564 significant peak-gene links in disomic and Ts21 HSCs respectively (FDR < 0.2), with a greater number of links being identified in Ts21 HSCs even after downsampling cells to the disomic HSC count (773 significant links; 4.1-times more) (Figure 4b, c; Supp. Table 13). Of all identified peak-gene links, only 69 were shared between disomic and Ts21 HSCs (representing 36.1% of those found in disomic HSCs). Thus, we tested whether Ts21 modifies the effect of peak accessibility on gene expression by using an accessibility-by-trisomy interaction term (see Methods, Figure 4c). We found a significant interaction term for 62.1% (72/116) of all significant peak-gene links that were specific to disomic HSCs. This finding shows that there is a widespread loss of disomic peak-gene links in the Ts21 background (Figure 4c).

**Figure 4.**
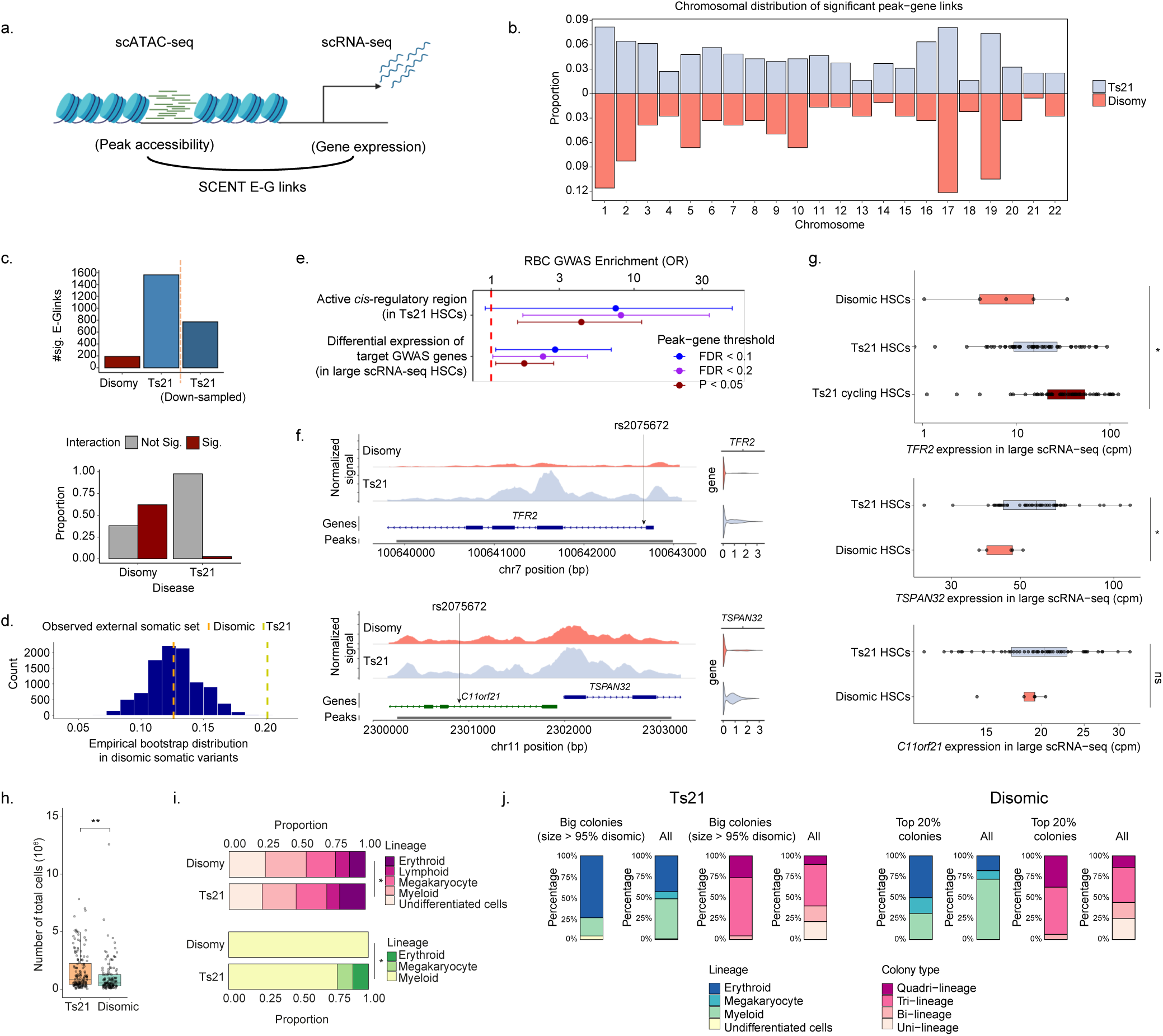
Integration of peak-gene links and GWAS SNPs. **a,** Schematic of the SCENT peak-gene multiome analysis performed in n=6,215 HSCs (n=3,784 Ts21 and n=2,431 disomic HSCs)**. b,** Distribution of significant peak-gene links identified in Ts21 (top) or disomic HSCs (bottom) (FDR < 0.2). The y-axis represents the proportion of links that are on that chromosome versus genome-wide. **c**, Top, number of significant peak-gene (E-G) links in Ts21 and disomic HSCs, including after downsampling Ts21 HSCs; Bottom, proportion of significant peak-gene links with a significant accessibility-by-trisomy interaction term. **d**, Cataloged Ts21 mutations in the introns of Ts21 HSC-expressed genes (derived from scRNA- seq) are more likely to overlap candidate cis-regulatory elements (CREs) from the ENCODE project than catalogued intronic mutations from disomic cells. The histogram represents the empirical distribution of this overlap from 10,000 bootstrapped sets of somatic variants found in disomic HSCs. **e,** Top, enrichment of fine-mapped GWAS SNPs from red blood cell counts (RBC) (PIP > 0.2) in SCENT peak-gene links (termed active *cis*-regulatory regions) as identified in Ts21 HSCs. Bottom, enrichment of target GWAS genes (based on Ts21 SCENT peak-gene links and using fine-mapped RBC SNPs) within differentially expressed genes, as identified in the large scRNA-seq analysis of HSCs (FDR < 0.1). Fisher’s exact test was performed for calculating odds ratios and standard errors. The analysis was repeated using three sets of peak-gene links based on SCENT significance: FDR < 0.1, FDR < 0.2, and nominal P-value < 0.05. **f,** aggregated accessibility and expression of HSCs from disomic and Ts21 HSCs for *TFR2* and *TSPAN32* using 10X multiome data. **g**, Pseudobulk expression within HSC populations using the large scRNA-seq data. Each point represents a different sample. **h-j,** 322 colonies generated by single-cell sorted HSC/MPPs from 3 independent disomic and Ts21 foetal livers. **h,** The total number of cells (live and dead) in Ts21 and disomic HSC-derived colonies. The difference between disomic and Ts21 colonies was determined by a Wilcoxon Rank Sum test (*P*=1.3×10^-3^). **i**, The upper bar graph shows proportions of different lineages detected in either uni-, bi-, or multi-lineage colonies. The lower bar graph shows proportions of different lineages detected in uni-lineage colonies. Statistical differences between foetuses with Ts21 and the expected distribution based on disomic foetuses were tested by a chi-squared test (p=0.01148 and p=0.0111, respectively). **j,** The lineage output of “big colonies” and “all” Ts21 HSC/MPPs-derived colonies. Big Ts21 colonies were defined as colonies in which the total number of cells was higher than in 95% of disomic HSC/MPPs colonies, representing 20% of all Ts21 colonies. For comparison, the top 20% disomic colonies (ranked by cell count) were analysed. The main lineage output of each colony was defined by the lineage with the highest cell count, determined by flow cytometry.

In contrast, only 2.4% (20/839) of Ts21-specific peak-gene links had a significant interaction term suggesting that trisomy strengthens a peak-gene connection. Compared to all tested peaks and genes, the Ts21-specific peaks and genes (without a significant interaction) were significantly more accessible and upregulated in Ts21 compared to disomic within differential accessibility analysis (1.9-fold enrichment; P-value = 5.6×10^-12^) and the large scRNA differential expression analysis (1.7-fold enrichment; P-value = 9.7×10^-7^). This suggests that the greater activation of those regulatory elements and more widespread transcription (i.e. more genes expressed and higher expression) in Ts21 HSCs allowed these peak-gene links to be discovered. Finally, 21.7% (15/69) of peak-gene links detected in both disomic and Ts21 HSCs (FDR < 0.2) had a significant interaction, resulting in the dampening or strengthening of the peak-gene relationship in Ts21. There was one example (*RAPGEF2*^59^) where the peak- gene link was directionally discordant. Deficiency of *RAPGEF2* has been shown to lead to defective foetal liver erythropoiesis^59^ and it was upregulated in the Ts21 HSCs (*P* = 0.04). Therefore, our results indicate that trisomy restructured the enhancer-gene landscape by altering the correlation between peak accessibility and gene expression, which led to a substantial increase in accessibility and upregulation of these elements in Ts21 compared to the control.

Our findings offer a potential link between altered chromatin organisation in Ts21 and oxidative stress with somatic mutations accrual, thus providing an alternative perspective on the aetiology of leukaemia in DS. Notably, chromatin organisation is well-known to impact regional variation of mutation rate in different cancers^60^, resulting in preferential accumulation of mutations in distinct chromatin regions. To test if Ts21 induced changes in chromatin accessibility impact mutagenesis in regulatory regions of genes, we examined patterns of somatic mutations identified in foetal Ts21 and disomic HSCs from Hasaart et al^10^. We found that Ts21 mutations located in the introns of Ts21 HSC-expressing genes are 60% more likely to reside within ENCODE cis-regulatory elements compared to intronic mutations identified in disomic HSCs (P = 3×10^-4^; Figure 4d). This result is in line with the hypothesis that altered chromatin accessibility landscape in Ts21 HSCs allows differential mutation acquisition in regulatory regions of genes, with the potential to increase the propensity of Ts21 HSCs to preleukaemia and leukaemia development.

### GWAS SNPs identify regulatory elements that drive red blood cell differentiation bias in Ts21 HSCs

Our peak-gene analysis identified active regulatory elements and their target genes in Ts21 and disomic HSCs. To pinpoint which regulatory elements are relevant for different blood cell types, and in particular for erythroid lineage bias in Ts21, we used published GWAS loci relevant for blood cell traits and evaluated their activity in HSCs. Genome-wide association studies (GWAS) have identified a myriad of genetic variants associated with a range of blood cell traits and diseases, many of which are non-coding and impact causal genes through *cis*- regulatory elements. We used variants associated with blood cell variation because many have been implicated in leukaemia risk despite limited knowledge of the genetic architecture of leukaemia risk^61^.

First, we evaluated the overall accessibility of regulatory elements important to blood differentiation across HSC branches. By tagging key non-coding elements using fine-mapped GWAS loci, we identified individual cells with increased accessibility for red blood cell (RBC), white blood cell (WBC), and lymphoid cell (Lymph) traits^62,63^. As expected, we found significant enrichment of GWAS SNPs in open chromatin regions for Lymph counts, WBC counts, and RBC counts within their corresponding cell types (Supp. Figure 17a, b, Supplementary Results), thus confirming the identity of annotated cell types and the utility of the approach. We next explored enrichment of blood cell traits in different HSC branches (identified by the cell state tree) from Ts21 and disomic foetuses. We found that HSCs from branch 1 were significantly enriched for cells with increased accessibility of GWAS SNPs relevant to WBC count, while HSCs from branch 2 were significantly enriched for cells with increased accessibility of RBC count GWAS SNPs (Figure 17b; FDR < 0.1). Considering that the branch 2 was mostly composed of Ts21 HSCs, our analysis suggests that RBC GWAS-harbouring enhancers were more accessible in a subset of Ts21 HSCs.

Next, we assessed whether RBC GWAS-harbouring enhancers influence gene expression. We intersected fine-mapped RBC GWAS SNPs with peak-gene links and Ts21 differentially expressed genes from our large scRNA-seq atlas. We found that fine-mapped GWAS- harbouring peaks were enriched for association with gene expression compared to peaks that do not contain GWAS SNPs (Figure 4e; *P* = 0.03). Furthermore, target genes of GWAS- harbouring peaks (as identified by peak-gene links) were more likely to be differentially expressed between disomic and Ts21 HSCs in our scRNA-seq data (*P* = 0.01), confirming the functional relevance of these peaks (Figure 4e).

As one example, the intronic variant rs2075672 lies in a peak primarily accessible in Ts21 HSCs (*P* = 5.0×10^-3^) and its accessibility correlates with *TFR2* expression (Figure 4f-top, g; *P* = 2×10^-4^), which has been implicated in the regulation of erythropoiesis^64^. Within our scRNA-seq dataset, *TFR2* expression was significantly higher in Ts21 HSCs compared to disomic HSCs (*P* = 1.8×10^-3^), with even higher expression found in Ts21 cycling HSCs (*P* = 3.5×10^-31^). Notably, integration of the Multiome dataset with the scRNA data also informed likely causal genes in blood cell development at GWAS-harbouring enhancers. We found that the *C11orf21* intronic peak carrying the fine-mapped RBC SNP rs2077078 had increased accessibility in Ts21 HSCs (*P* = 1.4×10^-3^) and its accessibility was associated with both *C11orf21* and *TSPAN32* expression (*P* < 0.03) (Figure 4f-bottom, g). However, based on our differential expression analysis of Ts21 and disomic liver HSCs in the scRNA dataset, only *TSPAN32* was dysregulated in Ts21 HSCs (*P* = 1.4×10^-3^) (Figure 4f, g). This suggests that *TSPAN32* can contribute to the observed lineage bias and that *TSPAN32* is the likely causal gene of the rs7775698-linked enhancer and its role in red blood cell development. While dysregulation of any individual gene would have only a minor effect on HSC differentiation, genome-wide reshaping of enhancer-gene maps in Ts21 is likely to have a major impact on haematopoiesis.

### Lineage bias validates *in vitro*

To confirm that the observed erythroid lineage bias can be recapitulated *in vitro*, we examined the differentiation potential of individual HSC/MPPs (CD34+, CD38-, CD62L+, CD52+) sorted (Supp. Figure 18a) from disomic (n=3) and Ts21 (n=3) foetal livers into 96-well plates without a feeder layer. After two weeks, 322 colonies were assessed for their size and the lineage output by flow cytometry [CD41a (megakaryocytic-Mk), CD235a (erythroid-Ery), CD3/CD56 (lymphoid-Ly) and CD11b (myeloid-My)] (Supp. Figure 18b). Although there was no significant difference in the proportion of colonies that consisted of one (uni-) or multiple lineages, the overall frequency of myeloid, lymphoid, MK, and Ery lineages showed significant differences (p=0.01) between trisomy and disomy (Figure 4i). In the disomic samples, uni-lineage colonies exclusively contained myeloid cells; whereas in Ts21, MK- and Ery- colonies were present as well (Figure 4i).

To assess cell proliferation of foetal liver HSPCs, we counted the total number of cells in individual colonies derived from individually sorted HSCs. Colonies obtained from Ts21 foetal liver HSCs showed a significantly higher number of total cells compared to the controls (*P* = 1.3×10^-3^; Figure 4h). Furthermore, Ts21 colonies were biased towards the Ery lineage, compared to disomic colonies (Figure 4j). This was markedly pronounced when comparing the largest colonies. The top 20% of Ts21 colonies were bigger than the 95th size percentile of disomic colonies; and, while 50% of the largest disomic colonies (top 20% in size) differentiated towards the Ery lineage, 73.9% of the largest Ts21 colonies (top 20% in size) had Ery lineage output (*P* = 0.035) (Figure 4j). These results support the notion that Ts21 HSCs show megakaryocytic and erythroid bias *in vitro* and that the observed lineage bias is more pronounced in cycling HSCs.

## Discussion

Our study represents the largest and most comprehensive single-cell multi-omics study of blood development in Down Syndrome to date. Compared to a previous study^65^, we increased the size over 50-fold to 1.1 million scRNA-seq cells, collected from haematopoietic and niche cells from matched foetal liver and bone marrow, and interrogated the spatial and epigenomic landscapes using spatial transcriptomics and 10X Multiome. This rich resource enabled discovering the molecular changes that occur in haematopoiesis as cells migrate from foetal liver to bone marrow, and further establishing how trisomy 21 impacts this process to drive haematological abnormalities and elevated leukaemia risk.

Our analyses revealed the key importance of cellular and genomic context for studying foetal hematopoiesis in DS. Expression differences between Ts21 and disomic cells were dependent on both cell type and environment. By studying these expression differences in-depth within HSCs across disomic and trisomic genetic backgrounds and liver and femur environments, we found that Ts21 liver HSCs exhibited higher levels of cycling behaviour, oxidative stress, and mitochondrial dysfunction. Furthermore, while oxidative stress has been proposed as a mutational process involved in DS blood samples, we experimentally validated increased mitochondrial mass and reactive oxidative species as phenotypes of Ts21 haematopoietic cells.

Down Syndrome infants are often born with high counts of and defects in red blood cells^6^. By analysing epigenomic and transcriptomic profiles, we found that Ts21 HSCs, while multipotent, exhibited an erythroid lineage bias. Compared to disomic HSCs, they exhibited greater concordance between RNA and ATAC profiles, which is a feature typical of mature cell states^34,57^. Further, Ts21 restructured the enhancer-gene map to increase accessibility and expression of regulatory elements and their genes. This effect was pronounced in enhancers critical to red blood cell differentiation (as tagged by GWAS variants), implying a genome-wide and regulatory role in impacting erythroid lineage differentiation. Thus, our results show that chromatin remodelling underlies the priming of Ts21 HSCs toward the erythroid lineage, which we further confirmed within *in vitro* differentiation assays.

Finally, we can hypothesise that the interplay between altered chromatin organisation, gene expression dynamics, and the strain of oxidative stress coalesces to create an environment potentially conducive to pre-leukemic and leukaemic changes. Intriguingly, our investigation revealed that DS-linked mutations^10^ are concentrated within regulatory regions^66^ of genes expressed by HSCs, hinting at a plausible mechanistic link between chromatin alterations, oxidative stress, and heightened leukaemia risk in Ts21. A future goal is to assess this hypothesis by pairing cell-type-specific whole genome sequencing with multi-omic data across cellular and tissue contexts in the same samples.

## Methods

### Ethics and Tissue acquisition

Human foetal bone and liver samples were obtained from 15 Ts21 foetuses aged 12-20 pcw and 5 disomic foetuses aged 11-19 pcw, following termination of pregnancy and informed written consent. The human foetal material was provided by the Joint MRC/Wellcome Trust (Grant MR/R006237/1) Human Developmental Biology Resource (http://www.hdbr.org), with maternal informed consent, in accordance with ethical approval by the National Health Service (NHS) Research Health Authority, REC Ref: 18/LO/0822. HDBR is regulated by the UK Human Tissue Authority (HTA; www.hta.gov.uk) and operates in accordance with the relevant HTA Codes of Practice.

### Dissociation of foetal tissues

Foetal livers and femurs were received in L15 media and processed within 3h from dissection. Livers were cut in smaller pieces with a scalpel and transferred to a tube containing prewarmed digestion media: RPMI (Gibco) supplemented with 10% FBS (Gibco), penicillin/streptomycin (10U/ml penicillin, 100ng/ml streptomycin, Sigma Aldrich), 2mM L-glutamine (Thermo Scientific), 1x MEM NEAA (Gibco), 1mM Sodium pyruvate (Gibco) and 1.6mg/ml Collagenase-IV (Sigma Aldrich). The tube was vortexed for 10”, then incubated at 37°C for 30 minutes, and vortexed for 10” every 15 minutes. The digested tissue was filtered through a 100μm filter and diluted in cold D-PBS (Gibco). Cells were centrifuged at 300g for 5 minutes, then aliquoted and cryopreserved in KnockOut Serum Replacement (Gibco) + 5% DMSO (Sigma Aldrich). For femurs, adherent material was removed, then the epiphyses were removed with a scalpel and the bone marrow flushed with D-PBS. The remaining bone was cut in small pieces, then ground with a mortar and pestle using digestion media and incubated at 37°C for 30 minutes, vortexing every 15 minutes. The digested material and the bone marrow flush were mixed and filtered through a 100μm filter. Cells were centrifuged at 300g for 5 minutes, then aliquoted and cryopreserved in KnockOut Serum Replacement (Gibco) + 5% DMSO (Sigma Aldrich). Cells were stored in liquid nitrogen until further analysis.

The karyotype for each sample used in this study was determined by Quantitative fluorescent PCR (QF-PCR). The analysis was performed by the tissue bank from which the samples were obtained. QF-PCR was performed using chromosome specific microsatellite markers. The analysis showed normal results with an apparently normal diploid complement for chromosomes 13, 15, 16, 18, 21, 22 and the sex chromosomes in disomic samples and trisomy for chr 21 in Ts21 samples. No mosaicisms were detected.

### FACS sorting for scRNA sequencing

On the day of FACS sorting, cells were rapidly thawed at 37°C and transferred to complete RPMI media (RPMI (Gibco) supplemented with 10% FBS (Gibco), penicillin/streptomycin (10U/ml penicillin, 100ng/ml streptomycin, Sigma Aldrich), 2mM L-glutamine (Thermo Scientific), 1x MEM NEAA (Gibco), and 1mM Sodium pyruvate (Gibco)). Live cell enrichment was performed using MACS Dead Cell Removal Kit (Miltenyi Biotec, cat#: 130-090-101) following the manufacturer’s instructions. When depleting for CD235a+ cells, a magnetic negative selection was performed using CD235a Microbeads (Miltenyi Biotec, cat#: 130-050- 501) and MACS LS columns (Miltenyi Biotec, cat#: 130-042-401) following the manufacturer’s instructions.

For FACS sorting, cells were stained with Zombie Aqua to exclude dead cells and the cocktail of antibodies (Supp. Table 14 - “sc-sorting” panel) for 30 minutes at 4°C. Cells were centrifuged for 5 minutes at 300 g, 4°C, resuspended in a final volume of 500 μl of 5% FBS in PBS, subsequently filtered into polypropylene FACS tubes (ThermoFisher, cat#: 352063) and sorted on a BD FACSAria Fusion.

### scRNA sequencing

Each cell suspension was submitted for 3’ single cell RNA sequencing using Single Cell G Chip Kit, chemistry v3.1 (10x Genomics Pleasanton, CA, USA), following the manufacturer’s instructions. Libraries were sequenced on an Illumina NovaSeq S4 targeting 50,000 reads per cell, and mapped to the GRCh38 human reference genome using the Cell Ranger toolkit (version 3.0.0).

### Nuclei preparation and multiome sequencing

Nuclei preparation was performed following 10x genomics recommendations. Live, CD45+ sorted cells were centrifuged at 300g for 10’ at 4°C. Pellets were resuspended in 45ul chilled lysis buffer (10mM Tris-Hcl pH 7.4, 10mM NaCl, 3mM MgCl2, 0.1% Tween-20, 0.1% Nonidet P40 Substitute, 0.01% Digitonin, 1% BSA, 1mM DTT, 1U/ml RNase inhibitor in nuclease-free water), and incubated on ice for 5’. 50ul of chilled wash buffer (10mM Tris-Hcl pH 7.4, 10mM NaCl, 3mM MgCl2, 1% BSA, 0.1% Tween-20, 1mM DTT, 1U/ml RNase inhibitor in nuclease-free water) were added and nuclei were centrifuged at 500g for 7’ at 4°C. After removing the supernatant, the nuclear pellets were washed with 45ul chilled diluted nuclei buffer (1x Nuclei buffer, 1mM DTT, 1U/ml RNase inhibitor in nuclease-free water) without pipetting. Nuclei were centrifuged at 500g for 10’ at 4°C. Nuclei were resuspended in 7ul of chilled diluted nuclei buffer and counted. 15,000 nuclei were targeted for library preparation. Each nuclei suspension was submitted for library preparation using Chromium Next GEM Chip J Single Cell Kit (10x Genomics Pleasanton, CA, USA), following the manufacturer’s instructions. Libraries were sequenced on an Illumina NovaSeq S4 targeting 50,000 reads per nucleus, and mapped to the GRCh38 human reference genome using the Cell Ranger Arc toolkit (version 1.0.1).

### Processing of tissues for 10x Visium spatial transcriptomics

Tissues were frozen in dry-ice cooled isopentane and stored in air-tight tissue cryovials at - 80°C. Prior to undertaking any spatial transcriptomics protocol, the tissues were embedded in an optimal cutting temperature compound (OCT) and tested for RNA quality with RNA integrity number (RIN). Tissues with RIN values >7 were cryosectioned in a pre-cooled cryostat at 10 μm thickness. Two consecutive sections were cryosectioned at 10 μm thickness in a pre-cooled cryostat and transferred to the four 6.5mm x 6.5mm capture areas of the gene expression slide. Slides were fixed in methanol for 30 minutes prior to staining with H&E and then imaged using the Nanozoomer slide scanner. The tissues underwent permeabilization for 6 minutes. Reverse transcription and second strand synthesis was performed on the slide with cDNA quantification using qRT-PCR using KAPA SYBR FAST-qPCR kit (KAPA Biosystems) and analysed on the QuantStudio (ThermoFisher). Following library construction, these were quantified and pooled at 2.25 nM concentration. Pooled libraries from each slide were sequenced on NovaSeq SP (Illumina) using 150 base pair paired-end dual-indexed set up to obtain a sequencing depth of ∼50,000 reads as per 10x Genomics recommendations.

### Single-cell *in vitro* culture

Single cell colony forming unit (sc-CFU) was performed on foetal HSCs, as previously described^92^. Single, live, Lin-, CD34+, CD38-, CD62L+, CD52+ cells isolated from the foetal liver of three different Ts21 (median 13 PCW) and disomic foetuses (median 12 PCW), were index-sorted into 96-well plates (Supp. Table 14 - “HSC sc-CFU” panel) containing StemSpan SFEM (Stemcell Technologies) supplemented with penicillin/streptomycin (10U/ml penicillin, 100ng/ml streptomycin, Sigma Aldrich), 2mM L-glutamine (Thermo Scientific), 20ng/ml G-CSF (Peprotech), 20ng/ml SCF (Peprotech), 20ng/ml Flt3-L (Peprotech), 50ng/ml TPO (Peprotech), 20ng/ml IL3 (Peprotech), 20ng/ml IL6 (Peprotech), 20ng/ml IL5 (Peprotech), 20ng/ml M-CSF (Peprotech), 20ng/ml GM-CSF (Peprotech), 20U/ml EPO (RnD). Cells were cultured for 15 days at 37°C at 5% CO2. At the end of the culture, colonies were assessed for their lineage output by the expression of CD41a (megakaryocytic-Mk), CD235a (erythroid- Ery), CD3/CD56 (lymphoid-Ly), and CD11b (myeloid-My) by flow cytometry using a BD LSR- Fortessa analyser (Supp. Table 14 - “sc-CFU lineage” panel). Colonies were considered positive for a lineage if > 30 cells were detected in the relative gate. The total number of cells in the colony was determined by Trypan blue exclusion using a Countess II cell counter (Thermo Fisher). To assess differences in the colony output between Ts21 and disomic, we performed a chi-square test using Ts21 as observed distribution and disomic as expected distribution.

To compare Ts21 Ery lineage output to disomic Ery lineage output in the largest colonies, we first subset the Ts21 colonies with a cell count higher than 95% of all disomic colonies (equivalent to 20% of all Ts21 colonies), and the top 20% size colonies in disomic. The output of multi-lineage colonies was binarised to the lineage with the highest number of cells in the relative gate. We then performed a binomial test with N=17 observed Ts21 Ery lineages, K=23 total Ts21 lineages, and p=0.5 (the proportion of the disomic lineages that are Ery).

### Foetal liver phenotyping by flow cytometry

Cells were rapidly thawed in complete RPMI media at 37°C, then centrifuged at 300g for 5 minutes and washed again with DPBS. Cells were then resuspended in DPBS and LIVE/DEAD blue was added at a 1:800 final concentration. Cells were incubated for 15 minutes in the dark at room temperature, then washed with DPBS. Cells were then stained for 30 minutes in the dark at room temperature with the antibody cocktail (Supp. Table 14, “phenotype” panel), in the presence of BD Horizon™ Brilliant Stain Buffer (final dilution 1:4) and Miltenyi FcR blocking reagent (final dilution 1:5) in a final volume of 200µl. Cells were washed with DPBS and immediately acquired on a Cytek Aurora (5 lasers setup). Data was analysed on FlowJo v10.8.2.

### MitoTracker and MitoSOX staining

Cells were rapidly thawed in complete RPMI media at 37°C, then centrifuged at 300g for 5 minutes and washed again with DPBS. Immediately before the incubation with the dyes, MitoTracker Green FM reagent was dissolved in DMSO, and MitoSOX green was dissolved in anhydrous N, N-Dimethylformamide at a concentration of 1mM. Cells were resuspended in 1 ml DPBS and incubated with MitoTracker green FM (final dilution 1:1000) or 2 µM MitoSOX green for 30 minutes at 37°C in the presence of 50 µM Verapamil (diluted from an aqueous 10 mM solution). Cells were then washed with DPBS and stained for 30 minutes in the dark on ice with the antibody cocktail (Supp. Table 14, “mito” panel) in the presence of BD Horizon™ Brilliant Stain Buffer (final dilution 1:4), Miltenyi FcR blocking reagent (final dilution 1:5) and 50µM Verapamil, in a final volume of 100µl. Cells were washed again in DPBS and immediately acquired on a Cytek Aurora (5 lasers setup). Data was analysed on FlowJo v10.8.2. The populations of interest (Lin+, CD38+, CD38-, HSCs) were exported as a fsc file and imported in R via *flowCore 2.2* to obtain the fluorescence data of each mitochondrial probe for each cell.

### MitoTracker and MitoSOX Data analysis

To test whether Ts21 cells have significantly different mitochondrial mass or mtROS from disomic cells, we fit a Gaussian generalized linear mixed model. At single-cell resolution, we transformed the MitoSOX and mitochondrial mass values using a rank inverse normal transformation and used the transformed values as the response variable. We accounted for age as a fixed effect and sample as a random intercept. We tested for the effect of disease status (a fixed effect) within our model and determined significance using FDR across the 8 fitted models (4 cell types by 2 response variables).

### Analysis of the scRNA sequencing data

For each CellRanger output corresponding to a specific technical and biological replicate, we identified low-quality cells or empty droplets by applying the *barcodeRanks* and *emptyDrops* functions using the R package *DropletUtils*^93^. We then merged all CellRanger outputs into a single Scanpy object^94^. Following per-sample droplets removal, quality control (QC) was applied based on three parameters: the total UMI count (lower-upper threshold [750, 110,000]), the number of detected genes (lower-upper threshold [250, 8500]), and the proportion of mitochondrial gene count per cell (an upper-bound of 20%). We further applied *Scrublet*^95^ to remove potential doublets.

Next, we subsetted to samples of the same organ and trisomy 21 status and merged into a single dataset (e.g. a dataset containing only trisomy 21 liver samples). We reasoned that Ts21 and disomic cell transcriptomes would be influenced heavily by an extra copy of chromosome 21 (e.g. 50% higher expression within Ts21 cells that would not be present in disomic cells). As a result, the highly variable genes will be impacted by disomic versus Ts21 differences and insufficiently capture the genes relevant to liver- or femur-residing cells. Therefore, we chose to create disomic liver, Ts21 liver, disomic femur , and Ts21 femur datasets to most accurately annotate the population of the individual cells in the data. Within each of the four merged datasets, we applied log-normalization, using the scaling factor 10,000 to correct for between-sample differences in library size, and calculated highly variable genes, using the Seurat implementation^96^. We performed Principal Component Analysis on the highly variable genes for dimensionality reduction, retaining the top 15 components using the Scree plot elbow rule. Data was batch-corrected using Harmony^16^ to account for additional technical variations arising between samples which are non-biological in origin.

We then performed an iterative clustering procedure to identify clusters in the single-cell data. Broadly, our iterative clustering procedure first finds initial clusters using the Leiden algorithm, next merges clusters from seemingly identical cell populations, and finally subclusters into further refined populations using K-means clustering. Thus, the iterative clustering allowed us to further refine initial clustering, such that initial clusters containing multiple cell types can be further split into lower-level cell types. This is particularly useful for broad cell types such as erythroid cells that were further split into early and late erythroid cells. Following the between-sample batch correction above, we computed a neighborhood graph using the UMAP approach implemented in Scanpy and subsequently clustered with the Leiden algorithm. For visualisation purposes, we used Uniform Manifold Approximation and Projection (UMAP) manifold embedding to capture the global features in two and three dimensions. We identified marker genes for each cluster by performing a Wilcoxon signed-rank test with Bonferroni correction, and we annotated clusters using these marker genes and canonical marker genes. We performed further clustering, by manually choosing clusters to subcluster using K-means clustering, merging clusters of the same cell type, and performing marker gene detection. With this approach we generated four separate annotated scRNA-seq datasets, together with associated marker genes, for the Ts21 (liver and femur) and disomic (liver and femur) datasets.

### Contrasting cell-type abundances between Ts21 and disomic samples

To compare cell-type abundances, we calculated the proportion of each major cell type group in each sample. We contrasted cell type proportions between developmental stage-matched Ts21 and disomic samples with the same sorting strategy using a Mann-Whitney U test. Finally, we corrected for multiple testing using FDR and assessed significance at FDR < 0.1.

### Ligand-Receptor analysis using CellPhoneDB

We inferred statistically significant L-Rs and their corresponding cell types using CellPhoneDB on a subsampled Ts21 liver dataset, such that the proportion of cells in the reduced sample recapitulated the proportion in the full Ts21 dataset and corresponded to the number of cells in the disomic dataset. We repeated the same analysis on the full disomic dataset (which now has an identical cell size). We kept any pairs that did not involve HLA or a protein complex, and kept only those that involved a single receptor. Among the significant L-Rs (*P* < 0.001), we selected ligands or receptors identified in HSCs/MPPs and used to communicate with vascular endothelial cells, and performed gene set enrichment analysis on those using EnrichR.

### Analysing differential trajectory of osteo-lineage

For the disomic and Ts21 femur, we computed PAGA graphs using all annotated stromal cells (with PAGA threshold=0.05). We also computed a force-directed diffusion graph using Pegasus^97^, and overlaid the Pegasus and PAGA outputs.

Next, we focused on two different cases of osteo-linage transitioning: (i) within CAR cells, LepR+ CAR cells, osteoprogenitors and osteoblasts, and (ii) within arterial endothelial cells, transitioning endothelial cells and osteoblasts. We computed pseudotime using Scanpy, and used the pseudotimes as input into CellRank’s PseudotimeKernel (without usage of RNA velocity information) to obtain Generalised Perron Cluster Cluster Analysis (GPCCA) estimators for identifying macrostates and computing transition probabilities among them. We set the terminal state number according to the shape of the force-directed graph. For case (i), we chose two states in disomic cells based on the observation that there are two clear branches splitting between CAR cells and osteoblasts; and we chose one state in Ts21 because we observed a single branch leading to osteoprogenitors. For case (ii), we chose two states considering osteoblasts as one end and some transitioning endothelial cells as the other. Next, we plotted the STREAM plot using the scVelo package^26^ to visualise the cell type transition matrix. Finally, we correlated gene expression with estimated absorption probabilities (Pearson correlation, as implemented in the CellRank package). We identified the positively or negatively correlated genes at a significance level of FDR < 0.05 separately in Ts21 and disomic femur. We checked the GO terms of the top 500 genes that were most positively/negatively correlated to absorption probabilities using clusterProfiler R package^98^.

### Performing differential expression analysis in scRNA-seq

Within each cell type, four distinct differential expression analyses were performed to identify DEGs due to disease status (Ts21 or disomic) or microenvironment (liver or femur).

1. Liver *versus* femur in disomic samples
2. Liver *versus* femur in Ts21 samples
3. Ts21 *versus* disomic in liver samples
4. Ts21 *versus* disomic in femur samples

Prior literature has shown that pseudobulk differential expression methods have improved false discovery rates compared to single-cell differential expression methods^99^. As a result, our analyses were performed by first computing cell-type-specific pseudobulk profiles for each sample and then analyzing pseudobulk RNA-seq profiles using limma^100^.

To calculate sample-level pseudobulk profiles, we aggregated the read counts across cells of the same type. We kept samples for analysis that contained at least 10 cells, and we used the filterByExpr() function in the edgeR package with default settings to retain genes for differential expression analysis and reduce the burden of multiple test correction, by removing genes with low expression across samples^101^.

Next, limma-voom was used to perform a statistical analysis for differential expression. Briefly, sample-level weights were calculated by computing normalization factors for transforming count data into log2-counts per million and deriving weights based on a mean-variance relationship (using edgeR’s calcNormFactors() and limma’s voom() functions in R). Log-fold changes for each gene were estimated using a linear model with sorting strategy as a covariate. P-values were estimated after empirical Bayes shrinkage (limma’s lmFit and eBayes() functions). A Benjamini-Hochberg false discovery rate (FDR) correction was applied across all gene P-values, and significance was assessed at FDR < 0.05.

### Analysis of Ts21 versus disomic differentially expressed genes

As we observed an exponential cross-dependency between the proportion of differentially expressed genes on chromosome 21 and other chromosomes, we investigated additional factors that could be relevant. We first tested whether cell-type-specific overexpression of a particular gene on chromosome 21 can lead to greater dysregulation on either chromosome 21 or on other chromosomes. Since the probability of a gene being differentially expressed is linked to the number of cells tested in the DE analysis, we tested this using log-fold change values. For each chr21 gene and across cell types, we tested the Pearson correlation between log2-fold change of gene expression (from Ts21 vs disomic samples) and (i) chr21 DEG % or (ii) non-chr21 DEG % and determined the significance of the correlation using FDR < 0.1. Second, we reasoned that overall expression of an important chr21 gene could lead to greater dysregulation, as a highly-expressed chromatin modifier or transcription factor might have a consistent 50% overexpression in Ts21 across cell types, but the overexpression might matter more in the cell types where the gene is expressed. We tested this for each chr21 gene. Across cell types, we tested the Pearson correlation between average cell-type-specific gene expression (from Ts21 and disomic samples) and (i) chr21 DEG % or (ii) non-chr21 DEG %.

### Identification of context-specific DEGs in HSC/MPPs

It is difficult to ascertain whether a gene is commonly or uniquely upregulated in single-cell data (for example, a gene upregulated in Ts21 liver HSCs compared to Ts21 femur HSCs, but not disomic liver HSCs compared to disomic femur HSCs). The presence of a differentially expressed gene in one cell type and the absence in another may be a result of differences in population size, and thus purely statistical. Thus, we used to identify Ts21-induced expression changes between Liver and Femur for HSCs (as we could not simply take the Ts21 differentially expressed genes as the Ts21 population was much larger).

Since there are sample and cell count differences between datasets, we could not directly take the Ts21 liver versus femur differentially expressed genes as the Ts21 population was much larger than the disomic datasets. Instead, we identified DEGs specific to disease status (in liver *versus* femur analyses) and microenvironment (in Ts21 vs disomic analyses) in HSC/MPPs by using a subsampling procedure. Downsampling allows the ability to compare two analyses from distinct datasets that are confounded by differences in size. We did not repeat the same procedure across additional cell populations to conclude whether genes are differentially expressed or not in specific cell populations, as this would require to downsample to the smallest population sizes. This would erode statistical power and be computationally expensive.

Within liver *versus* femur analyses, we downsampled the Ts21 liver and Ts21 femur dataset to have the same number of foetuses contributing the same number of samples with the same number of HSC/MPPs as the disomic liver and disomic femur data. As a result, the Ts21 data matched the disomic data in terms of foetus-sample-cell counts. Similarly, within Ts21 *versus* disomic analyses, we downsampled the Ts21 and disomic liver data based on foetus-sample- cell counts in the Ts21 and disomic femur data. As an additional restriction in our downsampling, we ensured that foetuses present in both liver and femur data, with equal or greater number of cells and samples in the liver compared to femur, would still be selected in the downsample. The downsampling routine was repeated 100 times, such that 100 new datasets were created that match the smaller dataset. Differential expression analysis was performed identically to the full data using sample-level pseudobulks and limma-voom. The median nominal P-value for each DEG was calculated across 100 iterations. We verified the robustness of this choice of 100 iterations by visualizing the variability of the median P-value across iterations, in order to assess its stability.

Next, we used differential expression analyses in the full data and in the downsampled data to categorise the context-dependence of DEGs. In the liver *versus* femur analysis, we implicate *Environment-driven* DEGs, *Ts21-induced* DEGs, and *Ts21-reverted* DEGs.

a. *Environment-driven* DEGs are genes with an adjusted FDR < 0.05 in either Ts21 or disomic samples, and nominal P-value < 0.05 in the other dataset.
b. *Ts21-induced* DEGs were discovered by identifying genes with adjusted FDR < 0.05 and median p-value across 100 subsamples of P < 0.05 in the Ts21 data, and a nominal P-value > 0.05 in the disomic data.
c. *Ts21-reverted* DEGs are those discovered in only the disomic dataset: an adjusted FDR < 0.05 in disomic samples but nominal P > 0.05 in the Ts21 dataset.
d. *Environment-driven or Ts21-induced* DEGs are those genes that have FDR < 0.05 in the full Ts21 dataset, but have a median nominal p-value across 100 Ts21 subsamples of P > 0.05 and FDR > 0.05 in the disomic data. For these genes, we do not have sufficient evidence to claim context-dependence nor reject context-dependence, as the discovery of these DEGs might be highly dependent on sample size.

To visualize gene-environment interactions, we examined the expression of *Environment-driven* DEGs and *Ts21-induced* DEGs across Ts21 and disomic liver and femur HSC/MPPs. We scaled expression across cells for each DEG to mean = 0 and variance = 1. We averaged the scaled expression across genes within the *Environment-driven* and *Ts21-induced* gene sets, such that each cell has its own value for each gene set. We visualised the mean and standard error of these values across cells in Fig. 2d.

Gene-set enrichment analysis of upregulated Ts21-induced genes was performed by inputting the list of genes into EnrichR. Scatterplots show the top Gene Ontology terms (GO:MF - Molecular Function, GO:BP - Biological Process) or ENCODE and ChEA TFs.

### Identifying differences in cell-cycling across cell types, environment, and Ts21

We assigned a cell cycle using Scanpy’s *score_genes_cell_cycle()* function with the standard Tirosh et al.^102^ list of cycling genes, as applied to all cells from samples of the same environment and disease type. We determined cycling by the predicted cycling phase being equal to “G1” or not (either “G2M” or “S”). We compared the proportions of cycling cells using a Mann-Whitney U test.

### Evaluating the regional distribution of catalogued somatic mutations

Using publicly available data, we evaluated the hypothesis that Down’s syndrome impacts the regulatory landscape to influence where somatic mutations occur. First, we downloaded somatic mutation data from Hasaart et al. in foetal Ts21 and disomic HSCs^10^, and converted the mutation positions from hg19 to hg38 using liftOver. Second, we identified genes expressed in Ts21 HSCs. We used the filtered set from the Ts21 cycling versus less-cycling HSC differential expression analyses, which were identified by the filterByExpr() function. Next, we narrowed down the Ts21 and disomic sets of mutations within the HSC-expressed genes to the set of non-coding intronic mutations. Finally, we downloaded cCREs in hg38 from ENCODE, and for Ts21 and disomic mutation sets, we calculated the proportion of intronic somatics in Ts21 HSC-expressed genes that overlap with ENCODE cCREs. To determine significant differences, we bootstrapped the disomic intronic mutation set for 1000 times, and compared the observed Ts21 proportion to the disomic distribution. We calculated P-values as the proportion of bootstrapped disomic mutation sets with larger values than the Ts21 value.

### Analysis of the 10x Visium data

For disomic and Ts21 liver datasets, we used our annotated cell types from the disomic and Ts21 large scRNA-seq datasets as our input reference data for Cell2location^19^. Next, we merged all SpaceRanger outputs of tissue sections for disomic liver and then separately for Ts21 liver to create two Scanpy objects^96^. We removed mitochondrial genes and spots with the total expressed gene count less than 800 (the remaining spots were of sufficient good quality for downstream analysis).

We then estimated cell type abundances for each spatial spot. Using Cell2location, we trained a negative binomial regression model on the input reference data. We applied our model to the Scanpy formatted data, considering tissue section as a covariate to account for distinct batches. We used the estimated posterior mean value of each cell type (from Cell2location) as the local abundances. For each section, we computed the section-level relative abundance of each cell type as the proportion of its estimated abundance across all spots over the total estimated abundance of all cell types across all spots. We compared relative abundances between disomic and Ts21 using a Wilcoxon Rank Sum Test, and corrected p-values by using Benjamini-Hochberg.

To evaluate cell type colocalisation, we computed spot-level relative abundance of each cell type, dividing each cell type’s abundance on an individual spot by the total abundance of all cell types on the same spot. We then computed a Pearson distance matrix among cell types, based on these spot-level relative abundances across sufficient-quality spots of tissue sections in the same disease status, respectively for Ts21 Liver and disomic Liver. We next did hierarchical clustering, with inter-cluster distance estimated by the Ward variance minimization algorithm.

### Processing the multiome data

We performed the initial processing of multiome data using *Seurat* and *Signac*. After cellranger processing, we identified high-quality multiome cells for downstream analysis if they satisfied the following criteria: >750 RNA UMIs, >250 expressed genes, <40% mitochondrial read fraction, TSS enrichment score > 3, and >1000 ATAC fragments in peaks.

We next identified and annotated transcriptionally distinct clusters within Ts21 and disomic samples using Seurat. We created Ts21-specific and disomic-specific expression matrices by merging the matrices across Ts21 or disomic samples respectively. Within the separate Ts21 and disomic expression datasets, we log-normalized with a scaling factor of 10000, identified 2000 highly variable genes, scaled and centered the data, performed PCA, and used Harmony with lambda = 1 to batch correct for sample-specific variation. We then constructed a k-nearest neighbours graph based on the euclidean distance in PCA space (using the first 30 components), and identified transcriptionally distinct clusters using the leiden algorithm. We nominated marker genes for each cluster by performing a Wilcoxon signed-rank test that compares cells within one cluster to all other cells. We performed further clustering by performing K-means clustering and we merged clusters of the same cell type. With this approach, we annotated each cell in two separate multiome datasets (Ts21 and disomic). The Harmony-corrected datasets were visualized in two dimensions using UMAP.

Our overarching process for creating the chromatin accessibility matrix was to call peaks within each sample before calling a final set of peaks from cell-type-specific ATAC profiles. We first called peaks within each sample using *macs2*, as implemented with default parameters in Signac’s *callPeaks*() function. We created a unified set of peaks across all samples by combining any intersecting peaks into a single peak, and removing the combined peaks that were less than 20 bp or more than 10 kb wide. This set of peaks was used to compute a cell x peaks matrix for each sample from the ATAC fragments file. Using all peaks present in at least 10 cells, we ran latent semantic indexing (a two-step procedure of first using term frequency-inverse document frequency (TF-IDF) normalisation and then singular value decomposition) to project the ATAC matrix into a reduced dimension representation. We performed batch correction over all samples using Harmony before constructing a k-nearest neighbours graph across the first 30 components (except omitting the first component, which correlates with sequencing depth). This graph was used to perform clustering using the leiden algorithm with resolution = 1, and within each cluster, a new set of peaks were called using macs2. This set of peaks was combined into a unified set of peaks, which was used to form the final cell x peaks matrix which contained all final Ts21 and disomic cells.

For downstream analyses involving a single combined dataset, the two datasets were merged into a single matrix. Log-normalization, scaling, PCA, Harmony batch correction, and UMAP were applied to the combined dataset.

### Myeloid Trajectory analysis of snRNA using CellRank

Processed snRNA data from multiome was subset to cells of myeloid lineage. Both disomic and Ts21 cohorts were downsampled to 15,000 cells. A trajectory graph was calculated using Pegasus’ force directed layout (FLE) function. Instead of UMAP space, the cell trajectories were plotted in FLE space. Trajectory analysis of the RNA expression data closely followed the CellRank tutorial “CellRank beyond RNA velocity.” Moments of connectivity were calculated using scVelo with 30 principal components and 15 neighbours. The root cell was manually selected according to the diffusion map of HSCs, selecting the cell with the greatest Euclidean distance in FLE space from the centre of the cluster, indicating a cell with a divergent transcriptome. The pseudotimeKernel was used to calculate pseudotime with default parameters. The CytoTRACEKernel was used to compute the transition matrix with the parameters used in the tutorial (threshold_scheme=“soft”, nu=0.5). To compute terminal states and the probability of each cell differentiating toward each terminal state, the GPCCA estimator was utilised with default parameters. Schur decomposition was performed, and 5 terminal states were automatically selected according to an eigengap in the real part of the eigenvalues. Terminal states were labelled according to the cell type with the closest association (Late erythroid, Monocytes, Mast cells, pDCs, Megakaryocytes) and absorption probability was calculated.

### Topic modelling of HSC multiome using MIRA

Only HSCs were included for all downstream MIRA analysis. Out of the 6,215 HSCs, 3,784 were Ts21 and 2,431 were disomic. All MIRA analysis closely followed the online tutorials.

To generate the latent topics for RNA, the variational autoencoder (VAE) framework uses raw expression counts as input. Rare genes were removed by filtering genes only expressed in 15 or fewer cells. 7,905 exogenous genes were selected using Scanpy’s highly variable gene function, selecting for all genes with a minimum mean dispersion of 0.1. Exogenous genes are genes which will be captured in topics but will not be used as VAE features. 4,359 endogenous genes were selected by filtering the exogenous genes for those with a normalised dispersion greater than 0.5. Endogenous genes will be used as features for the VAE network. The ExpressionTopicModel was instantiated with the default parameters. The learning rate bounds were manually tuned to cover the portion of the learning rate vs loss curve with the steepest slope. The model was then tuned using TopicModelTuner with default iterations, a minimum number of topics set to 2, a maximum number of topics set to 15, a batch size of 32, 3-fold cross-validation, and a training size of 0.8.

To generate the latent topics for ATAC, the VAE framework uses binarized peak counts. Peaks were filtered according to the epiScanpy tutorial. Using the same process for RNA, 72,541 exogenous peaks and 45,095 endogenous peaks were selected according to a minimum mean dispersion of 0.05 and a normalised dispersion greater than 0.5. The AccessibilityTopicModel was instantiated with the default parameters and “dataset_loader_workers” set to 3. The learning rate bounds were manually tuned to cover the portion of the learning rate vs loss curve with the steepest slope. The model was then tuned using TopicModelTuner with default iterations, a minimum number of topics set to 2, a maximum number of topics set to 15, a batch size of 8, 1-fold cross-validation, and a training size of 0.8.

Gene set enrichment analysis (GSEA) was performed on the expression topics using MIRA’s wrapper of enrichr and the top 200 genes associated with each topic. For the accessibility topics, transcription factor (TF) binding sites were annotated in peaks using MIRA’s motif scanning and the hg38 reference from the UCSC repository. Each peak was scanned according to the JASPAR 2020 vertebrate collection of TF binding motifs. TF’s that were not expressed in the RNA data were removed. For comparing RNA and ATAC topics, the joint data was split into a disomic and Ts21 dataset and MIRA’s “get_topic_cross_correlation” function was performed.

### Trajectory analysis of HSC multiome using MIRA

First, a joint embedding space was calculated using MIRA’s “make_joint_representation” function to combine both modalities. A neighbourhood graph and UMAP embedding was performed on the joint representation (15 neighbours and a minimum distance of 0.1). The datasets were then batch corrected using Harmony on the joint UMAP features. The neighbourhood graph and UMAP embedding were re-calculated on the harmony adjusted feature space.

We next calculated several different HSC branches. First, the diffusion map was calculated using Scanpy “diffmap” with default parameters and then normalised by MIRA to regularise distortions in magnitude of the eigenvectors. Schur decomposition was performed and the eigengap heuristic was used to automatically select the proper number of diffusion components within the data (3). The data was subsetted to 3 diffusion components and the neighbourhood graph was calculated on the diffusion map embedding and components were connected. Pseudotime was calculated using MIRA’s “get_transport_map” function which defines a transport map using a Markov chain model of forward differentiation. A root cell was selected as the maximum value of the third diffusion component based on the suggestion in the tutorial. This root cell was located in the centre of the highest density of HSCs. Terminal cells were identified using MIRA’s “find_terminal_cells” function with 8 iterations and a threshold of 0.01. There were 3 distinct clusters of terminal cells, the cell in each cluster farthest away from the root cell was selected, and the probability of each cell differentiating toward those 3 branches was calculated. The lineage probabilities were parsed into a bifurcating tree structure using “get_tree_structure” with the threshold set to 1. Expression and accessibility dynamics across time were plotted using a streamgraph that depicts this tree structure.

### Regulatory potential modelling of HSC multiome using MIRA

MIRA’s LITE modelling was performed to link gene expression to nearby cis regulatory elements. For every gene, MIRA learns a regulatory window describing a range where changes in local accessibility appear to influence gene expression. The regulatory window decays exponentially both upstream and downstream according to the transcription start site (TSS) of each gene. Consequently, each gene is associated with a unique TSS using non-redundant human TSS annotations (hg38 GENCODE VM39). The model was trained on the union of genes that were highly variable and were the top 5% most activated in each of the 10 expression topics (n=5,367). The genes that did not have an annotated TSS were removed leaving 4,454 genes. The LITE model was instantiated with default parameters and the raw expression and accessibility data was then fit (4 out of 4,454 genes failed to fit). Now that each gene contained a trained regulatory potential model, the expression of each gene was estimated by calculating the maximum a posteriori prediction given the accessibility state of each gene in each cell.

Next, we identified TFs that regulate the expression of genes specific to each of the three HSC branches using MIRA’s probabilistic *in-silico* depletion method. MIRA simulates “computational knockouts” of each TF. MIRA uses regulatory potential (RP) modelling to predict gene expression based on local chromatin accessibility, and then masks cis-regulatory elements with specific motifs to define TFs where motif accessibility is important to gene expression prediction. This is measured by the changes in performance of the gene’s RP model to predict expression after computationally masking the binding sites of every TF. In this manner, TFs that strongly regulate a gene’s expression will be prioritised because masking of their binding site will significantly decrease the accuracy of the LITE model prediction. “probabilistic_isd” was run with default parameters across all modelled genes. In order to counteract the inefficiency of noisy TF binding site predictions, co-varying genes associated with individual topics were queried for a shared association across many TFs. The “driver_TF_test” function was utilised to identify potential TFs regulating the expression of branch specific topics. The top 150 topic specific genes were included, and a Wilcoxon rank sum test was performed over the association scores. After identifying TFs regulating topic expression, the top genes being regulated by these TFs were queried using “fetch_ISD_matrix” and selected according to the ranked association score. We additionally performed this procedure across all RNA topics, which we reported in the Supplementary Tables.

### Divergence in accessibility and expression of HSC multiome using MIRA

In order to identify genes where expression cannot be accurately predicted by local chromatin accessibility alone, MIRA NITE modelling was performed. In addition to cis regulation, the NITE model expands upon the scope of the LITE model by incorporating accessibility topics, which are genome wide. The NITE model was initialised using the same parameters, topics, and genes as the LITE model using “spawn_NITE_model.” The model was fit and expression was predicted using default parameters. The difference between LITE and NITE model performance for every gene was calculated using “get_chromatin_differential.” “get_NITE_score_genes” was used to calculate a cumulative metric per gene describing the divergence of local accessibility and expression across all cells. Genes with a high cumulative NITE score indicate genes that are regulated in part by non-local mechanisms. The top 500 genes with the highest NITE score were incorporated into MIRA’s GSEA analysis. To ascertain if Ts21 HSCs expressed genes enriched for non-local regulation, pseudobulk differential expression was performed (DESeq2) between disomic and Ts21 HSCs. The distribution of cumulative gene NITE scores for the top 300 condition specific genes (Log2FC) was compared using Wilcoxon rank sum test with Benjamini Hochberg correction.

### Assessing enrichment of GWAS SNPs in single cells using SCAVENGE

Trait-relevant individual cells were calculated using SCAVENGE, which combines network propagation and SNP enrichment analysis to map causal variants to their cellular context. First, variants with posterior inclusion probability (PIP) > 0.001 were downloaded in the format used by Yu et al.^63^, which were originally processed and described within Vuckovic et al.^62^ Second, the full multiome ATAC dataset (including all cell populations) was used as input to downstream tasks. A mutual *k* nearest neighbor graph was computed to represent the relationship between neighbouring single cells, using *k*=30. Next, g-chromVAR was used to calculate bias-corrected Z-scores for each tested trait and each single cell to estimate cell-trait relevance. The top 5% ranked cells served as seed cells for the SCAVENGE network propagation, which were further scaled and normalised to calculate the final SCAVENGE trait relevance score (TRS).

SCAVENGE TRS were contrasted between different lineage branches using a pseudobulk approach similar to the one we employed for differential expression analysis. On a per-sample-and-branch level, we pseudobulked the SCAVENGE scores by summing SCAVENGE scores across all cells of the same branch and of the same sample. We then applied three linear models to assess the impact of each of the three branches on SCAVENGE TRS. In each model, we accounted for the number of cells within the pseudobulk as a covariate. We determined significance by multiple test corrections at FDR < 0.1 across the 9 analyses (3 branches (1, 2, 3) and 3 traits (red blood cell, white blood cell, and lymphocyte counts)).

### Identification and analysis of peak-gene links using SCENT

SCENT^103^ was used to identify peak-gene links in disomic HSCs and Ts21 HSCs, by correlating peak accessibility (binarized) and gene expression (raw) counts. SCENT is a Poisson regression model that recomputes the standard errors in the model coefficients by using bootstrapping, which helps maintain false positive rates.

We first identified all peak-gene combinations within 500 kb from each other. Separately within disomic and Ts21 HSCs, we retained all peak-gene combinations where >5% cells were accessible and expressed the peak and the gene, retaining a total of 38,478 peak-gene combinations. Due to biological differences between disomic and Ts21, this was an overlapping but not identical peak-gene set tested in disomic and Ts21 HSCs.

Next, we applied a SCENT model using binarized peak accessibility, % mitochondrial reads, log(number of UMIs), and sample as the covariates, and expression counts as the dependent variable to each peak-gene combination. Based on the SCENT paper, we set up an iterative bootstrapping scheme to balance runtime and *P*-value accuracy based on the Poisson regression model *P*-values, where *P* > 0.1 consisted of 100 bootstraps, *P* < 0.1 consisted of 1,000 bootstraps, and *P* < 0.01 consisted of 10,000 bootstraps. SCENT was performed on a computing cluster using chunks of 100-500 peak-gene sets. We corrected for multiple testing using FDR, and determined significant peak-gene links at *FDR* < 0.2. Finally, we repeated the Ts21 analysis after downsampling the 3,784 Ts21 HSCs to match the sample size of 2,431 disomic HSCs to assess the impact of cell population size.

Further, we used SCENT to test whether the effect of peak accessibility on gene expression is modified by trisomy. To do so, we included an interaction term between ATAC peak accessibility and trisomy 21 status in the SCENT model to address whether the effect of accessibility on expression depends on trisomy. We applied the new SCENT interaction model to a combined Ts21 and disomic HSC dataset. The analysis was performed on all significant peak-gene links in the disomic-only or Ts21-only analyses, and significant interaction terms were assessed at FDR < 0.2.

One reason why Ts21-only peak-gene links were only identified in Ts21 yet have no significant interaction term would be a lack of gene expression or peak accessibility in disomic cells. To test whether Ts21-only peak-gene links without a significant interaction term were more accessible and expressed in Ts21 HSCs compared to disomic HSCs, we used differential accessibility (from the 10X multiome ATAC) and differential expression results (from the large scRNA-seq analysis). To calculate differential accessibility in the 10X multiome ATAC, we computed pseudobulk profiles and performed limma-voom with trisomy status as the covariate of interest, as similarly described in the large scRNA-seq analysis. We filtered peaks for differentially analysis using the filterByExpr() function in edgeR. We performed two separate binomial tests to assess whether the list of (a) Ts21-only peaks or (b) Ts21-only genes were upregulated in Ts21 compared to all other peaks or genes tested in differential analyses. We defined upregulated in Ts21 as nominal P < 0.05 & logFC > 0 in the limma-voom results.

We performed two enrichment tests with regard to our Ts21 peak-gene links and RBC GWAS. We defined SCENT peaks and SCENT genes according to three significance thresholds from the SCENT peak-gene analysis: FDR < 0.1, FDR < 0.2, and nominal P < 0.05, and identified GWAS-related peaks and genes using fine-mapped RBC GWAS SNPs at PIP > 0.2. In the first analysis, we assessed the enrichment of fine-mapped RBC GWAS SNPs (PIP > 0.2) within Ts21 SCENT peaks (also known as active cis-regulatory regions) using a Fisher’s exact test. Here, we are comparing whether peaks with a GWAS variant are more likely to be associated with gene expression compared to a background set of peaks without a GWAS variant, which include peaks accessible (in >5% cells) and close to a gene (< 500 kb) that is expressed (in >5% cells). In the second analysis, we assessed the enrichment of differential expression at target GWAS genes defined by Ts21 SCENT peak-gene links. We defined differential expression as FDR < 0.1 in the differential expression analysis of HSCs using the large scRNA-seq data. We subsetted to significant peak-gene links, and calculated a 2x2 contingency table reflecting (i) whether the peak contained a fine-mapped variant, and (ii) whether the target gene is differentially expressed. A Fisher’s exact test assessed the hypothesis that important enhancers (containing fine-mapped GWAS SNPs) are having a role in differential expression within HSCs (affecting their target genes).

Within the Results, we report the enrichment *P*-values within the text when using the set of peak-gene links with nominal *P* < 0.05 (from SCENT). Additionally, when reporting peak-gene examples within the Results, we use the nominal *P* values for the SCENT peak-gene test, the differential expression test, or the differentially accessibility test. We include the full summary statistics within the Supp. Tables.

## Acknowledgement

The human embryonic and foetal material was provided by the Joint MRC / Wellcome Trust (Grant # MR/006237/1) Human Developmental Biology Resource (http://www.hdbr.org). We thank the Cambridge NIHR BRC Cell Phenotyping Hub for their advice and support in cell sorting; D. Nachun for discussions regarding somatic mutations found in Ts21 and statistical modelling of mitochondrial mass data; J. Xu, C. Nicu, A.M. Ranzoni, B. Myers and E. Panada for sample collection and processing and M. Nelson for computational support with initial processing and clustering of scRNA-Seq and 10x Visium data; Cancer Research UK Cambridge Institute (CRUK CI) (Grant # CTRQQR-2021\100012) Genomics Core Facility for library preparation and sequencing services; Wellcome Sanger Institute (WSI) DNA pipelines for their contribution to sequencing the data; S. Leonard from New Pipeline Group (NPG) for pre-processing of sequencing data; R. Moller, P. Rainer, and E. Padhi for critical reading of the manuscript.

## Code and data availability

The code will be organised and released at: https://github.com/drewmard/t21-proj,https://github.com/drewmard/t21_multiome, and https://gitlab.com/cvejic-group/downsyndrome/.

The data has been uploaded to ArrayExpress and will be released upon publication.

## Author contributions

A.C. designed the study and oversaw all experiments. A.R.M., S.B.M. and A.C. led and oversaw all analyses. M.D.Z. performed experiments. S.W. performed experiments and created figures. H.X. led spatial transcriptomics analyses. J.B. led application of MIRA to multiome data. A.R.M., H.X., J.B. provided computational support. A.C. and A.R.M. wrote the manuscript, and all authors edited and reviewed the manuscript.

## Supplementary Results

### Cellular taxonomy of Down’s Syndrome and disomic foetal liver and bone marrow

Based on differential expression analysis and marker genes we manually annotated distinct blood populations in the foetal liver and BM. Within the hematopoietic progenitor compartment, we annotated clusters as HSC/MPPs (*CD34*, *SPINK2*, *MLLT3*), megakaryocyte-erythroid-mast progenitors (MEMPs - expressing *GATA1*, *KLF1*, *TESPA1*), granulocytic progenitors (GPs - expressing *MPO*, *AZU1*, *SPI1*, *LYZ*, *CD34*), as well as numerous mature blood cell types.

We identified clear transcriptional signatures of early and late erythroid cells (with graded expression of *AHSP*, *HBA1*, *ALAS2*, and *GYPA*), megakaryocytes (MKs - expressing *PLEK*, *ITGA2B*, and *GP9*), mast cells (*GATA2*, *CPA3* and *HDC*), monocyte precursors (*LYZ* and *SPI1*), plasmacytoid dendritic cells (pDCs - expressing *IL3RA*, *IRF8*, *CLEC4C*, and *JCHAIN*), and conventional DC2 (cDC2 - expressing *CD1C*, *CLEC4A*, *CLEC10A*), monocytes (*CD68, S100A9*, *MNDA*, *FCN1*), liver-specific Kupffer cells and femur-specific tolerogenic macrophages (expressing *CD163*, *MS4A7*, *CTSB*) and osteoclasts (*ACP5, MMP9, CTSK*).

In the lymphoid compartment, we identified NK progenitors and NK cells (expressing varying levels of *NKG7*, *PRF1*, and *GZMA*); *SPI1+* NK cells that were exclusively present in the Ts21 BM and had gene expression signature of both NK cells (*NKG7*, *PRF1*, and *GZMA)* and myeloid cells (*SPI1, CD14*). The B cell lineage included pre-pro B cells, which showed expression of *IL7R*; pro-B cells, expressing higher levels of B-cell genes *VPREB1, MME*, *PAX5, RAG1, RAG2* compared with pre-pro B progenitors and cluster of mature B cells expressing high levels of *CD79B, IGHM* and decreased levels of *IGLL1* compared with pre-pro/pro-B cell clusters. In addition, we identified clusters of highly cycling cells expressing lineage specific genes and proliferation markers e.g., *MKI67*.

### Cell populations in the foetal liver niche

Hepatocytes are the main and the most abundant parenchymal cell type in the liver. Cells in this cluster expressed well known markers of hepatocytes such as *ALB*, *AFP*^67^. Liver sinusoidal endothelial cells (LSECs) are the most abundant non-parenchymal cell type in the liver that form a barrier between the circulation, and the underlying hepatocytes and hepatic stellate cells within the space of Disse^68^. In line with this we identified a large cluster of LSEC cells that expressed typical endothelial markers (*KDR*, *CDH5*), lymphatic vessel endothelial hyaluronan receptor 1 *(LYVE1)*^69^ but also scavenger receptors such as stabilin 1 and 2 (*STAB1* and *STAB2*)^70^.

In addition, we identified non-LSEC endothelial cells i.e. arterial endothelial cells expressing *KDR*, *CDH5* but also calcitonin-receptor-like receptor (*CALCRL)* and *RAMP2* suggesting their sensitivity to adrenomedullin signalling^71^. RAMP2 is required to transport CALCRL to the plasma membrane where it acts as adrenomedullin receptor. LSECs maintain close interactions with hepatic stellate cells which are liver-specific mesenchymal cells.

We identified population of hepatic stellate cells based on the expression of a vitamin A-associated transcript *RBP1*^72^ as well as *DCN, COL1A1* and a cluster of activated hepatic stellate cells that expressed α smooth muscle actin (*ACTA2*), *TAGLN* and *MYL9*^67^.

### Cell populations in the bone marrow niche

In total we analysed 79,512 niche cells from Ts21 and 32,501 niche cells from disomic BM. We thereby identified, pericytes (*PDGFRB*, *TAGLN*, *ACTA2, RGS5, NOTCH3, MYH11, MCAM*), myofibroblasts (*PDGFRB, PDGFRA, TAGLN*, *ACTA2, NOTCH3)*, non-myelinating Schwann cells (*MPZ*, *NRXN1*)^73^, *PDGFRA-*positive mesenchymal population that expressed high level of *CXCL12* therefore representing CXCL12-abundant reticular (CAR) cells as well as LepR+ CAR cells^74^. LepR+ CAR expressed *LEPR, CXCL12, PDGFRA, PDGFRB* and low level of nestin (*NES),* but not key HSC niche factors such as stem cell factor (*KITLG)* and Angiopoietin-1 (*ANGPT1*). disomic BM had additional clusters of mesenchymal cells (annotated here as 1 and 2).

In the osteo-lineage, we identified two clusters, osteoprogenitors (present in both disomic and Ts21) and osteoblasts (present only in disomic). Osteoprogenitors expressed *RUNX2,* the master regulator transcription factor (TF) known to control the commitment of mesenchymal stem cells (MSCs) to osteo-lineage and osterix (*SP7*), its downstream target TF^75^. Other osteogenic genes included osteopontin (*SPP1*) and integrin-bone sialoprotein (*IBSP*) which together are required for early and late osteo-lineage differentiation^76^. Cells in osteoprogenitor cluster co-expressed *FOXC1* which is essential TF for development of adipo-osteogenic progenitors and which up-regulates *CXCL12* and inhibits adipogenic processes in CAR cell progenitors^77^; *SOX9*, a TF essential for chondrocyte differentiation^78^ and, aggrecan *(ACAN*) a chondrocyte marker gene^79^. Immature osteoblasts (present only in disomic BM) further upregulated *SP7* and bone gamma-carboxyglutamate protein (*BGLAP),* an osteoblast maturation marker^80^. In contrast, in Ts21 a smaller proportion of cells up-regulated osteo- lineage related genes such as *IBSP, BGLAP* and *SP7* (Supp. Figure 4).

Finally, we identified three main endothelial cell (EC) clusters all expressing *CDH5* and *KDR*. The endothelial population included *STAB2*, *STAB1*, *LYVE1, TSPAN7 -* expressing liver sinusoidal ECs (LSECs), arteriolar endothelial cells (high level of *CD34, KITLG*^81^ and *CXCL12*), and transitioning endothelial cells that co-expressed endothelial as well as stromal markers (such as *DCN*, *COL1A1, PDGFRA)*. Specifically in disomic samples we found an additional cluster of immature endothelial cells which had a low expression of *CDH5*, *KDR and CD34*.

### Evaluating cell-type abundances following data-set integration and label transfer

Cell types exist on a continuum, and clear and distinct clusters on a UMAP are not expected during haematopoiesis. Thus, to test whether technical biases that arise from separate annotation will lead to confounding in differential cell-type abundance tests, we re-clustered our data using 1) integration (with both Harmony and scVi methods) and 2) reference-query mapping (for disomic samples with both Popescu et al. Nature 2019^18^ and our Ts21 data). Using the new clusters, we repeated the cell type abundance comparison.

First, we integrated the full dataset using Harmony (Supp. Figure 7a, b) using 5000 highly variable genes to construct the PCA space. We estimated 107 principal components (PCs) as the optimal number of PCs based on random matrix theory (implemented in pegasus’ pegasus.elbowplot function). Next, we applied 3 iterations of Harmony to integrate data across samples, correcting for sample/biological replicate, environment, and diseases. The output from Harmony was used as input for calculating the K-nearest neighbor graph and computing clusters with Leiden clustering (resolution = 3.5). We annotated the clusters based on their top marker genes. Overall, the Harmony integration-identified cell types expressed key marker genes across all the anticipated cell types (Supp. Figure 7 c), validating accurate annotation of cell types.

Next, we compared the Harmony integration-identified cell type abundance analysis to our separate cell type abundance analysis shown in Figure 1c. The results were remarkably consistent between the two approaches (Supp. Figure 7 d, e). The only difference being that, while the trend remained consistent, mast cells were not anymore significantly differentially abundant (FDR = 0.103).

Given that benchmarking studies^82^ have shown that Harmony is not the preferred integration method for complex integration tasks such as our dataset, which combines two environments (liver and femur) with two highly different genetic backgrounds (disomic and trisomic), we repeated the integration and clustering using scVI (Supp. Figure 8). We ran scVI on the raw counts for the top 5000 highly variable genes with sample, environment, and disease used as batch covariates and a *max_epochs=400* (as the default falls to *max_epochs=*7 when there are 1.1 million cells such as our dataset). The embedding space from scVI was used as input for calculating the K-nearest neighbor graph, computing Leiden clusters, and finally annotating clusters.

The scVI-integrated proportions across samples correlated closely with separate dataset proportions (*r* = 0.957) (Supp. Figure 8 d), a slight improvement compared to the Harmony-based integration (*r* = 0.949). Furthermore, the statistical results from our differential cell abundance analysis confirmed all results from the separate analysis, including a significant increase in mast cells found in Ts21 (Supp. Figure 8e).

Next, we performed reference query mapping from Popescu et al. 2019 Nature to our disomic liver dataset. This allowed us to compare the cell type abundances based on annotations from a published dataset to what we observed in both our original separate-dataset annotations. First, we performed the reference query mapping using scArches. To improve the reference query mapping accuracy and rejection rate, we considered reference mapping for only the 45,497 haematopoietic cells identified through our separate annotations for disomic liver. We combined all scRNA datasets into a single matrix and identified the top 5000 highly variable genes. We used scVI to embed both datasets into the same latent dimension using a variational autoencoder. Finally, we predicted cell-type labels using a KNN classifier (scHPL) with default settings (50 neighbors for knn, a rejection threshold of 0.5, and a match threshold of 0.25). We mapped the reference query cell type labels to the broad cell type groups and compared the group abundances to those found using the separate-dataset annotation labels. Given that the Ts21 liver cells were vastly different (due to presence of trisomy) and much larger than the Popescu et al. dataset, we did not transfer labels for the Ts21 data for this analysis and perform a cell-type abundance test.

Overall, in the three disomic samples that were tested for cell-type abundance differences, we found that the new reference query labels correlated strongly with the previous annotations, helping validate our prior annotations (*r*=0.998; Supp Figure 9 a).

Next, we performed a reference query mapping from Ts21 liver to disomic liver and compared the Ts21-transferred annotations to our separate-dataset annotations. Compared to the original separate-dataset annotations, we found that the estimated cell-type abundances were highly correlated with abundances from the Ts21-based reference query mapping in the three stage-matched disomic liver samples (r=0.998; Supp. Figure 9a). Furthermore, we repeated the Ts21 *versus* disomic cell abundance analysis using the original Ts21 liver annotations and Ts21-label transfer for disomic liver annotations, and replicated our original separate-dataset differential abundance results. Overall, our results demonstrate that the cell-type annotations are consistent across all annotation strategies and the differential abundance results are robust.

Finally, we projected our disomic BM dataset onto the Jardine et al. data set^65^. Since we mostly used CD235 depletion strategy, late erythroid cells were largely depleted from our samples as evident from the UMAP (Supp. Figure 9b). In this analysis, we filtered our disomic BM to only include blood cells, and then embedded our blood cell-only disomic BM and Jardine et al. disomic BM into a single latent representation using scVI integration. Considering that cells from these two studies were collected from very different developmental stages and sorting strategies, our cells mapped to similar cell types in the Jardine et al. data, reflecting that similar cell populations were captured in both studies.

### Evaluating cross-dependent dysregulation of chr21 genes versus other genes

Our analysis suggests that when greater than 50% of genes on chromosome 21 are dysregulated in a cell type, there is dysregulation of more than 10% of genes on the other chromosomes within an exponential non-linear relationship. This trend was observed in both femur and liver. Since the cell types that show high dysregulation of chr21 genes are different in the two environments, our analysis demonstrates that the environment plays a role in driving this phenomena, however additional factors might also be relevant.

To explore this further, we first tested whether the overexpression of specific genes on chromosome 21 significantly correlates with the overall extent of dysregulation on both chr21 or other chromosomes. For each chr21 gene and for all cell types, we tested the Pearson correlation between the log2-fold change of gene expression (Ts21 vs disomic samples) and the % of DEG on (i) chr21 or (ii) non-chr21. After FDR correction, we did not find any significant correlation.

Next, we considered the alternative hypothesis that the expression level of genes on chr21 could influence their dysregulating effect. The rationale behind this idea is that important chromatin modifiers or TFs might consistently exhibit 50% overexpression in Ts21 cells, yet their impact would become critical only when their baseline expression is naturally sufficiently high. We tested this hypothesis for each chr21 gene and each cell type. We tested the Pearson correlation between the average cell-type-specific gene expression (from Ts21 or disomic samples) and the % of DEG on (i) chr21 or (ii) non-chr21. After FDR correction, we did not find any significant correlation.

Therefore, with our data in these analyses, we were unable to identify individual genes (such as particular chromatin modifiers or transcription factors) that drive genome-wide dysregulation of gene expression. We speculate that alternative hypotheses that we were unable to address include the existence of feedback loops that modulate chr21 gene expression (which would hinder analyses from a single time point) or synergistic regulation (where overexpressed genes lying in close proximity synergistically contribute exponential regulatory effects to other genes).

### Impact of chromosome 21 on differential expression analysis

We developed a simulation framework to carefully consider the statistical impact of the additional chromosome 21 on the differential expression analysis (DEA) results, specifically for genes not on chromosome 21. To do so, we simulated trisomy-like samples from disomic samples by increasing the reads counts of all genes on chromosome 21 by 50%. For example, if there are 100 reads counted for the chromosome 21 gene ABCG1 in a pseudobulk HSC sample, we increased the reads by 50% such that there are 150 ABCG1 reads in total (which was done for all chromosome 21 genes). We then assessed how the differential expression results of disomic samples were affected by this manipulation.

First, we performed DEA between the HSCs (pseudobulk) of two disomic foetuses (foetus #15633 and #15781). We then simulated a trisomy-like sample from one of the foetuses and ran DEA again. Finally, we compared the results by computing the median log fold change for genes on each chromosome (Supp Figure 12c), and comparing log fold change between the two analyses for all genes (Supp Figure 12 d).

Overall, we found that, in the real data, the median log fold change on each chromosome was roughly centered around zero (Supp. Figure 12c, left). When looking at the chromosome 21 spiked analysis, there was roughly a 50% increase on chromosome 21, while the median log-fold changes on other chromosomes were similar to the real data (Supp. Figure 12c, right). As a further check of the impact of the chromosome 21 dosage effect on analysis, we found that the estimated log-fold change for each gene was nearly identical between both analyses, except for chromosome 21 where there is an induced increase in expression (Supp. Figure 12d). Therefore, the increased expression on chromosome 21 has minimal impact on the differential expression results on other chromosomes.

### Additional signalling is necessary to enact transcription of erythroid lineage related genes in Ts21 HSCs

We next asked if the expression of genes upregulated in each of the HSCs branches were best predicted by an RP model based on local (cis) accessibility alone (Local chromatin accessibility-Influenced Transcriptional Expression (LITE) model) or if their expression is better predicted by a model additionally considering genome-wide accessibility (Non-local chromatin accessibility-Influenced Transcriptional Expression (NITE) model). To address this question, we used MIRA to compute LITE and NITE models for each gene and then a likelihood ratio test to contrast the two models. We found that the top 300 genes upregulated in Ts21 HSCs had a much higher NITE score than those downregulated in Ts21 HSCs (Supp. Figure 16a, b), indicating that expression of Ts21-specific genes cannot be fully modelled by local chromatin accessibility and that additional factors, such as additional signalling, are necessary to enact transcription of these genes. The genes with the highest NITE score were mainly expressed in branch 2 (Supp. Figure 16c). One such example is *TFR2,* which is required for efficient development of red blood cells^64,83^. Based on the observed *cis* accessibility, *TFR2* was predicted to be expressed in both branch 2 and 3, but it was only expressed in branch 2, causing an overestimation of its expression in branch 3 by the LITE model (Supp. Figure 16d). Its expression specifically in branch 2, despite much broader accessibility, suggests that *TFR2* expression in Ts21 HSCs requires additional signalling that does not primarily affect transcription via remodelling local accessibility. Genes with the high NITE score were associated with “interleukin 2 signalling pathway”, which plays an important role in erythropoiesis^84^, and “myelodysplastic syndrome” (Supp. Figure 16e). This is in line with the model in which subpopulation of Ts21 HSCs attained a “primed” chromatin state which in response to signalling (i.e. additional factor(s)) activated the expression of erythroid lineage related genes.

### Enrichment of leukaemia GWAS SNPs in multiome peaks

To address whether leukaemia GWAS SNPs showed accessibility differences in Ts21, we used the largest published childhood acute lymphoblastic leukaemia (ALL) GWAS study^85^ and performed SCAVENGE analysis similar to the analysis we presented for RBC, WBC, and lymphocyte count. We found a significant enrichment in B cell populations (Supp. Figure 17), which confirms the utility of the SCAVENGE approach, but no significant difference was observed between Ts21 and disomic HSCs or B cells (Mann-Whitney U test P-value > 0.05). This might be the case because any potential differences that might exist are currently undetectable due to the limited number of identified GWAS signals associated with childhood ALL risk.

### Evaluating the robustness of differential expression results to analysis decisions

Our main analysis did not account for sex and age or any genetic markers as covariates in analysis. We only pseudobulked cells on a per-sample level. Considering that analysis decisions such as sample-level pseudobulking or omission of covariates have the possibility of obscuring true differences, we first aimed to evaluate the robustness of our differential expression analysis results to these decisions. In all analyses, we ignored individual genetic polymorphisms and genetic ancestry, as we did not have access to individual genotype data. Furthermore, common regulatory variants should have similar frequency between trisomy 21 samples and disomic samples, as germline trisomy 21 is approximately independent from genome-wide SNP frequencies. Therefore, it is not expected that regulatory variants which majorly impact expression would cluster within Ts21 cases or disomic controls; ultimately, impacting statistical results as a confounder that hasn’t been accounted for. We conducted the HSC Liver *versus* Femur differential expression analysis using four different statistical approaches for both disomic and Ts21 datasets (Supp. Figure 19).

1. Sample-level pseudobulks with the addition of sorting accounted for as a fixed effect (the original analysis in the manuscript)
2. Sample-level pseudobulks with the addition of sorting and age (PCW) accounted for as fixed effects
3. Foetus-level pseudobulks with no other covariates besides Liver vs Femur
4. Foetus-level pseudobulks with the addition of age (PCW) accounted for as a fixed effect

We additionally repeated analyses 2 and 4 using sex instead of age.

For age, we did not find any impact of these analysis decisions on the differential expression results, as the correlation between log-fold changes across all analyses had a Pearson correlation *r* > 0.98. Therefore, our differential expression results are consistent regardless of pseudobulking on sample or foetus. Furthermore, PCW had little to no impact on differential expression results (correlation between log-fold changes for all genes before and after including age as a covariate: r=0.999 in the disomic cells and *r*=1 in the Ts21 cells), (Supp. Figure 19 a). Thus, we found our results to be statistically robust regardless of these analysis choices.

Next, we repeated the above analysis except for sex as a covariate in place of age. Using the correlation of log-fold changes across genes as a metric of the sensitivity of our results, we found very high correlations between adjusting for sex and not adjusting for sex (*r* > 0.96 for disomic HSCs and *r* > 0.99 for Ts21 HSCs). Furthermore, 99.1% of genes that were significantly differentially expressed (*FDR* < 0.1) in the original Ts21 HSC analysis were recovered after accounting for sex as a covariate (*FDR* < 0.1), Supp. Figure 19 b. Therefore, sex did not have a large impact on differential expression results.

We additionally conducted an exploratory analysis of developmental age association with cell-type composition (Supp. Figure 19 c). We first estimated cell type composition for the 8 major cell types detailed in Figure 1c across disomic and Ts21 samples. Next, we performed: (1) a correlation test between PCW and abundance, and (2) a non-parametric test (Kruskal-Wallis) to discern differences between PCW groups. For both tests, the 8 major cell types were not significantly associated with cell composition (nominal P > 0.05 for all tests). We visualised the association between cell type group and age, and saw potential outliers existing in PCW 12. Thus, we performed a Mann Whitney U test of PCW 12 abundances versus PCW 13 + 14 abundances. Again, we found no significant association (nominal P > 0.05 for all tests). As a result and in addition to the identical differential expression analysis results above, we did not consider PCW further in cell abundance results.

Furthermore, we note that the observed biological differences in Liver vs Femur are supported by similar differential expression analysis results in the Ts21 and disomic datasets. We compared the correlation for all genes between the log-fold changes in the Ts21 dataset and in the disomic dataset (Supp. Figure 19 d). Overall, while the correlation within a single dataset was *r* > 0.98, the correlation between the two datasets ranged from 0.87 < *r* < 0.93. This high correlation reflects that not only are our results robust to statistical decisions, our results pick up on consistent biological differences between Liver and Femur environments that are present regardless of trisomy. Furthermore, the lower correlation reflects inherent biological differences that exist in Ts21 that do not exist in the disomic dataset, and vice versa. This motivates the analysis related to Figure 2b in the original manuscript, since there is a need to distinguish purely environment-driven expression differences from Ts21-induced expression differences. Finally, we found that these high correlations are not expected given random chance; for example, the correlation between log-fold changes from a Liver vs Femur analysis in NK cells versus the analyses above HSCs/MPPs was minimal, with a Pearson correlation *r* = 0.03 (*P* = 7.7 x 10^-6^) when compared to the original analysis from the manuscript.

Finally, we assessed the impact of integration and cell type annotation on our differential expression analysis. We conducted the HSC Liver *versus* Femur differential expression analysis again using cluster annotations from a Harmony-based integration across all datasets. We observed highly similar results between the new annotations and the original analysis, with *r* = 0.95 in the disomic dataset and *r* = 0.96 in the Ts21 dataset when comparing log-fold changes across genes between the two analyses. This indicates the robustness of our findings, as differential expression results were consistent across the various annotation processes.

### Comparison of cell-type frequencies from 10X multiome versus scRNA-seq

We observed similar frequencies of cell types in the multiome and scRNA-seq data, while acknowledging that the frequency of cell types in the multiome does not fully replicate the results observed in the scRNA-Seq experiment (Supp. Figure 20). There are a few rational explanations for this:

1. The multiome sample size was smaller than for scRNA-Seq, both in terms of the number of cells sequenced and the number of biological replicates, leading to lower statistical power.
2. The samples used in the scRNA-Seq and multiome experiments were partially matching - two out of three foetal Ts21 samples were used for both experiments, and none of the disomic samples used in multiome were used in scRNA-seq (Supp. Table 1, 2 and 8). This discrepancy in sample overlap, combined with relatively lower number of samples used for multiome, may potentially account for the observed differences in cell abundances between the two technologies.
3. Moreover, previous studies have demonstrated that different technologies are likely to produce differences in cell abundances. For example, a previous study by Massier et al.^86^ shows clearly that two technologies can sometimes recover cell types at different abundances. While abundances of three of the four major cell classes within Massier et al. were consistent between single-cell and single-nuclei technologies, vascular cells were found at nearly three-times higher abundance in single-nuclei data compared to single-cell data (Figure 1c from Massier et al.^86^).

### Detecting *GATA1* mutations in foetal samples using RNA-seq

Samples were not tested experimentally for *GATA1* mutation. However, mutations in *GATA1* occur at the later stages of foetal development compared to the ones used in this study and are not observed before 20 pcw^87^. In line with this, the larger screening studies in newborns have shown that the incidence of *GATA1* mutations is around 4%^88,89^. Furthermore, Haasart et al.^10^ did not observe a mutation in *GATA1* in any of the foetal stem and progenitor cells following WGS of HSPCs from 9 independent human foetuses gestational age week 12–17^10^.

While our Down syndrome samples have not been tested for GATA1 mutations, we set out to detect putative somatic variants from the single cell RNA sequencing data. There are, however, several limitations to this approach:

- The mutant copy of GATA1 needs to be expressed to be detected in single cell RNA sequencing.

- Sequencing reads need to span the sites of the mutations. Since we are using the 10x Chromium 3’ platform, the vast majority of reads will cover the 3’ UTR and terminal exons of GATA1 but will not extend to the earlier exons. The majority of GATA1 mutations that have been described in the Down syndrome context are truncating mutations in the first exons, towards the 5’ end of the gene.

- The RNA editing machinery can cause sequence variation without an underlying DNA mutation.

- The coverage of GATA1 per cell is very sparse.

These limitations notwithstanding, we used the variant calling framework based on deepSNV/Shearwater^90,91^ to detect variants in the GATA1 locus.

- Extract reads mapping to the GATA1 locus and generate a per position pileup of nucleotide counts. These counts are aggregated pseudo-bulks per sample.

- Estimate a site-specific error profile per position estimating from the single cell RNA sequencing data of the non-trisomy 21 samples.

- For every site, in every pseudobulk, use a likelihood ratio test to determine whether the observed variant depth and total depth is likely to come from the background error distribution. We correct for multiple hypothesis testing using the Benjamini-Hochberg approach.

With this approach, we detected seven variants in the GATA1 locus that passed the significance threshold of q<0.01. Of these, two variants are inherited mutations, consistent with high VAFs across all samples of the same individuals (chrX:48787974 G>A in TS21_6 and chrX:48791310 G>A in TS21_13). The remaining five variants are all confined to single sample and while likely somatic, are unlikely to impact the function of GATA1. Of the five variants, three are intronic, one is synonymous and one missense (G352A), which is predicted to be a benign variant. In addition, even though none of the variants listed here were previously identified as the consequences of RNA editing, we cannot definitively rule out the A>G and C>T may be caused by A>I and C>U RNA editing, respectively. Overall, none of the variants detected in the GATA1 locus from the single cell RNA sequencing data are likely to contribute to any preleukaemic phenotype.

### Supplementary Figure Legends

**Supplementary Figure 1.**
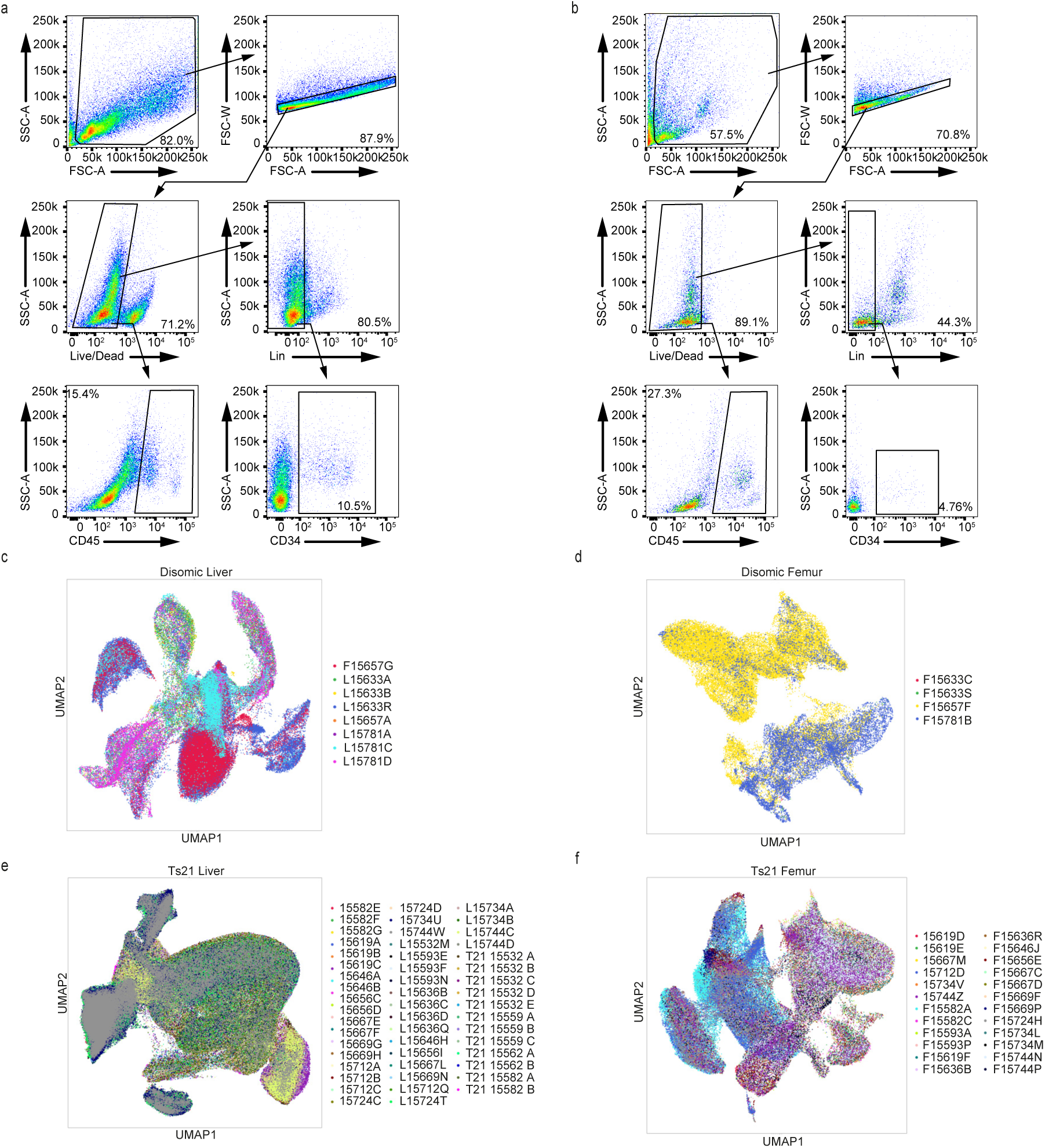
Sorting strategy and UMAP visualisation of batch effect corrected scRNA-Seq datasets. **a-b,** FACS sorting panel and gating strategy for the isolation of CD45+ and CD34+Lin- cells from foetal liver (**a**) and foetal BM (**b**). **c-f**, UMAP visualisation of the foetal scRNA-Seq samples after the batch effect correction with Harmony. Each colour represents a different sample. **c,** disomic liver (n=110,671, k=3, k*=8). **d**, disomic femur (n=53,807, k=3, k*=4). **e**, Ts21 liver (n=780,299, k=15, k*=50). **f**, Ts21 femur (n=162,775, k=12, k*=24). n refers to the total number of cells, k indicates the number of biologically independent samples, k* indicates the total number of replicates (both biological and technical).

**Supplementary Figure 2.**
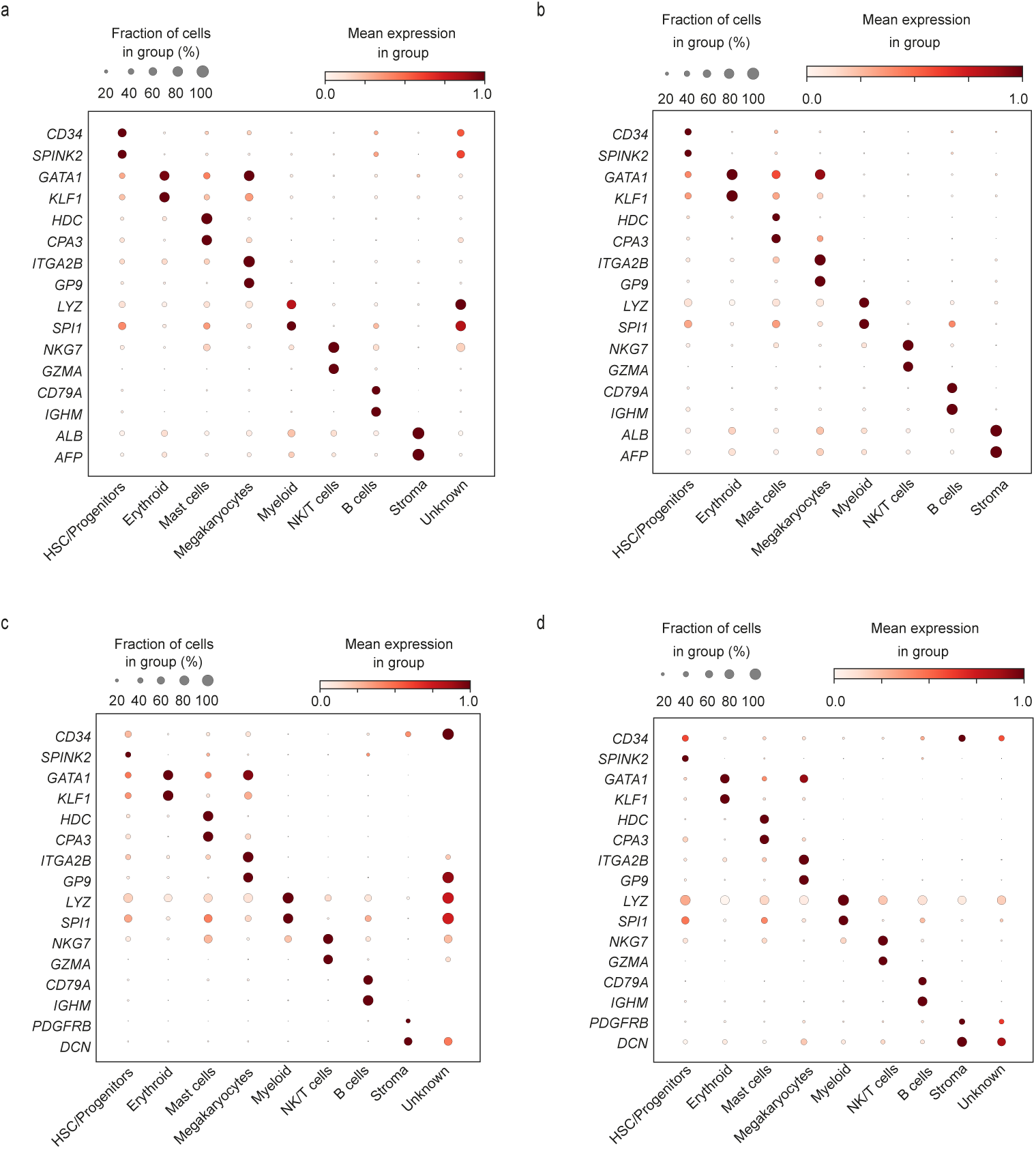
Single-cell transcriptome analysis of disomic foetal liver and bone marrow. a-d, Dot plot of the standardised gene expression of manually selected marker genes in annotated broad cell types in Ts21 foetal liver (**a**), Ts21 foetal BM (**b**), disomic foetal liver (**c**), and disomic foetal BM (**d)**. For each gene, the minimum value of its expression is subtracted and the result is divided by the maximum value of its expression. The dot size indicates the percentage of cells that express the gene of interest within each cell type.

**Supplementary Figure 3.**
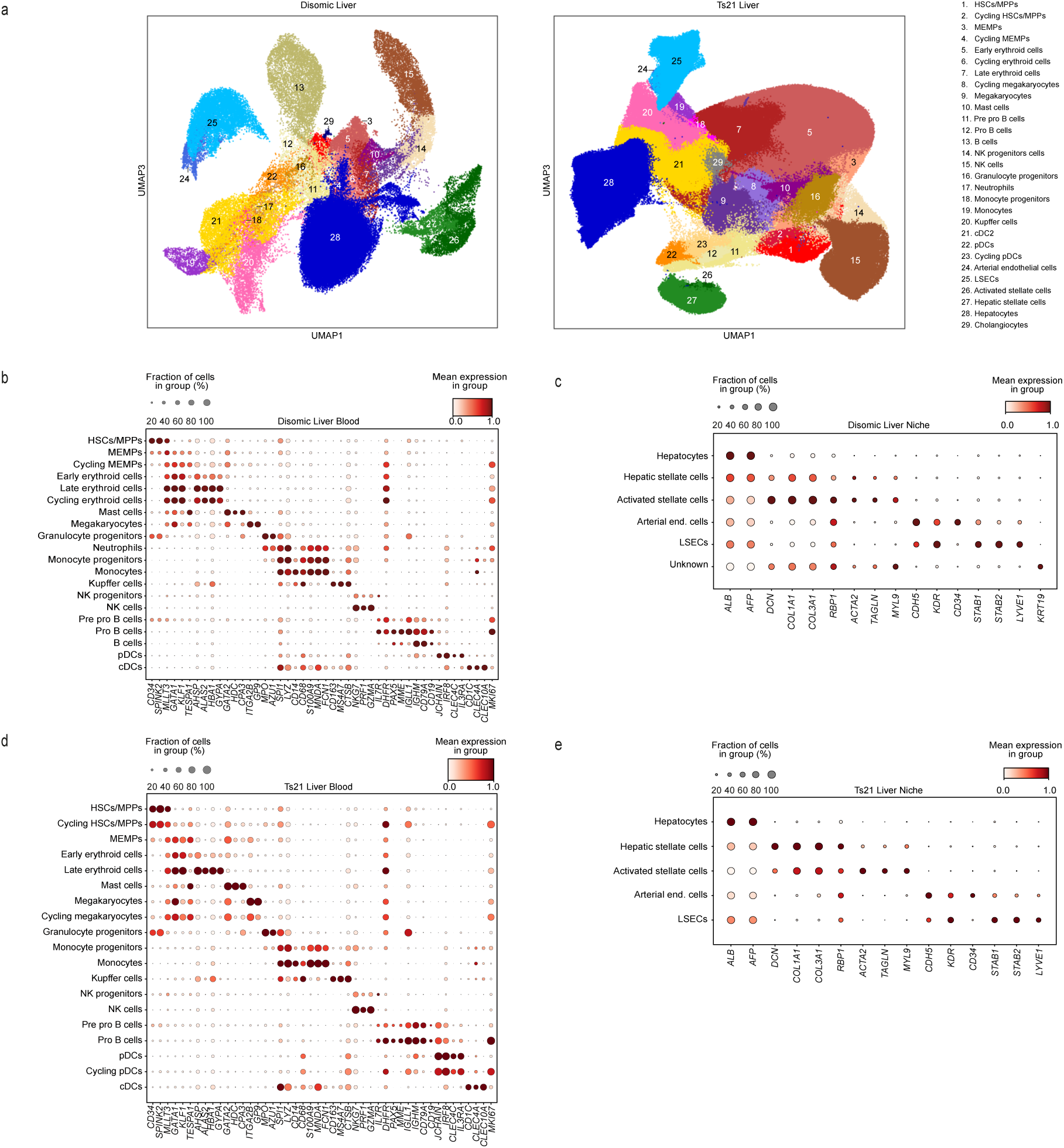
A single-cell atlas of disomic and Down’s Syndrome foetal liver. a. UMAP visualisation of single cells from disomic foetal liver (left; n=110,671, k=3) and Ts21 (right; n=780,299, k=15) coloured by cell type/state. **b-e,** Dot plot of the standardised gene expression of manually selected marker genes in the identified cell types. For each gene, the minimum value of its expression is subtracted and the result is divided by the maximum value of its expression. The dot size indicates the percentage of cells that express the gene of interest within each cell type. **b**, disomic liver haematopoietic cells. **c**, disomic liver niche cells. **d**, Ts21 liver haematopoietic cells. **e**, Ts21 liver niche cells. n refers to the total number of cells, k indicates the number of biologically independent samples.

**Supplementary Figure 4.**
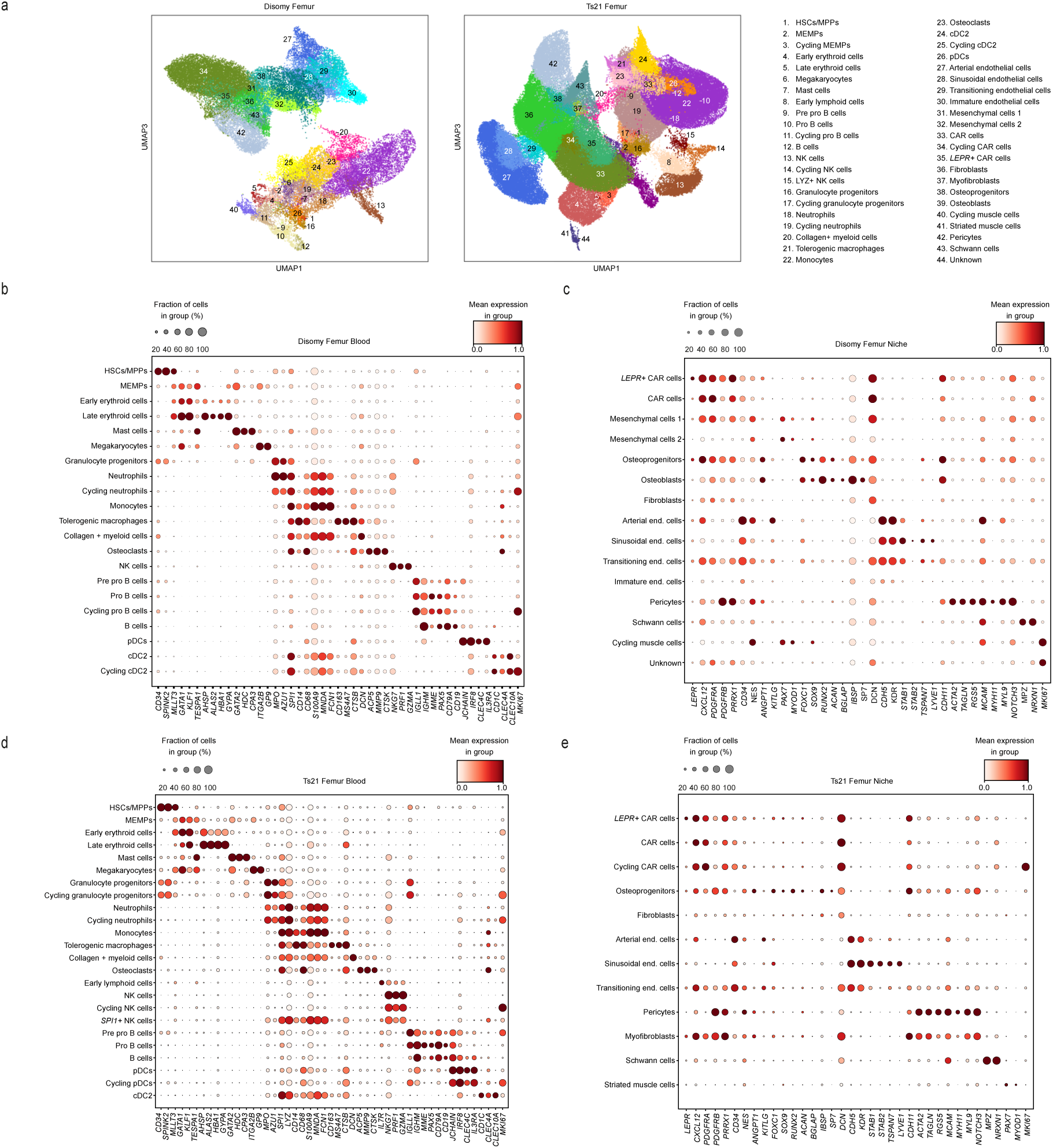
A single-cell atlas of disomic and Down’s Syndrome foetal bone marrow. a. UMAP visualisation of cells from disomic foetal BM (left; n=53,807, k=3) and Ts21 (right; n=162,775, k=12) coloured by cell type/state. **b-e,** Dot plot of the standardised gene expression of manually selected marker genes in the identified cell types. For each gene, the minimum value of its expression is subtracted and the result is divided by the maximum value of its expression. The dot size indicates the percentage of cells that express the gene of interest within each cell type. **b**, disomic BM haematopoietic cells. **c**, disomic BM niche cells. **d**, Ts21 BM haematopoietic cells. **e**, Ts21 liver BM cells; n refers to the total number of cells, k indicates the number of biologically independent samples.

**Supplementary Figure 5.**
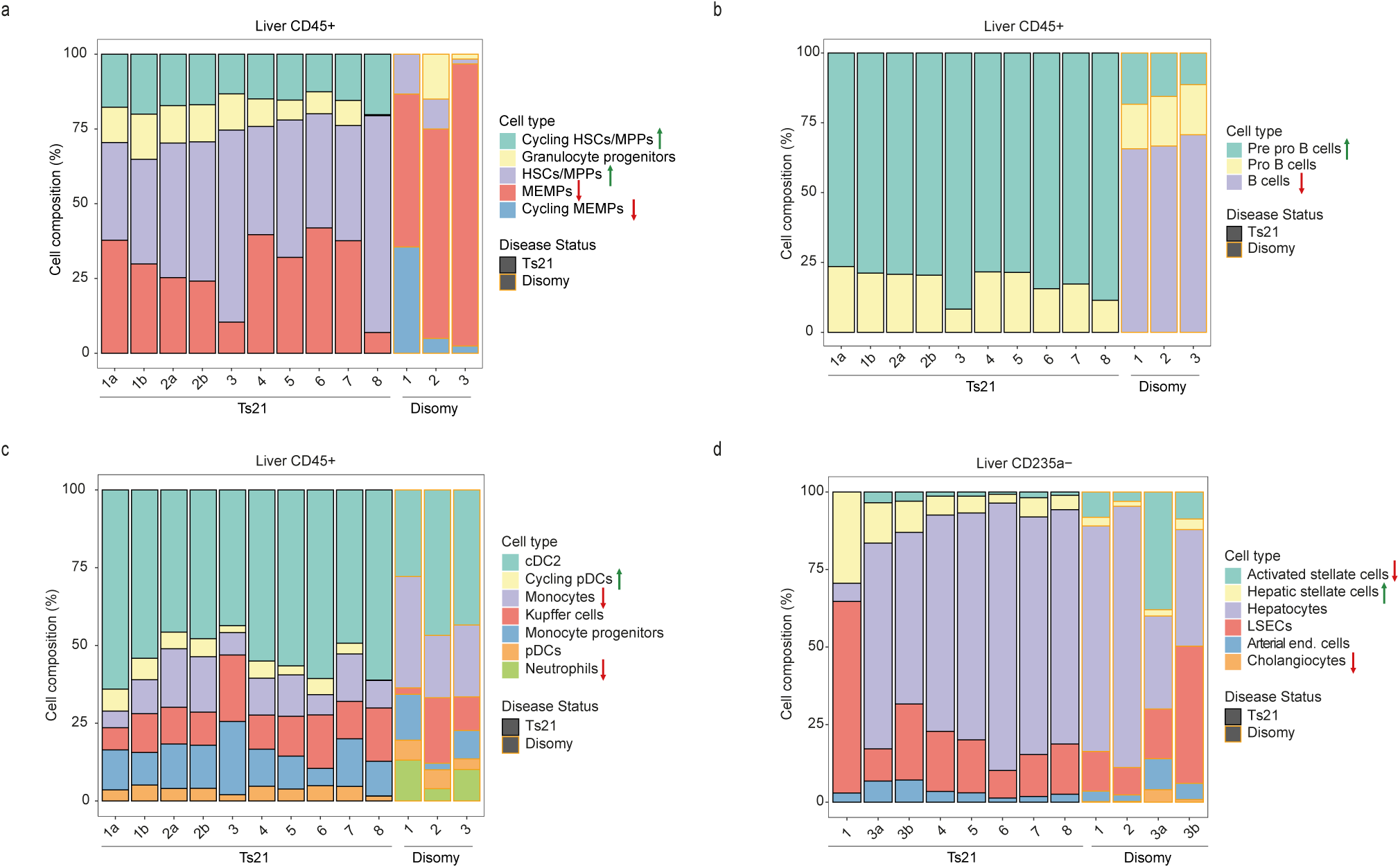
Relative abundances of blood and stroma cells in Ts21 and disomic liver. **a-d**. Stacked bar plots of relative abundances of different cell types/states in stage-matched Ts21 (black outline) and disomic (orange outline) foetal liver. Arrows at the side of the labels indicate a significant increase (↑) or decrease (↓) in the number of cells in Ts21 compared to disomic liver. Difference in cell proportion was tested by Wilcoxon Rank Sum test followed by Bonferroni correction. **a**, Haematopoietic progenitors subsets identified within CD45+ cells. **b**, B cells and B cell progenitors identified within CD45+ cells. **c**, Myeloid cells identified within CD45+ cells. **f**, Non-haematopoietic cells (niche) identified within CD235- depleted cells.

**Supplementary Figure 6.**
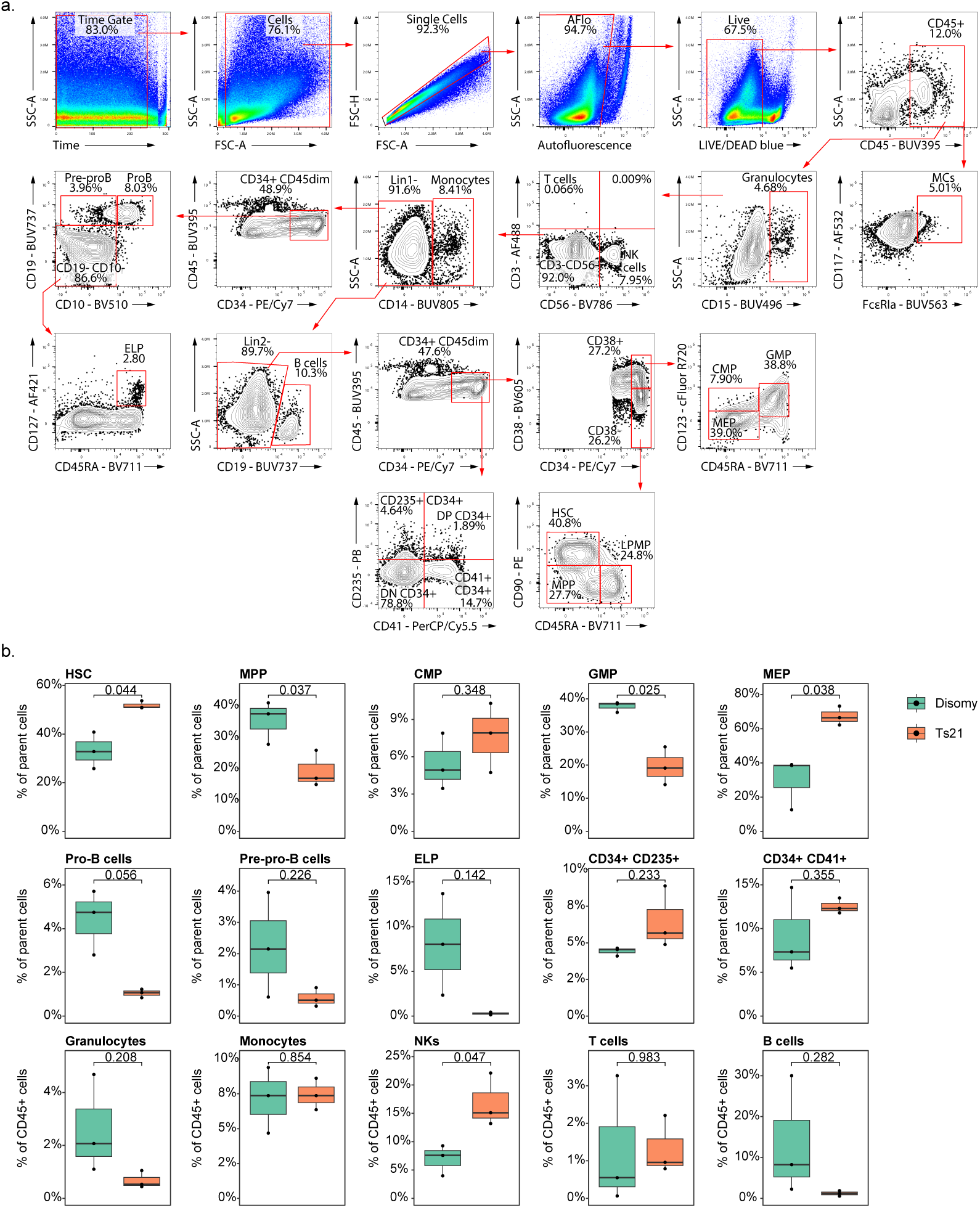
Sorting strategy and frequency of immunophenotypic blood cells. **a**, Representative gating strategy used to identify stem cell subsets, progenitors and mature cells in foetal liver. Gates are based on the disomic foetal liver sample. b, Frequency of progenitor and stem cell subsets in T21 and disomic foetal liver. The values on the boxplot indicate the results of a Welch-corrected t-test comparing the percentage of cells in disomic and T21 foetal livers.

**Supplementary Figure 7.**
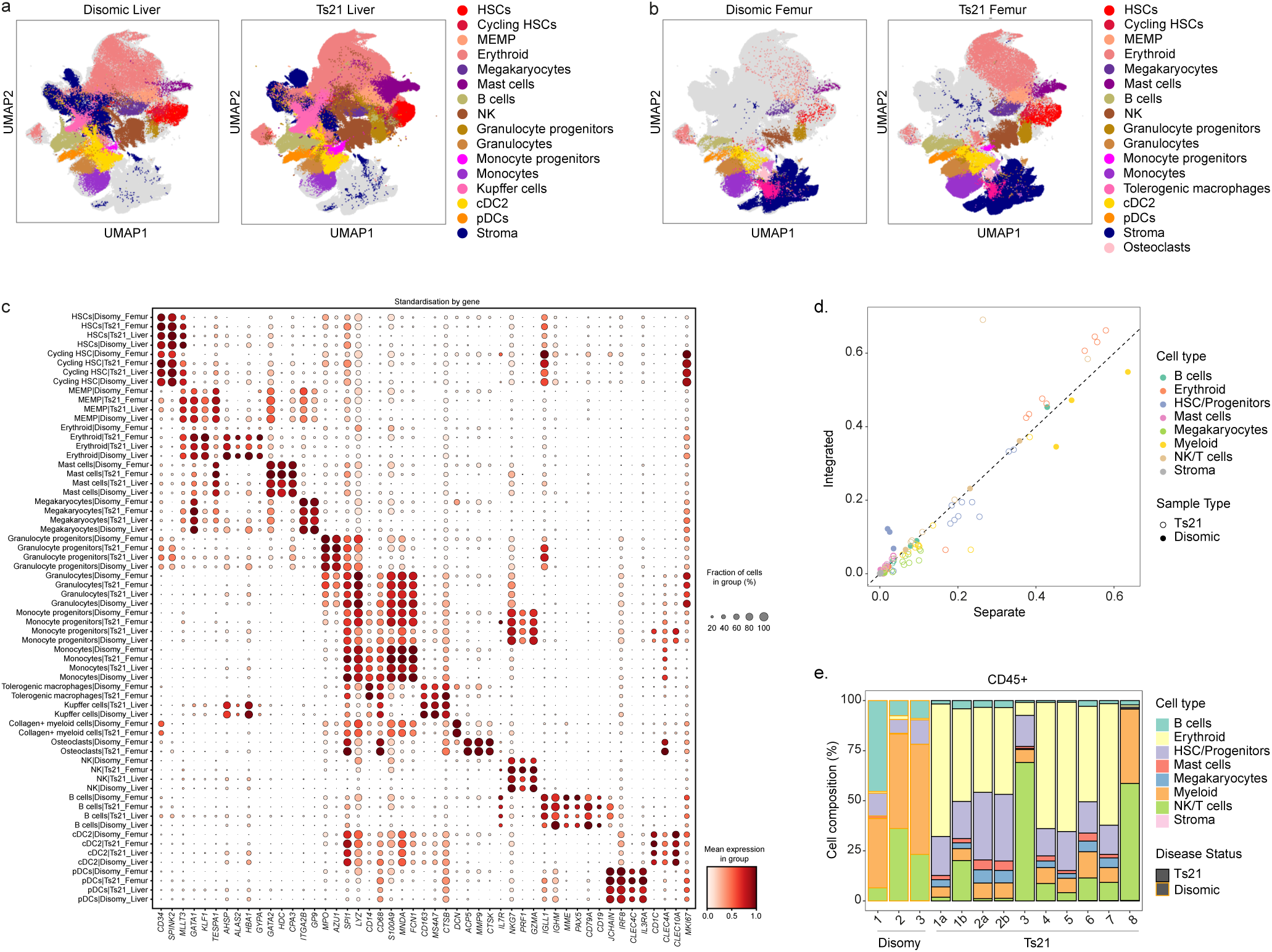
Integration of liver, femur, Ts21 and disomic cells using Harmony. 3D Uniform manifold approximation and projection (UMAP) visualisation of cells from (**a**) Ts21 liver (n=780,299, k=15), and disomic liver (n=110,671, k=3) and (**b**) Ts21 femur (n=162,775, k=12), and disomic femur (n=53,807, k=3) following integration of datasets using Harmony. Cells are coloured by broad cell type categories. n refers to the total number of cells, k indicates the number of biologically independent samples (foetuses). **c,** Gene expression dot plot of marker genes from Supp. Figure 3 and 4 across cell types identified from Harmony-based dataset integration. **d,** Using the Harmony-integrated cell type labels in stage-matched Ts21 and disomic foetuses, we compared cell type proportions between separate and integrated. Each point represents the estimated abundance in a different sample. **e,** A stacked bar plot of relative abundances of different broad cell types based on annotations after integration.

**Supplementary Figure 8.**
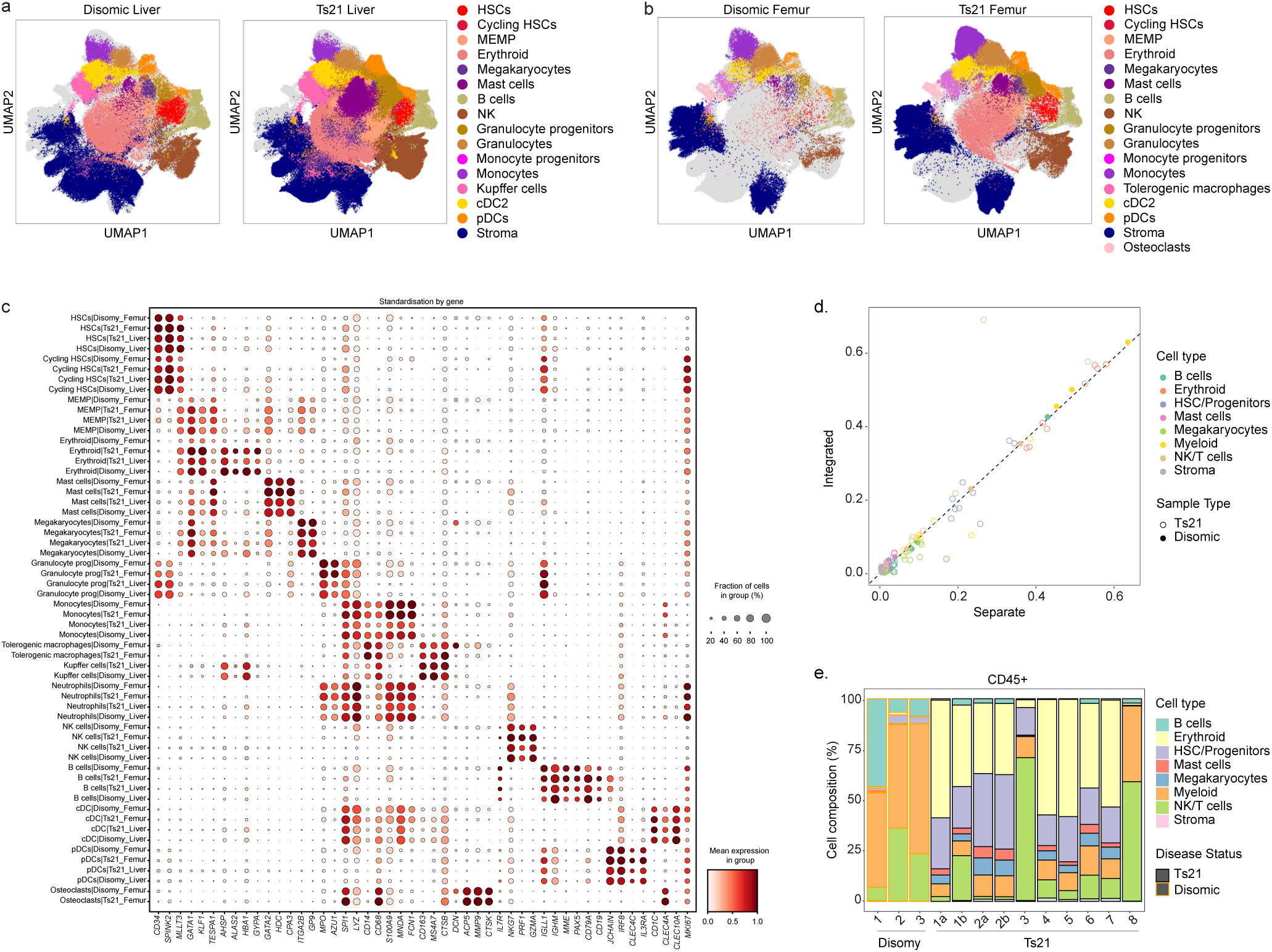
Integration of liver, femur, Ts21 and disomic cells using scVI tools. 3D Uniform manifold approximation and projection (UMAP) visualisation of cells from (**a**) Ts21 liver (n=780,299, k=15), and disomic liver (n=110,671, k=3) and (**b**) Ts21 femur (n=162,775, k=12), and disomic femur (n=53,807, k=3) following integration of datasets using Harmony. Cells are coloured by broad cell type categories. n refers to the total number of cells, k indicates the number of biologically independent samples (foetuses). **c.** Dot plot showing expression of marker genes from Supp. Figure 3 and 4 across cell types identified following dataset integration using scVI tools. **d.** Using the scVI-integrated cell type labels in stage-matched Ts21 and disomic foetuses, we compared cell type proportions between separate and integrated. Each point represents the estimated abundance in a different sample **e.** A stacked bar plot of relative abundances of different broad cell types based on annotations after integration.

**Supplementary Figure 9.**
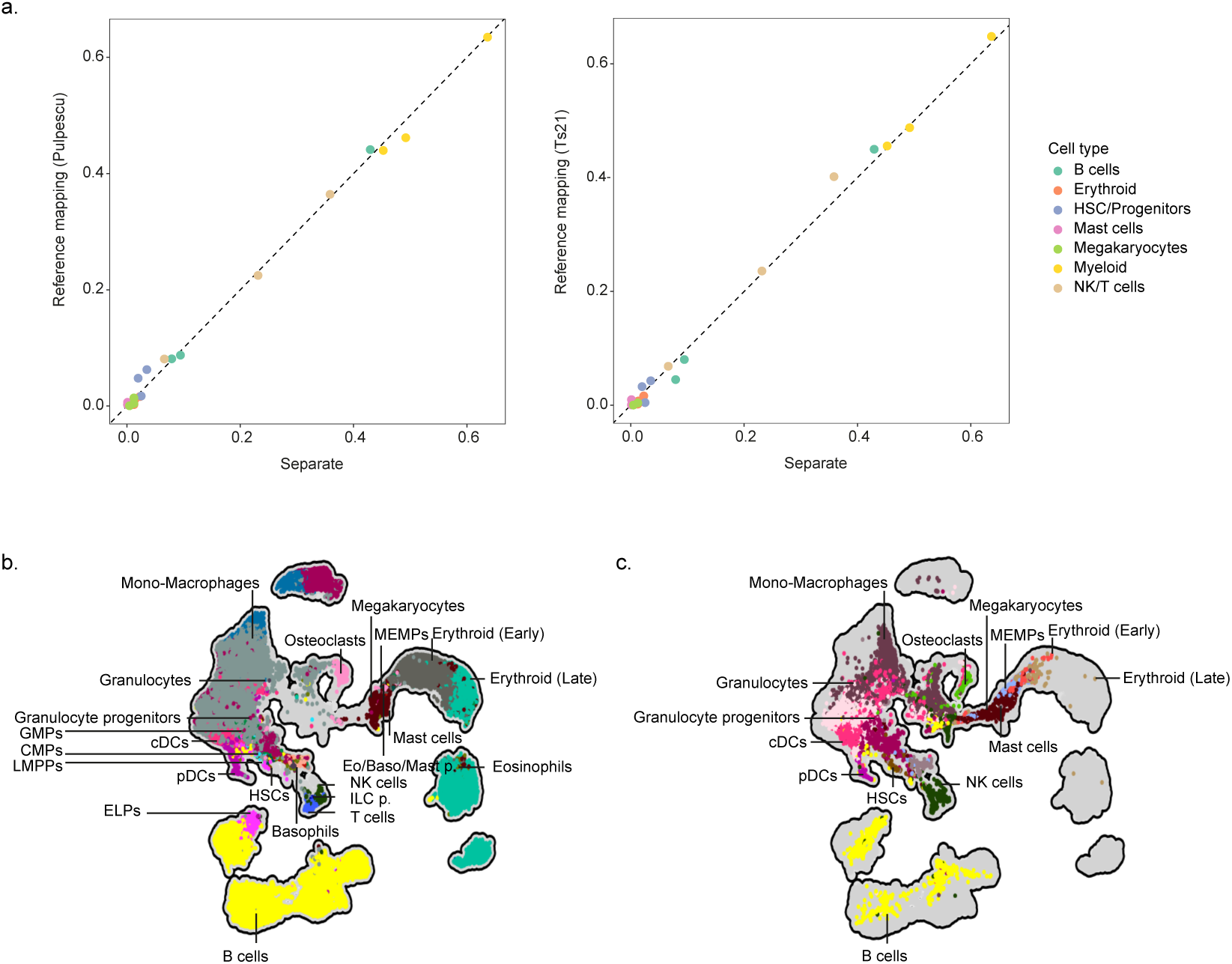
Integration of liver and disomic cells and label transfer. **a**, Comparison of cell type proportions for stage-matched disomic liver samples from the separate-dataset annotations to the reference query label transfer annotations using Popescu et al. (left) or the Ts21 dataset as a reference (right). **b**, Integration of Jardine et al. and Marderstein et al. cells from disomic BM using scVI. The Jardine et al. cells are shown on the left, and the Marderstein et al. cells are shown on the right. Cells are coloured by the annotated population from their original studies.

**Supplementary Figure 10.**
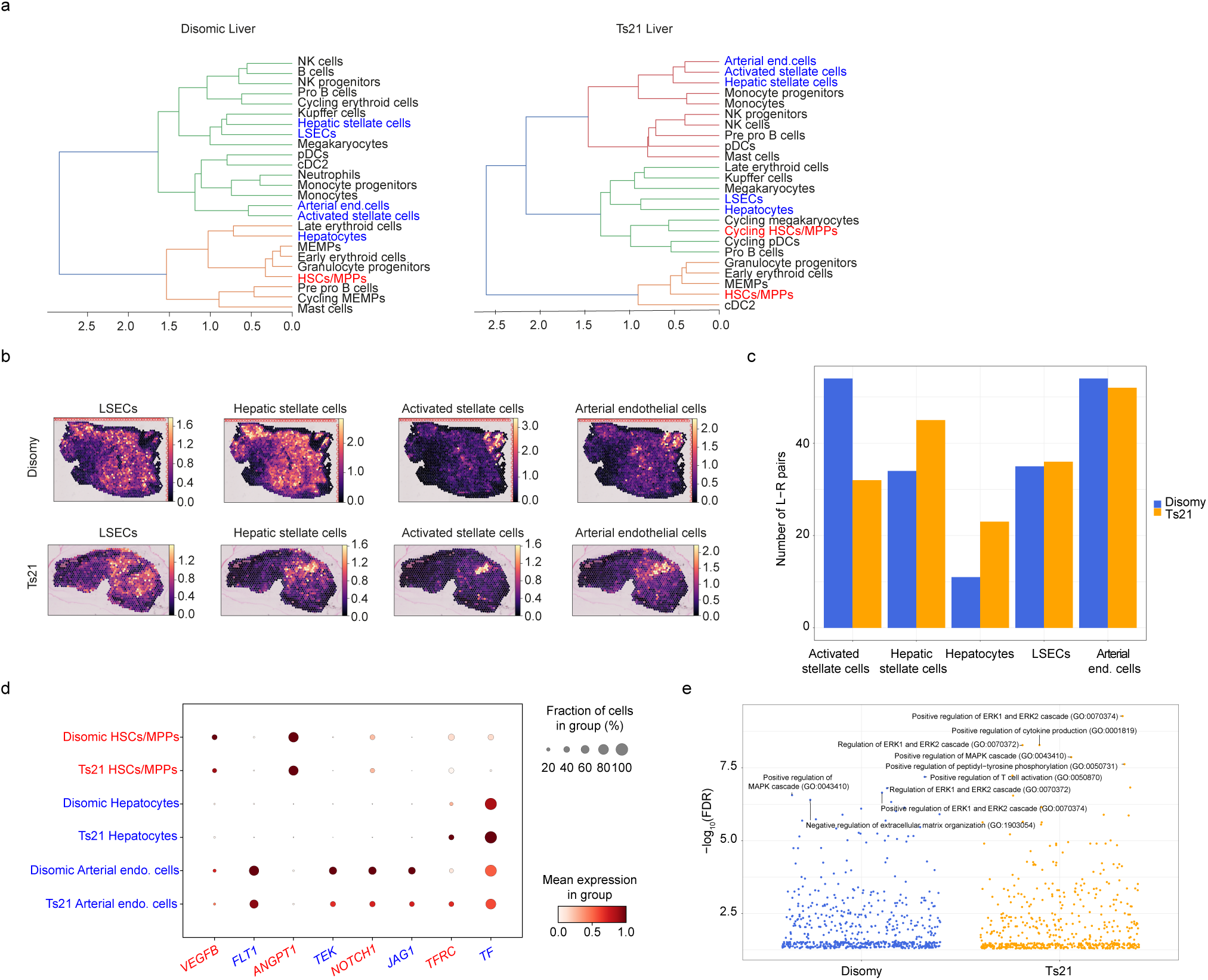
Colocalisation and L-R analysis from Visium data. **a,** Dendrogram from hierarchical clustering results based on correlation distance, with inter-cluster distance estimated by Ward variance minimization algorithm. HSCs/cycling HSCs are coloured in red, stromal cells are coloured in blue, all other blood cells are coloured in black. **b,** cell2location mapping of LSECs, hepatic stellate cells, activated stellate cells and arterial endothelial cells on two representative Visium sections from disomic foetal liver (top) and Ts21 liver (bottom). The colour of each spot indicates the cell type abundance estimated by cell2location (mean value). **c,** The number of L-Rs that HSC/MPPs use to interact with different niche cells. **d**, Dotplot of L-Rs expressed in HSC/MPPs, hepatocytes and arterial endothelial cells. Dot size represents the fraction of cells expressing the gene, and colour indicates the mean expression level, both computed within the cell type cluster. The genes coloured in red are L/Rs associated with HSCs/MPPs, and the genes coloured in blue are those associated with hepatocytes and arterial endothelial cells. **e,** Gene-set enrichment analysis of L-Rs expressed in HSC/MPPs used for their interaction with arterial endothelial cells. Top 5 terms in each group were labelled. The presented results include all sections from Ts21 (n=20 sections; 2 consecutive sections from 10 biological replicates) and disomic (n=10 sections; 2 consecutive sections from 5 biological replicates) tissues.

**Supplementary Figure 11.**
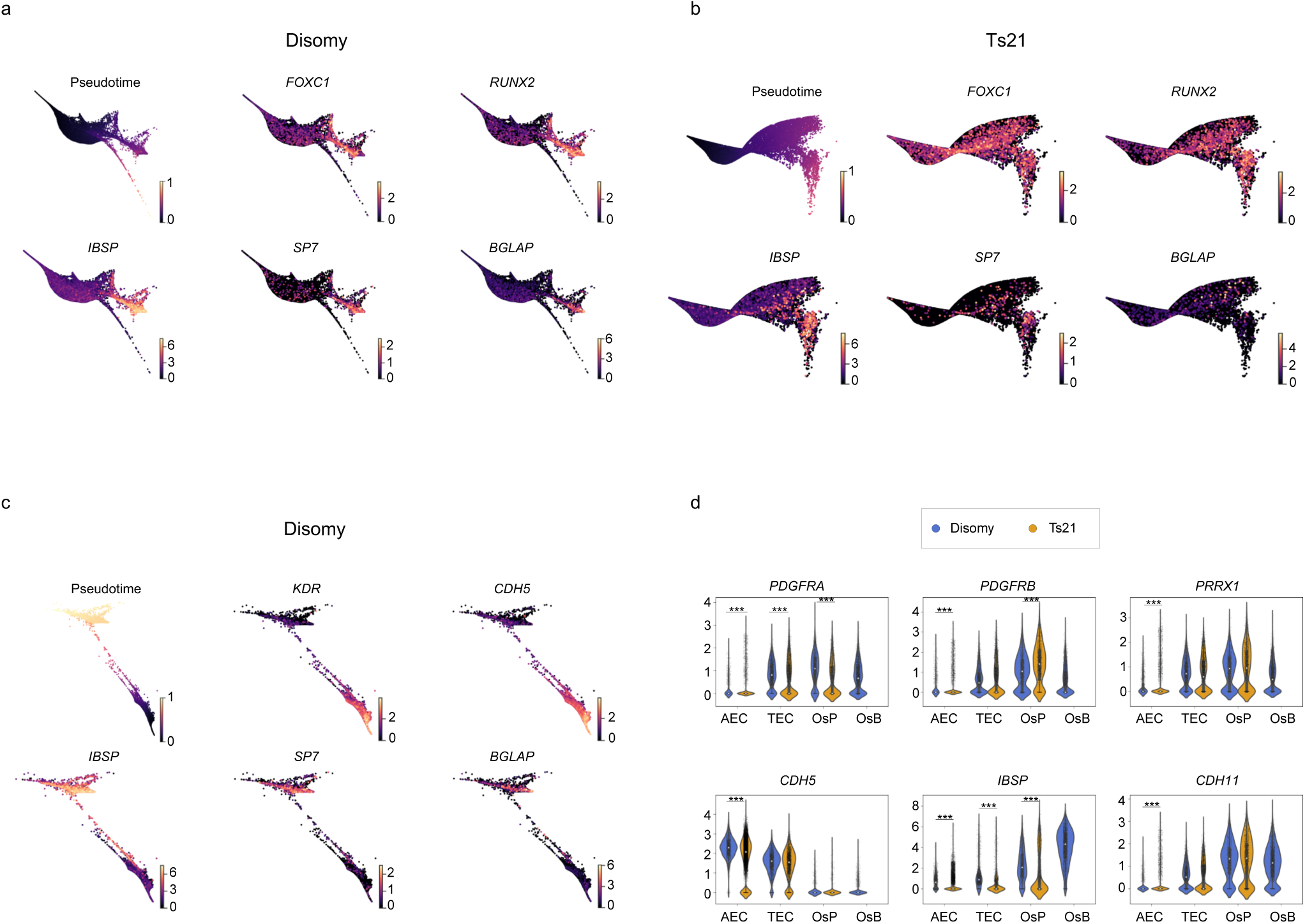
Inferring differentiation trajectory of osteoblasts. a-c, Pseudotime and gene expressions (normalised and log1p transformed) on the diffusion map in: **a,** disomic CAR cells, LepR+ CAR cells, osteoprogenitors and osteoblasts**; b,** Ts21 CAR cells, LepR+ CAR cells, osteoprogenitors**; c,** disomic arterial endothelial cells, transitioning endothelial cells and osteoblasts. **d,** Violin plots showing expression of genes in arterial endothelial cells (AEC), transitioning endothelial cells (TEC), osteoprogenitors (OsP), and osteoblasts (OsB). Normalised and log1p transformed gene expression was compared between disomic and Ts21; asterisk marks p-values < 0.001, computed by Wilcoxon rank sum test followed by Bonferroni correction.

**Supplementary Figure 12.**
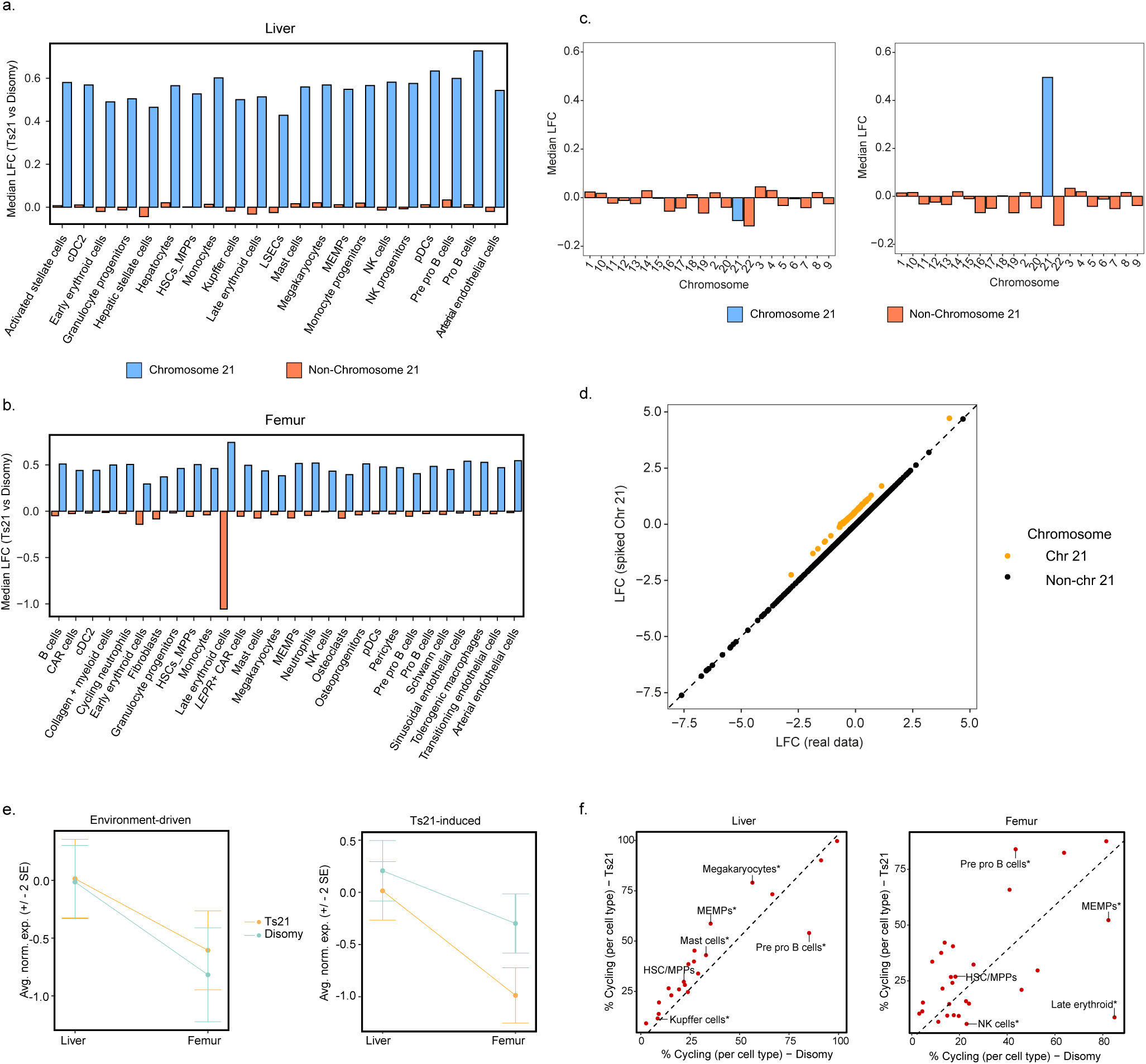
Impact of chromosome 21 on differential expression analysis. In liver **(a)** and femur **(b)**, differential expression analysis was performed between Ts21 and disomic cells using limma-voom and sample-level pseudobulks. The median log_2_-fold change (LFC) was calculated for chr21 genes (blue) and all other genes (Not chr21, red). **c,** Left, median log-fold changes for genes on each chromosome between fetuses #15633 and #15781. Right, median log-fold changes for genes on each chromosome between fetuses #15633 and #15781, after increasing mapped reads to genes on chromosome 21 by 50% in two #15781 pseudobulk samples. **d,** Comparison of log-fold changes for all genes between the two analyses. Orange points represent the genes on chromosome 21, which have increased expression within the two #15781 pseudobulk samples in the spiked chr21 analysis (y-axis). The log-fold changes of the underlying real data is along the x-axis. **e,** Average normalised gene expression of statistically significant *Environment-driven* DEGs and *Ts21- induced* HSC/MPPs DEGs within each dataset. Normalised expression was calculated by first transforming a gene’s expression to have mean = 0 and variance = 1, and then averaging transformed expression across all genes. Mean normalised expression across cells is shown, with vertical bars representing 2 standard errors around the mean. **f,** Comparison of the percentage of cells cycling within disomic samples (x-axis) and Ts21 samples (y-axis). Each dot represents a different cell type. Asterisk (*) signifies statistically significant difference in cycling of cells in Ts21 vs disomic by performing a Mann-Whitney U test on sample-level pseudobulk proportions. Dashed line represents *y = x*.

**Supplementary Figure 13.**
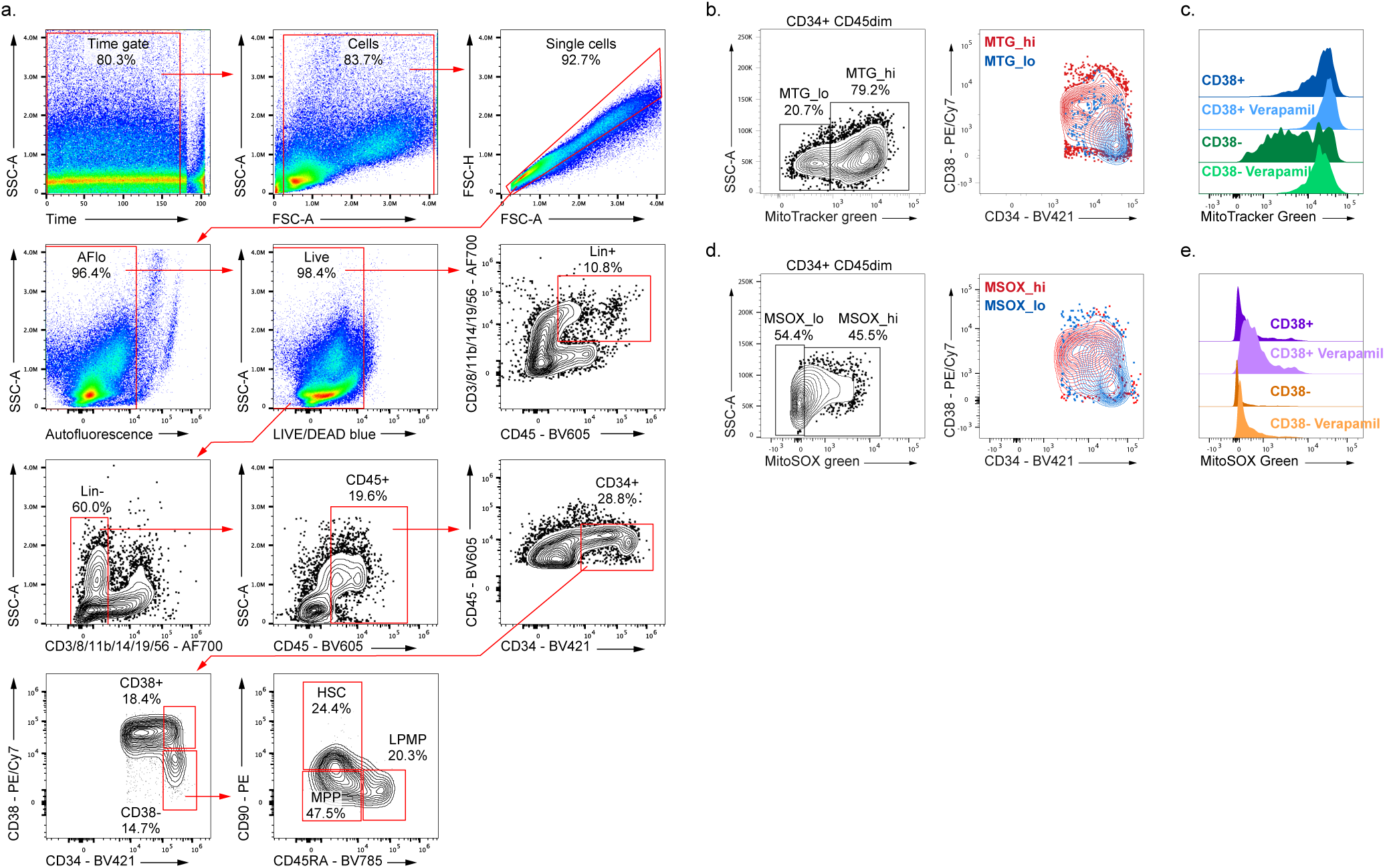
Sorting strategy and measuring mitochondrial mass and oxidative stress in haematopoietic cells. **a**, Representative gating strategy used for the mitochondrial probe experiments. Both MitoSOX and MitoTracker stainings shared the same antibody panel. **b**, Left: Representative gating strategy used to identify MTG_hi and MTG_lo populations. Right: MTG_hi and MTG_lo cells were plotted against CD34 and CD38, showing that MTG_lo cells are enriched in the CD34+ CD38- fraction. **c**, Density plot of the MitoTracker Green FM fluorescence in CD38+ and CD38- cells treated or not with Verapamil 50µM during the MitoTracker incubation. Cells were gated on Time gate/Cells/Single cells/AFlo/Live/Lin-/CD45+/CD34+. **d**, Left: Representative gating strategy used to identify MSOX_hi and MSOX_lo populations. Right: MSOX_hi and MSOX_lo cells were plotted against CD34 and CD38, showing that MSOX_lo cells are enriched in the CD34+ CD38- fraction. **e**, Density plot of the MitoSOX Green fluorescence in CD38+ and CD38- cells treated or not with Verapamil 50µM during the MitoSOX incubation. All cells were gated on Time gate/Cells/Single cells/AFlo/Live/Lin-/CD45+/CD34+.

**Supplementary Figure 14.**
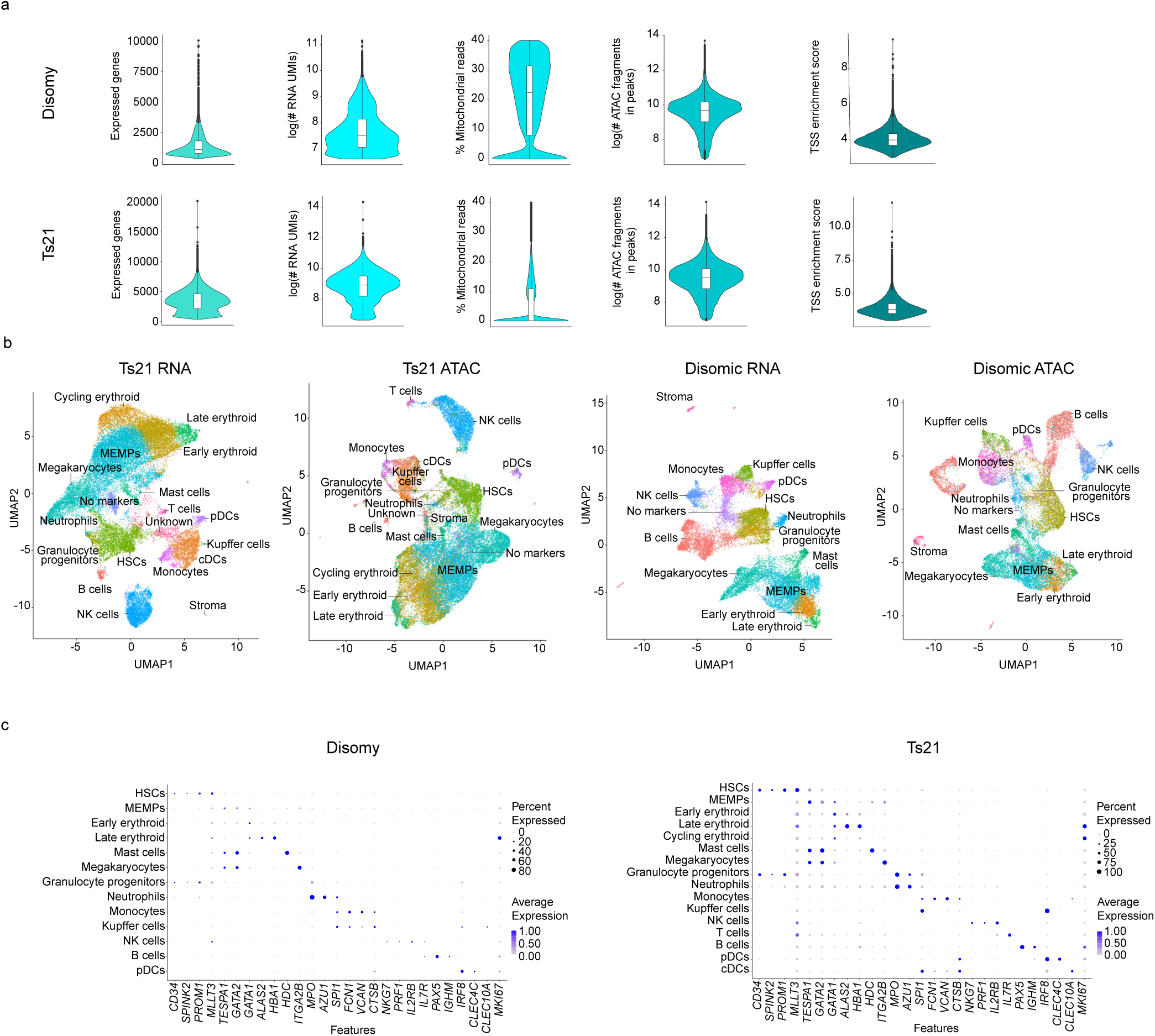
Multiome atlas. **a**, Violin and boxplots for quality control metrics from multiome data. The bottom, middle, and top lines of the boxplot reflect the 25th, 50th (median), and 75th TRS percentiles, and the boxplot whiskers reflect 1.5x the interquartile range. **b,** UMAP visualisation of snRNA-seq (left) and corresponding scATAC-seq (right) of CD45+ cells isolated from **b,** Ts21 foetal liver (n=35,633, k=3, median PCW=13) and disomic foetal liver (n=21,257, k=3, median PCW=13). n- number of cells; k- number of biological replicates. **c,** Dot plot of the standardised gene expression of manually selected marker genes in the identified cell types. For each gene, the minimum value of its expression is subtracted and the result is divided by the maximum value of its expression. The dot size indicates the percentage of cells that express the gene of interest within each cell type.

**Supplementary Figure 15.**
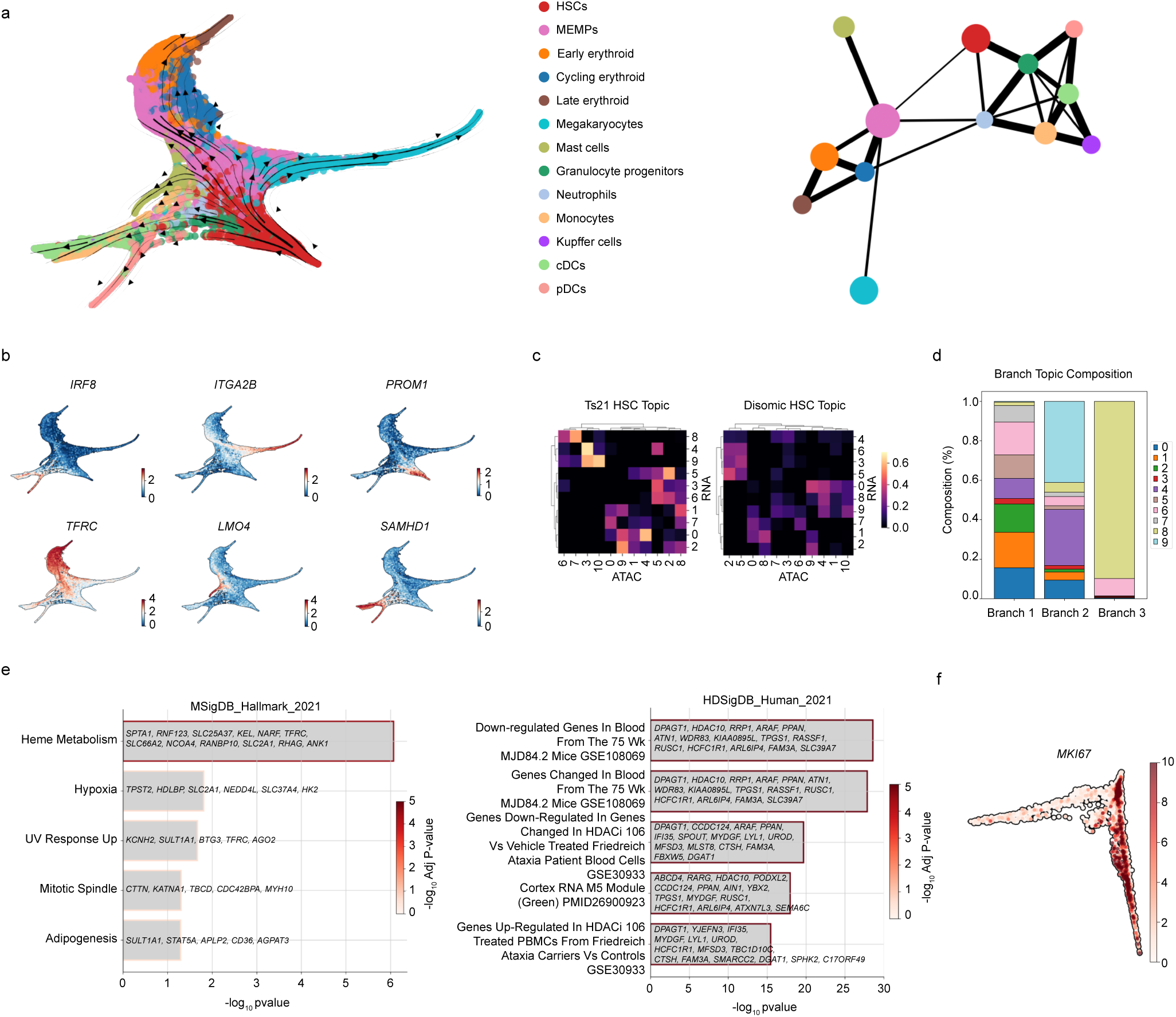
CellRank and MIRA differentiation analysis. **a,** Left, force directed layout (FLE)(ForceAtlas2) of cells belonging to the myeloid lineage. Harmony batch corrected and PAGA initialised. HSC cluster set as the root for pseudotime calculation. Black arrows indicate the projected transition matrix (CellRank). Right, PAGA representation. **b**, Five terminal states were automatically identified using CellRank. Normalised expression of genes with highest correlation with each terminal state plotted in FLE space (CellRank lineage drivers) (IRF8:pDCs, ITGA2B:Megakaryocytes, TFRC:Late erythroid, LMO4:Mast cells, SAMHD1: monocytes). PROM1 was added to illustrate the location of HSCs. **c**, MIRA topic analysis performed on data including both disomic and Ts21 HSCs. Topic analysis was performed separately on RNA and ATAC raw data. Cross-correlation matrix between RNA topics and ATAC topics. Left, Ts21 HSCs. Right, disomic HSCs. **d**, Each cell is associated with a single RNA topic according to the topic with the highest cumulative expression in that cell. Bar graph illustrating the composition of topics each MIRA identified branch, as measured by topics assigned to each cell of that branch. RNA topic 9 has the highest frequency in branch 2 and topic 8 has the highest frequency in branch 3. **e**, Left, gene set enrichment analysis (GSEA) of branch 2 specific topic 9 from querying “MSigDB_Hallmark_2020” (enrichr). GSEA analysis of branch 3 specific topic 8 from querying “HDSigDB_Human_2021” (enrichr). **f**. UMAP plot representing the joint RNA and ATAC MIRA latent space of HSCs. Expression of cell-cycling gene MKI67 (raw counts).

**Supplementary Figure 16.**
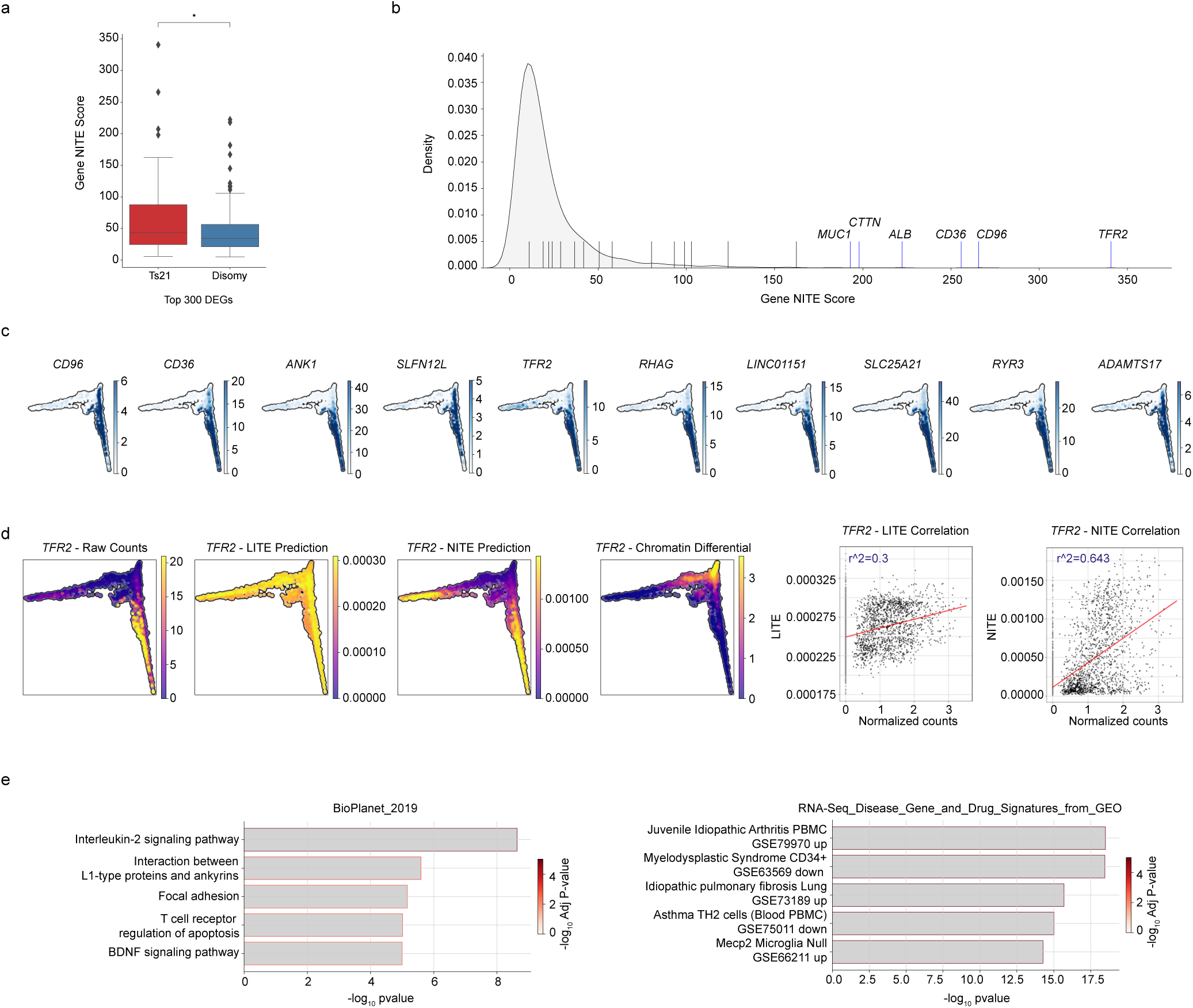
Trans-regulated genes in HSC multiome. **a,** The cumulative gene NITE score (MIRA) across all HSCs separated by differentially expressed genes (DEGs) (pseudobulk DESeq2) between disomic and Ts21. Significance performed using a Wilcoxon rank sum with Benjamini-Hochberg correction. **b**, Distribution of the cumulative gene NITE score across all modelled genes (n=4,446). Black, top genes regulated by branch 2 transcription factors. Blue, top genes with highest NITE score. **c**, UMAP plots (MIRA’s joint RNA and ATAC topic model latent representation) indicating the raw expression of genes with the highest cumulative NITE score. **d**, Representation of the gene, *TFR2*, with the highest NITE score. UMAP plots from left to right illustrate raw expression, LITE modelling prediction, NITE modelling prediction, and the chromatin differential (difference between LITE and NITE). Scatter plots (right) indicate the correlation between regulatory potential modelling prediction and the normalised expression of the gene. The NITE model that incorporates both cis and trans accessibility greater than doubles the predictive efficiency of *TFR2* expression (LITE: *R*^2^=0.3, NITE: *R*^2^=0.64). e, Gene set enrichment analysis (GSEA) (enrichr) results from the top 500 genes according to the cumulative NITE score. Left, GSEA results from querying “BioPlanet_2019.” Right, GSEA results from querying “RNA- Seq_Disease_Gene_and_Drug_Signatures_from_GEO.” Colouring indicates adjusted p-value (-log 10).

**Supplementary Figure 17.**
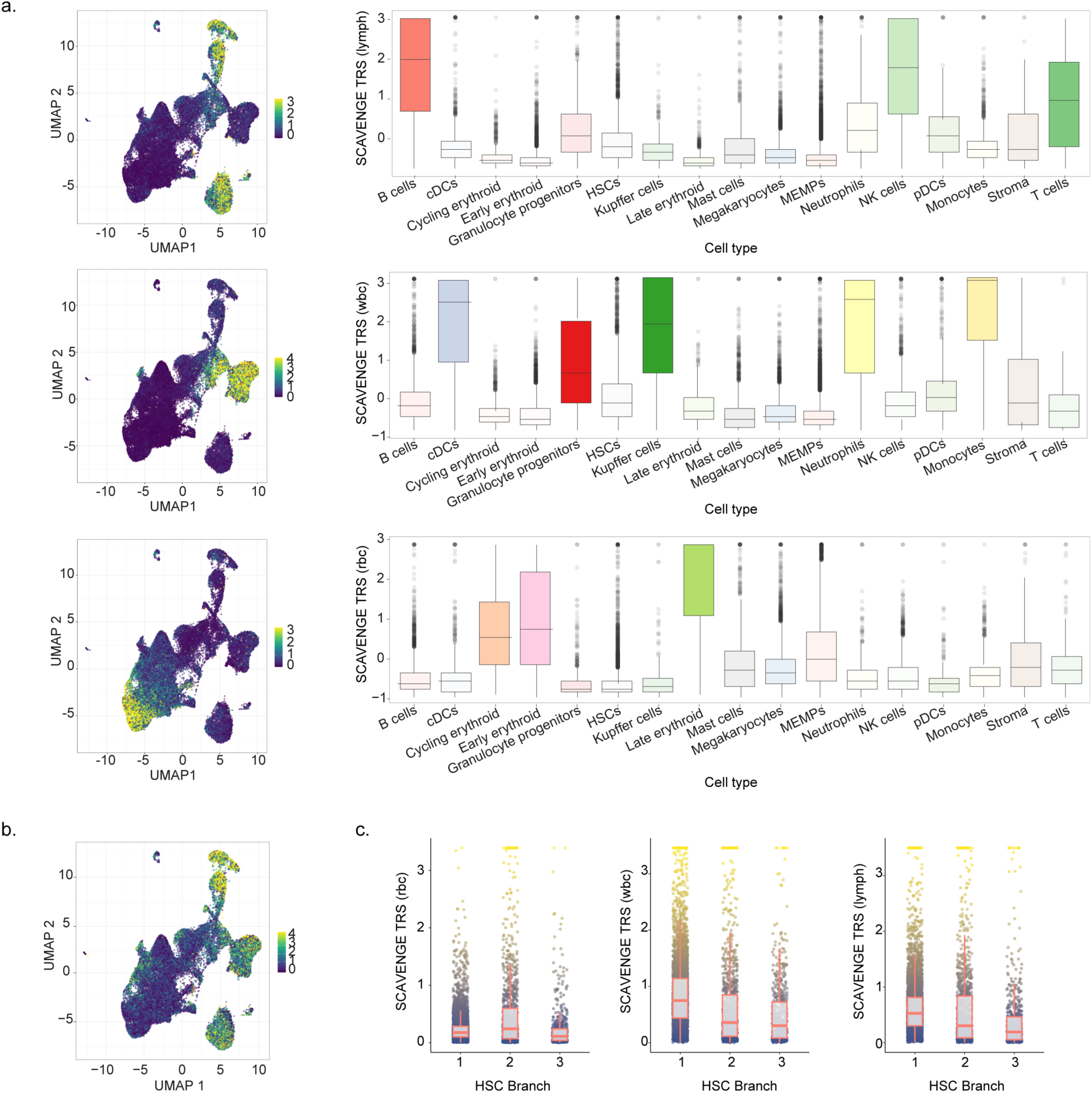
SCAVENGE application to scATAC. **a,** Left, UMAP of scATAC data in the combined data (disomic and Ts21). Each point is a different cell, coloured by SCAVENGE TRS for lymphocyte counts (top; lymph), white blood cell counts (middle; wbc), and red blood cell counts (bottom; rbc). Right, boxplots display SCAVENGE TRS across cell populations. The bottom, middle, and top lines of the boxplot reflect the 25th, 50th (median), and 75th TRS percentiles, and the boxplot whiskers reflect 1.5x the interquartile range. **b**, SCAVENGE trait relevance scores (TRS) for 3 blood cell traits (red blood cell counts (rbc), white blood cell counts (wbc), and lymphocyte counts (lymph)) across the 3 identified HSC branches. Each point is a different cell. The bottom, middle, and top lines of the boxplot reflect the 25th, 50th (median), and 75th TRS percentiles, and the boxplot whiskers reflect 1.5x the interquartile range.

**Supplementary Figure 18.**
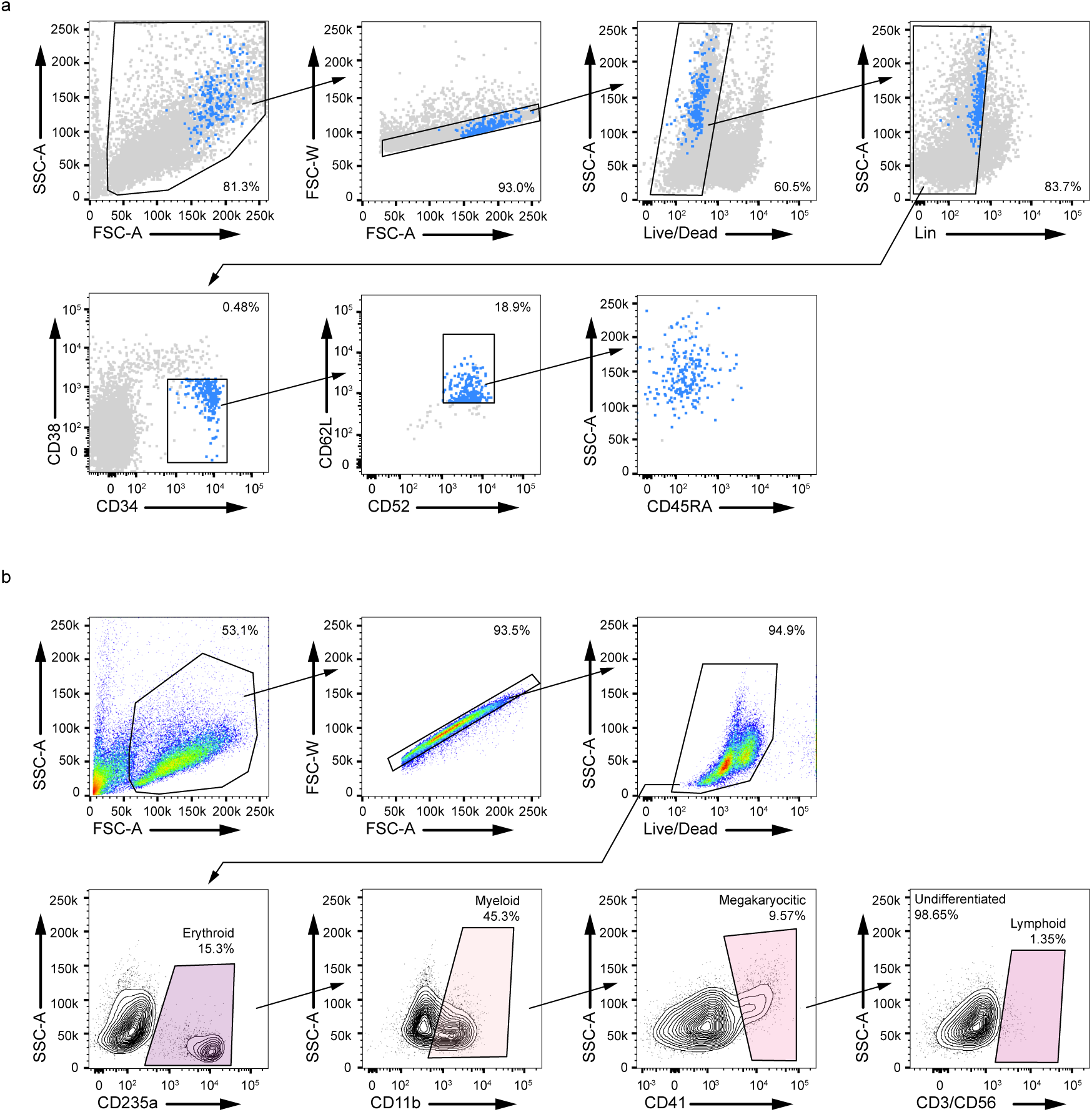
Single-cell CFU assay. **a**, Representative FACS sorting panel and gating strategy for the isolation of HSC/MPPs (CD34+CD38-CD52+CD62+Lin-). All HSC/MPPs were CD45RA-. **b,** Representative FACS plot showing the lineage composition of a quadri-lineage colony.

**Supplementary Figure 19.**
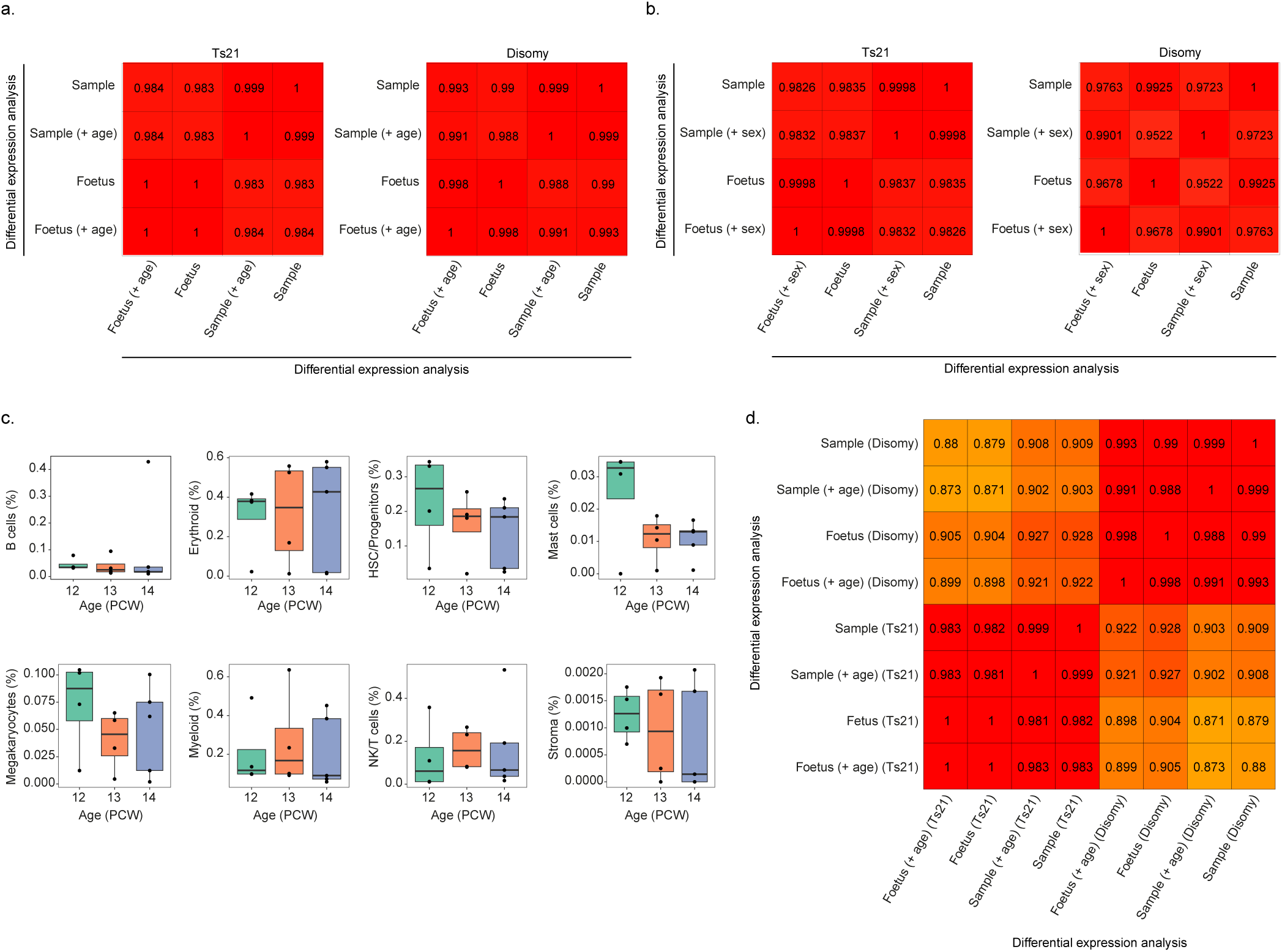
Evaluating the robustness of differential expression results. Pearson’s correlation coefficient between Liver vs Femur log-fold changes across all genes for the four analysis types, using age (**a**) or sex (**b**) as covariates. **c**, The relationship between post-conceptual weeks and cell type abundance (estimated as proportions) for the 8 broad cell type groups. **d**, Pearson’s correlation coefficient between Liver vs Femur log-fold changes across all genes for the four analysis types in the two datasets (8 distinct analyses in all).

**Supplementary Figure 20.**
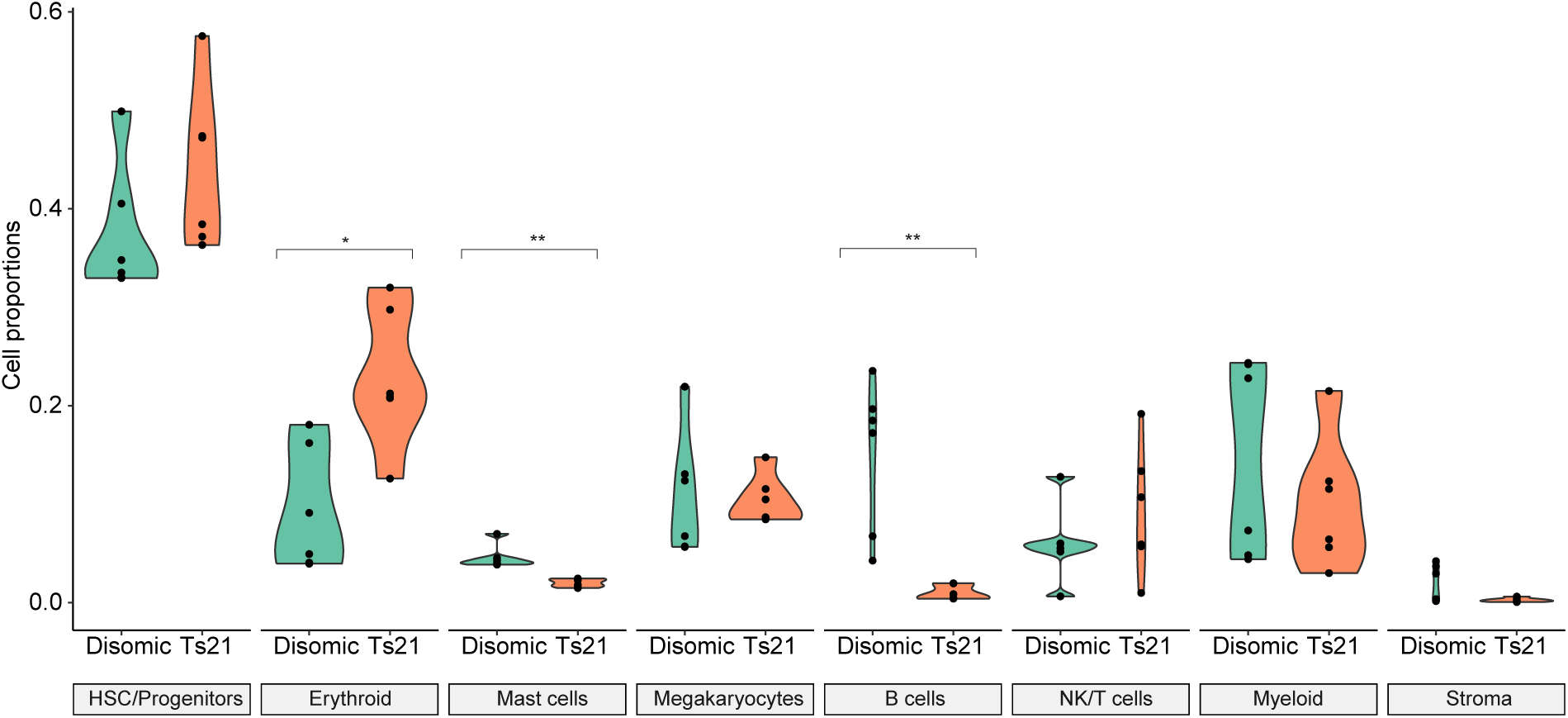
Comparison of cell-type frequencies from 10X multiome versus scRNA-seq. Mann-Whitney U test of differences in cell-type proportions between Ts21 and disomic samples. Significance was determined after FDR correction across cell types. P<0.05(*); P<0.01(**).

**Supplementary Table 1 - disomic foetal cells quality control**

**Supplementary Table 2 - Marker genes for each cell population.** Column descriptions: type = DownSyndrome or Disomic, organ = Femur or Liver, numerical_labels = cluster number, cell_type = labelled cell type population for the cluster, Rank = gene rank for the cell type, logfoldchanges = log-fold change of differential expression between cells in cluster and all other cells, pvals = p-values, pvals_adj = FDR-adjusted p-values.

**Supplementary Table 3 - Ts21 foetal cells quality control Supplementary Table 4 - Cell types per replicate, Ts21 liver scRNA-seq**

**Supplementary Table 5 - Cell types per replicate, disomic liver scRNA-seq Supplementary Table 6 - Cell types per replicate, Ts21 femur scRNA-seq Supplementary Table 7 - Cell types per replicate, disomic femur scRNA-seq Supplementary Table 8 - Cell type abundance comparison.** Median abundance in Ts21 samples and disomic samples, Mann-Whitney U test p-values and their FDR corrected values. **Supplementary Table 9 - 10X Visium summary**

**Supplementary Table 10 - Multiome summary**

**Supplementary Table 11 - Key characteristics of RNA and ATAC topics.** Top 200 genes and enriched TFs relevant to those genes in RNA topics, and top 10% peaks and enriched TFs relevant to ATAC topics. For calculating enriched TFs in RNA topics, we used regulatory potential modelling to calculate association scores between each TF and gene. A Wilcoxon test is performed to test for differences in TF association scores between the top genes for each topic and all other genes, thus identifying which TFs are “regulating” that topic. For calculating enriched TFs in ATAC topics, we tested the enrichment of TF motifs in the top peaks for the topic. Only TFs with P-value < 0.05 are shown. Column descriptions: datatype=RNA or ATAC; feature = Top 200 Genes, Top 10% Peaks, TF enrichment of Top 200 Genes, or TF enrichment of Top 10% Peaks; variable = topic number; Rank = rank of value in feature set; value = either TF label, gene, or peak; pval = p-value for TF enrichment (otherwise NA).

**Supplementary Table 12 - TF regulator results from MIRA**. Source (src) specifies which branches are the TF regulator results related to. The columns id, name, parsed_name, pval, and test_statistic detail motif identifier, TF name, a computationally parsed TF identifier, enrichment significance (nominal P-value), and enrichment test statistic from MIRA. **Supplementary Table 13 - Peak-gene links tested by SCENT.** Gene_peak, gene, and peak describe the peak and gene tested. chr = chromosome, snp_pos = basepair position, rsid, and pip = finemap posterior inclusion probability are filled if the peak overlaps with a fine-mapped GWAS SNP (PIP > 0.2) for red blood cell counts. beta_H, se_h, z_H, p_H, boot_basic_p, fdr_H contain the estimated effect, standard error, z-score, non-bootstrapped P-value, bootstrapped P-value, and Benjamini-Hochberg corrected P-value of chromatin accessibility for gene expression within disomic HSCs. *_t21 contains the same information for Ts21 HSCs, *_t21_dn contains the same information for Ts21 HSCs after downsampling to the disomic dataset size, and *_int contains the same information for the interaction term from the SCENT interaction model. Prefixes logFC, AveExpr, t, P.value, adj.P.Value, and B (which respectively represent log2 fold change, average expression across all samples in log2 counts per million, test statistic, nominal P-value, Benjamini-Hochberg corrected P-value, and log-odds of differential expression) are limma outputs from differential analyses of Ts21. The suffixes smATAC, smRNA, and lgRNA represent the limma analysis for multiome ATAC, multiome RNA, and large scRNA HSCs, where logFC > 0 is an increase in Ts21. The differential analyses were performed using pseudobulks.

**Supplementary Table 14 - Antibodies used in the study**

**Supplementary Table 15 - Differential expression between Ts21 and disomic cells.** Column descriptions: organ = Femur or Liver, cell = cell type population, names = gene name, logFC = log-fold change of differential expression between Ts21 and disomic samples, AveExpr = average expression in all samples, t = test statistic, P.Value = p-values, adj.P.Val = FDR-adjusted p-values

**Supplementary Table 16 - Cell barcode to cell type mapping.** Column descriptions: cell = barcode,patient = fetus identifier, sample = sample identifier, sorting = sorting strategy for the sample, type = DownSyndrome or Disomic, organ = Femur or Liver, numerical_labels = cluster number, cell_type = labelled cell type population for the cluster, cell_type_groups = mapping

